# Deciphering the Replication-Division Coordination in *E. coli* : A Unified Mathematical Framework for Systematic Model Comparison

**DOI:** 10.1101/2025.07.25.666816

**Authors:** Alexandre Perrin, Marie Doumic, Meriem El Karoui, Sylvie Méléard

**Affiliations:** CMAP, Ecole polytechnique, CNRS, Institut polytechnique de Paris, Inria, route de Saclay, 91128 Palaiseau Cedex-France; LBPA UMR8113, CNRS, Ecole Normale Paris Saclay, 91190, Gif-sur-Yvette, France

## Abstract

Despite extensive research, the quantitative principles that govern the coordination between DNA replication and cell division in bacteria remain debated. Multiple theoretical models have been proposed, some postulating that a single regulatory process is sufficient to ensure replication–division coordination, while others argue that two concurrent processes are required for robust control. In this work, we develop a unifying mathematical framework within which models can be consistently formulated, qualitatively analysed and quantitatively compared. This framework also allows us to propose a new double-process model. Through theoretical analysis, we establish the necessary and sufficient conditions under which single-process models can reproduce physiological cell behaviours. Beyond the correlation-based analyses extensively used to date, we further demonstrate within a comprehensive statistical framework that double-process models more accurately recapitulate experimental data across all growth conditions. Specifically, the new model we propose robustly captures the replication-division coordination in every growth regime, thereby providing a foundation for future mechanistic studies.

**Author Summary:** How bacteria regulate their cell size and ensure complete chromosome replication before division remains only partly understood, posing a fundamental question. A key challenge is understanding how bacterial cells reliably distribute their genetic material to daughter cells while managing randomness in division timing, especially since DNA replication is often initiated in earlier generations.

Over the past decade, several models have been developed to describe the coordination of DNA replication and cell division in *Escherichia coli*, combining different mechanisms and levels of complexity. These models have mainly been evaluated through correlation analysis between cell cycle parameters, such as the size at which bacteria divide or initiate DNA replication. However, such analyses depend on strong underlying assumptions that are not always met by the experimental data. Here, we present a general framework that systematically integrates multiple models, allowing both qualitative analysis and quantitative comparison without relying on correlations. Our work concludes that under all growth conditions, *E. coli* cells most likely divide only after meeting two combined requirements: adding a critical size since birth and a critical size since the initiation of DNA replication.

## Introduction

The regulation of the cell cycle in *Escherichia coli* has been investigated for more than half a century to understand how bacteria coordinate growth, chromosome replication, and division. These processes require the control of several quantities such as the cell size at division and the timing of the initiation of DNA replication, to ensure equal partitioning of genetic material between daughter cells. How these processes are integrated remains a central question in quantitative bacterial physiology [1, 2, 3]. Advanced microscopy techniques have enabled observations at the level of individual cells, revealing significant variability in both division size and inter-generation time between isogenic individuals. This highlights that control mechanisms that maintain cell homeostasis include stochastic variables [4, 5, 6, 7]. This stochasticity has challenged the deterministic approaches historically used to describe cell division in *E. coli* [8], necessitating alternative frameworks that better capture its regulation. As a result, cell-cycle dynamics in *E. coli* have been formalised using stochastic modelling frameworks, where each division event depends on the state of the cell in the previous cycle and on a control mechanism. This control is modelled with the help of independent identically distributed random variables. Analysis of experimental data has led to the identification of the so-called division adder model. In this model, the control variable is neither the cell size at division (sizer model) nor the inter-division time (timer model), but the volume added between birth and division [9, 10, 11, 12].

In parallel to these developments, the control of chromosome replication initiation has been extensively studied. As initially predicted by [13], and later confirmed experimentally, replication initiation events are tightly linked to the cell volume per origin of replication [14, 15, 5, 16]. Similar to the modelling of the division cycle, replication initiation cycles have been described using stochastic modelling, where the key variable is the volume per origin of replication at the time of initiation. Hence, an initiation adder model has been proposed, which assumes that the variable controlling the replication initiation event is the volume added per replication origin between successive initiation events [17, 18, 19].

A complete explanation of the bacterial cell cycle raises an additional challenge, which is the quantitative understanding of how division and replication initiation cycles coordinate, given that cells must ensure an exact partitioning of their genetic material to their progeny. This problem is especially intricate in bacteria such as *E. coli*, where replication cycles may overlap. Indeed, whereas under slow growth conditions, the cell replicates its chromosome entirely between birth and division, under faster growth conditions, as it takes more time to replicate a full chromosome than the duration of the cell cycle, cells are born with a partially replicated chromosome (Figure 1B). To describe this coordination, two parameters were introduced: the C period (i.e the time between replication initiation and termination) and the D-period (time between termination and the division separating the resulting sister chromosomes) by [8]. In fast growth conditions, the C+D periods, i.e. the time between replication initiation and the division that separates the resulting sister chromosomes, overlap (see Figure 1). While numerous molecular studies have provided mechanistic insights into this coordination [20, 21, 22, 23], various theoretical models have been developed over the past decade to account for the coordination of the division and replication cycles. The objective of these models is to identify the key independent random variables governing this coordination. Since most of these models assume that replication initiation cycles are governed solely by an initiation adder, the main question has been to determine the extent to which replication controls division, if at all. Among these models, two major categories can be distinguished. The first category of models, which we refer to as “single-process” models, supposes that division is set by a single process. Among this class of models, some models assume division to be independent of replication [24, 18, 25] and some assume replication to be the only factor that controls division [17, 19, 26]. The second category of models, which we call “double-process” models, assumes that division is triggered by the slowest of two independent processes, one linked to replication and another that is replication-independent [27, 28, 29, 30, 31].

**Figure 1.**
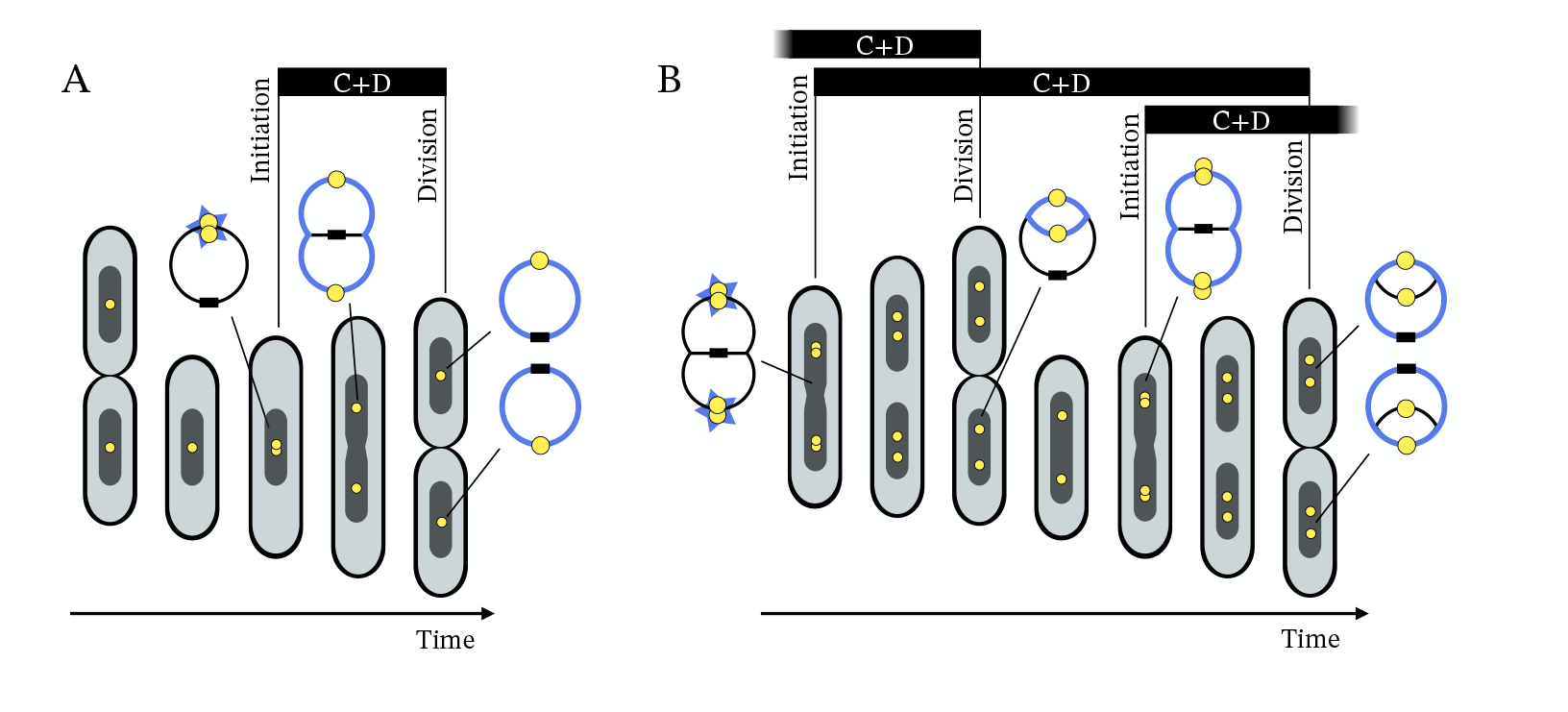
The replication and division cycles in *E. coli*. (A) In slow growth conditions, the replication of the chromosome is framed between the birth and the division. The period spanning from the initiation of replication to the division is called the C+D period. (B) In faster growth conditions, the initiation of replication may occur in previous generations, leading to overlaps of C+D periods. The yellow circles depict origins of replication, and sister chromosomes are colored in blue.

Based on correlation analysis, several studies have tested and compared replication-division coordination models in unperturbed cells [27, 28, 18, 19, 32, 29, 31]. For single-process models, which assume that only one variable controls division, the validation criterion consists of observing no correlation with the other variables [18, 19, 32]. This has been done systematically in [32] using correlation matrices. However, the validity of this criterion relies on the implicit assumption that all variables are normally distributed, a premise that has not been tested so far. A more in-depth analysis, using correlation-based conditional independence tests, showed that in intermediate and fast growth conditions, single-process models may not be able to explain the experimental data [31], suggesting that it is necessary to include double-process models in any comparison framework. Nonetheless, this analysis only accommodated models that rely on a linear combination of cell variables, thereby excluding the original formulations of the double-process models [27, 28, 29]. Additionally, double-process models include latent variables that cannot be experimentally observed. To overcome this issue, the validity criterion used so far has been the adequation between experimental and predicted correlations for the observable variables [27, 28, 29]. However, a limit of this approach is that the predicted correlations are computed using linear approximations of these inherently nonlinear models. More broadly, correlations only capture part of the structure of the data and provide limited information when the underlying variables are not normally distributed or exhibit nonlinear relationships. This highlights the need for a comparison framework that circumvents these limitations.

The diversity of methods used and the apparent contradictions between the conclusions obtained in previous studies have created uncertainty in understanding the mechanisms of replication-division cycle coordination [33, 34]. Adding to this complexity is the variability in observations across growth conditions. For instance, in slow growth conditions, the division adder model fails to describe cell division [16], and replication appears to play a central role, while it becomes less important in fast growth conditions [30, 31]. Besides, under experimentally perturbed conditions, some results support the independence of the division from replication [18], while other results highlight the involvement of two concurrent processes in the division [29, 30].

In this work, using experimental data from unperturbed cells in [18, 19, 30] and building on previous analyses, we define a unified mathematical framework for studying replication-division coordination models. This enables us to relax the Gaussian assumption, which we show is not always satisfied experimentally. This approach allows us to gain a clearer qualitative understanding, define a new model and quantitatively compare model predictions with experimental data. We compare four models: a replication-independent one proposed by [18]; a division-independent one introduced in [19]; and two double-process models, one proposed by [27]; and a second one introduced here for the first time. We first derive theoretical results and show the necessary and sufficient conditions on single-process models to capture key physiological behaviours in *E. coli*. Our unifying framework then allows us to conduct a statistical analysis comparing models based on their ability to reproduce the conditional distribution of division volume, given the birth volume, initiation volume, and cell elongation rate. We propose a systematic comparison method that applies consistent evaluation criteria across models. Importantly, we infer the distribution of the so-called *latent* (i.e., not directly observable) variables involved in double-process models, which has never been done to date. Our results support the hypotheses underlying double-process models and lend support to a new model that reconciles previously conflicting evidence regarding the interdependence between replication and division cycles. Overall, this work presents a methodology and a model that open up new avenues for developing a universal replication-division coordination model.

## Results

### A unified mathematical framework for replication-division coordination

#### General assumptions and observed quantities

Multiple mathematical models have been proposed to describe the replication-division coordination [17, 27, 28, 18, 19]. To compare them and capture the mathematical differences between these models, we introduce a general framework which reproduces the monitoring of a cell lineage in a Mother Machine device [4]. At each generation, a mother cell is randomly replaced by one of its two daughters. In this framework, the following assumptions are done, and justified by biological observations.

#### Assumptions

1. The division is perfectly symmetric, i.e. the volume of the daughter cell at birth is equal to half of the volume of the mother at division [11, 35].
2. At each replication initiation event, every origin of replication in the cell fires a new round of replication at the same time, thus doubling its number [16].
3. The origins of replication of a mother cell are equally partitioned into the daughters at division [16].
4. Over the lineage, the cell volume grows exponentially according to a cell elongation rate function of time *λ*(*t*), which is positive and locally integrable [4].

In this framework, two main quantities are studied: the cell volume, and the volume per origin of replication. According to Assumptions 1 and 4 the cell volume trajectory is continuous except at division events, when it is halved. According to Assumptions 2 and 4 the volume per origin of replication is continuous except at initiation events, when it is halved, and remains continuous at division events according to Assumptions 1 and 3. In consequence, the total cell volume is governed by the cell elongation rate and the division cycle, while the cell volume per origin is governed by the cell elongation rate and the rounds of replication initiation. We focus on these two quantities as their coupling directly relates to the coupling between the division cycles and the replication cycles.

These two quantities link one generation to the next along a given lineage. We define 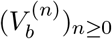, the sequence of birth volumes, where for any generation *n*, 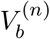 represents the birth volume of the mother cell giving rise to a daughter cell with a birth volume 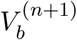. Identically, we define 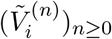 as the sequence of volumes per origin of replication measured immediately after each initiation event. For every *n*, the initiation corresponding to 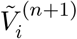 is the one that occurs directly after the initiation associated with 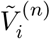 (see Mathematical Supplementary). In the context of a dynamics over time, we additionally introduce 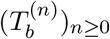 the dates of division resulting in a cell of volume 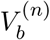, and 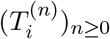, the dates of the initiation of replication for the cell with a volume per origin of replication 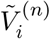. The indexation of all these sequences is set in such a way that the sister chromosomes resulting from the replication initiated at 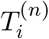 are separated by the division occurring at 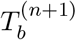, defining the difference 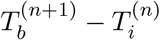 as the C+D period for every *n*. Using a specific assumption over the initialisation of the sequences, one can show the following expression (see Mathematical Supplementary, Lemma 1):

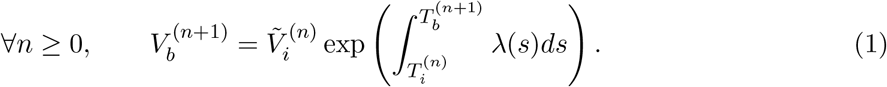

This expression is true regardless of whether the C+D periods overlap or not and thus is true for any growth condition. Additionally, we monitor the number of origins of replication within a cell at birth (which can be more than one if the C+D periods overlap). Let (*ω*^(*n*)^)_*n≥*0_ be the number of origins of replication at the *n*-th birth. This quantity is defined on the space *{*2^*k*^ : *k ∈* ℤ*}*, since it is doubled at each initiation event and it is halved at each division. Note that to make sense from a biological point of view, this quantity should be greater or equal to one. However, we do not impose a priori a condition over the sequences 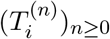 and 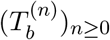 to respect this constraint.

One can show, under the condition that 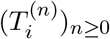 is non-decreasing, that this quantity satisfies (see Mathematical Supplementary, Lemma 2), for every *n ≥* 0 and *k ≥ −n* + 1 :

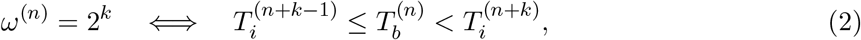

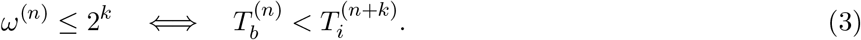

### Three key models from the literature

Here, we introduce and formulate three models from the literature that have drawn the most interest among the single-process and double-process models: the Double Independent Adders Model (DIAM), the Replication Double Adders Model (RDAM), and the Concurrent Processes Model (CPM).

All these models assume that replication initiation events are triggered according to an initiation adder [5, 19, 18, 17]. The initiation adder states that the initiation of replication is triggered once the cell lineage has accumulated a critical volume per origin of replication since the last initiation (Figure 2A). This can be formulated mathematically as follows. Let 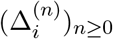 be an independent and identically distributed (i.i.d.) sequence of positive random variables that admit a density 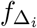, modeling the added volume per origin between the *n*-th and *n* + 1-th initiations (Table 1). The sequence 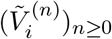 satisfies according to the initiation adder,

**Table 1.**
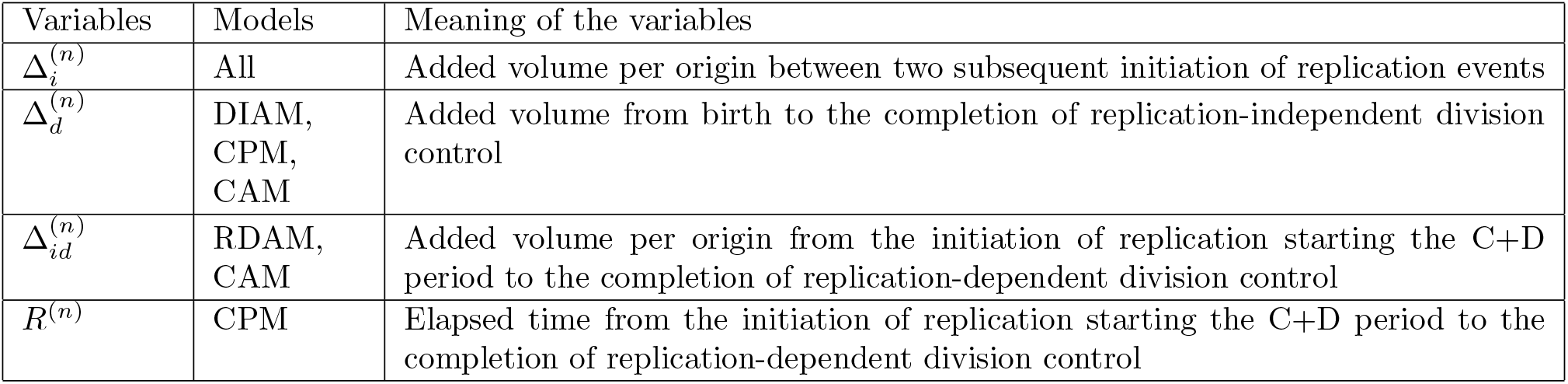
Random variables involved in the models and their meaning.

**Figure 2.**
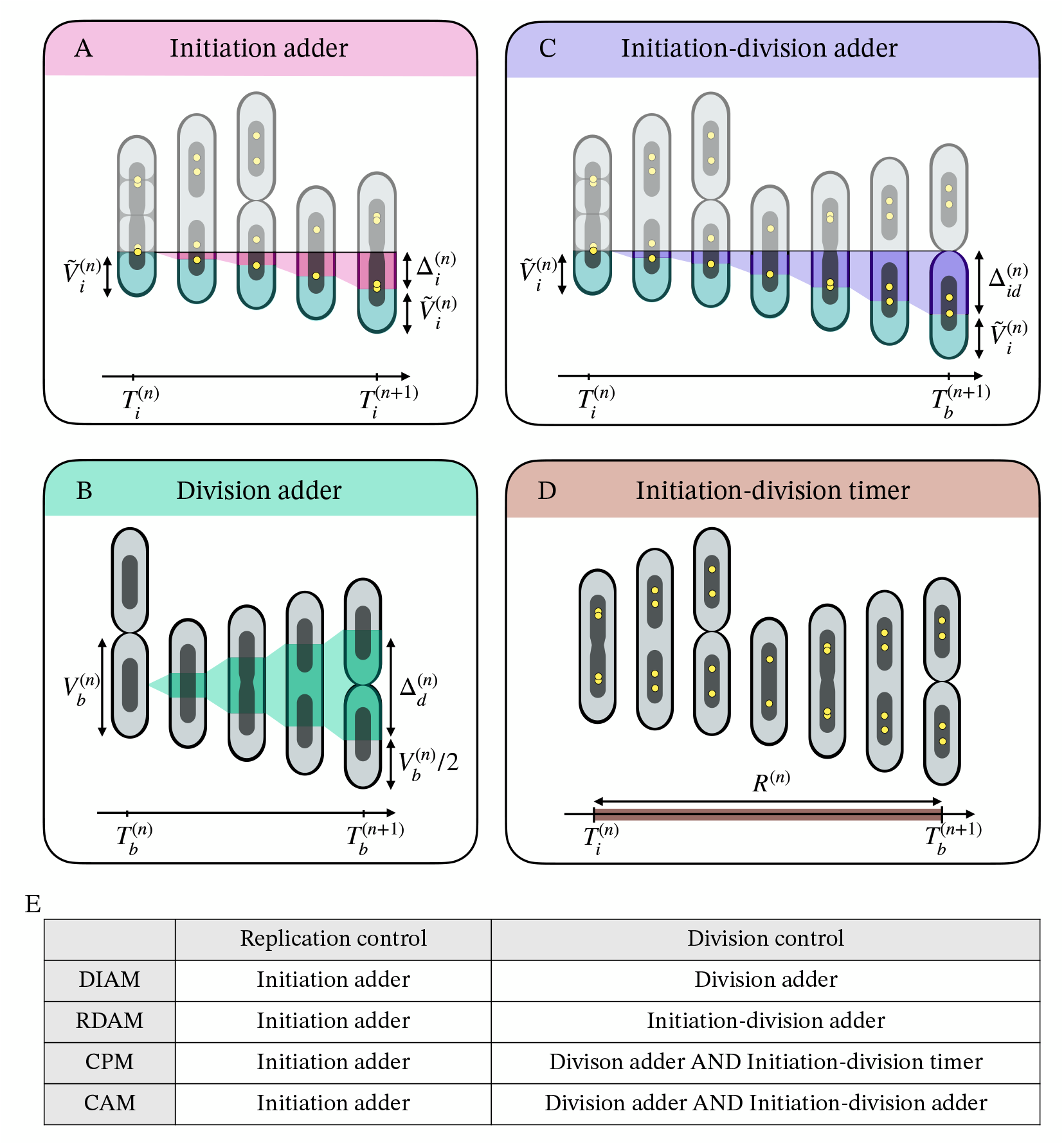
Summary of the models studied. (A) The initiation adder model assumes that the initiation of replication event setting the initiation volume per origin 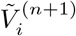 is triggered once the cell has added a volume per origin of replication 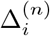 assumed to be independent of 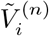, since the last initiation event. (B) The division adder assumes that the division occurring at 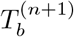 is set once the cell has added a volume 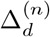 since birth, assumed to be independent from 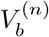. (C) The initiation-division adder assumes that the division occurring at 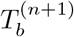 is set once the cell has added a volume per origin 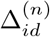 since the beginning of the C+D period, assumed to be independent to 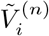. (D) The initiation-division timer assumes that the division occurring at 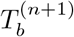 is set after an amount of time *R*^(*n*)^ after the beginning of the C+D period, assumed to be independent from 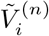. Table (E) displays the control mechanisms of replication cycles and division cycles for each model studied in this work.

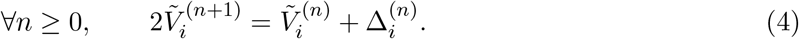

The **DIAM**, proposed by [18], supposes that the cell cycles are headed by a double adder mechanism, assumed to be independent: the initiation adder and the division adder (Figure 2B, E). This decoupling between replication and division makes 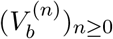 and 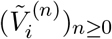 independent processes.

Similarly as the initiation adder, one may introduce 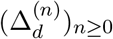, the i.i.d. random variables drawn from a density 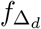, that model the added volumes over cell generations (Table 1). The couple 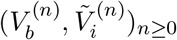 satisfies according to the DIAM:

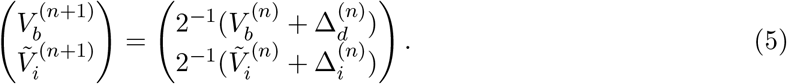

The **RDAM**, first proposed in *Mycobacterium smegmatis* [36], and then adapted in *E. coli* by [19], suggests that replication is central to the control of division. This model assumes an initiation-division adder such that division is triggered once the cell lineage has added a critical volume per origin formed at initiation over the corresponding C+D period (hereafter referred to simply as the added volume per origin over the C+D period) (Figure 2C, E). Let 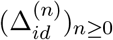 be an i.i.d. sequence of positive random variables of density 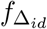 that models the added volumes over the C+D periods (Table 1). The couple 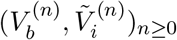 satisfies according to the RDAM:

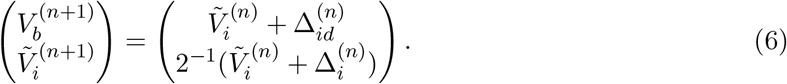

The **CPM** proposed by [27, 28] assumes that the division is set by the slowest of two independent concurrent processes: a replication process and a division-related process [27, 29, 30]. According to this model, a cell divides after adding a critical volume since birth AND completing chromosome replication and segregation. In this model, replication and segregation are supposed to follow an initiation-division timer (Figure 2D, E). From the assumptions of the model, we can write the sequence of volumes at birth 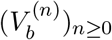 as the maximum between a timer over the C+D period modeled by the i.i.d. sequence 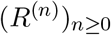 drawn from the density *f*_*R*_, and a division adder modeled by the i.i.d. sequence 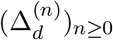 drawn from the density 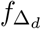(Table 1):

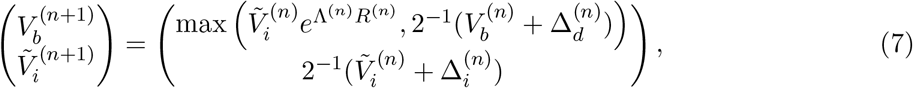

With 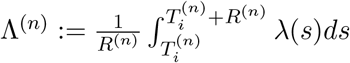 . As cell elongation has been shown to be approximaltely exponential (*λ* constant over time despite stochastic fluctuations) [9, 37, 38], in the following we will do the approximation that (Λ^(*n*)^)_*n≥*0_ and (*R*^(*n*)^)_*n≥*0_ are independent.

Every model introduced so far supposes 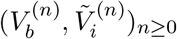 to be a homogeneous Markov chain, with the state of the chain at *n* + 1 depending on the state at *n*, and some i.i.d. random variables. Note the implicit assumptions made by assuming these random variables to be identically distributed are that the growth conditions are constant and that no ageing is happening.

Among the models considered, the DIAM and RDAM both involve two i.i.d. random variables, while the CPM incorporates three i.i.d. random variables, introducing an additional degree of complexity and making model comparison less straightforward. Furthermore, one difficulty lies in inferring the distribution of the latent random variables Δ_*d*_ and *R*, in the CPM: because this model relies on taking the maximum of two quantities, the measured added volumes do not directly correspond to observations of Δ_*d*_ and similarly, the measured C+D periods do not directly correspond to observations of *R*, see Figure S2. This implies that the distributions of Δ_*d*_ and *R* cannot be directly deduced from the experimental observations of the added volumes and C+D periods.

### The CAM: a new model that brings together all three models

Along with these models, we introduce a novel one called the Concurrent Adders Model (CAM) that combines previous hypotheses differently. The CAM is a CPM-like model that incorporates replication-based division control via an initiation-division adder (as in the RDAM) rather than the initiation-division timer used in the CPM (Figure 2 C, E). The size control is ensured by a division-adder mechanism (as in the DIAM), and the division occurs at the completion of both processes (as in the CPM). In line with the other models, we assume that the CAM also involves an initiation adder. Let 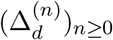 and 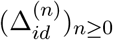 be i.i.d. sequences of positive random variables with respective densities 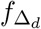 and 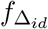 (Table 1). The couple 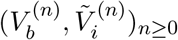 satisfies according to the CAM:

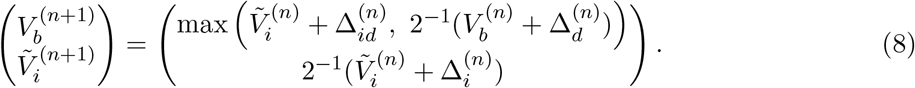

Similarly to the CPM, the random variables Δ_*d*_ and Δ_*id*_ are latent, so their distribution cannot be inferred directly from the experimental observations of inter-division added volume and added volume per origin over the C+D period.

### Conditions for the validity of the DIAM and the RDAM

Stating that division and replication are decoupled processes as proposed by the DIAM, implies that the sequences 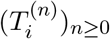 and 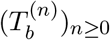 are independent and thus events such as 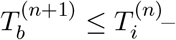 interpreted as negative C+D periods – could occur if no additionnal assumptions are made.

Similarly, if cell division is solely regulated by replication, as proposed by the RDAM, this implies that the sequence 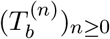 depends exclusively on 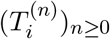 and 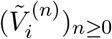, without explicitly considering the timing of the previous divisions. Consequently, events such as 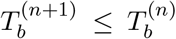, where a daughter cell divides before its mother, may not necessarily be prevented in a stochastic model with no additional assumptions. Here, we show that single-process models may lead to inconsistencies without additional assumptions, and we provide necessary and sufficient conditions for the DIAM and the RDAM to avoid these irregularities.

As in the DIAM division and replication are independent processes, one could envisage a case where a cell lineage accumulates more division events than initiation events, which would lead to the formation of an anucleated cell [39]. In the context of our framework, this event corresponds to the birth of a cell with less than one origin of replication. Below, we explore the distribution of the number of origins of replication at birth, which provides the probability of such an event. We first prove that after a large number of generations, the distributions of birth and initiation volumes 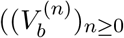 and 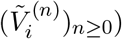 both converge to a stationary distribution, (Propositions 1 and 2 in the Mathematical Supplementary). This allows us to express the stationary distribution of the limit number of origins of replication at birth, denoted *ω*_*∞*_, as:

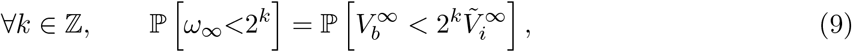

with 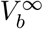 and 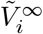 the random variables limit in distribution of 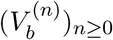 and 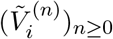 respectively. Using this distribution, we show that the number of origins of replication at birth *ω*_*∞*_ is greater or equal to one almost surely if and only if:

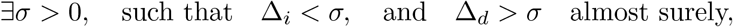

see Theorem 1 in the Mathematical Supplementary. These results provide the necessary and sufficient condition to apply to the DIAM to prevent the division of a cell with one origin of replication and show that when it is met, the DIAM ensures the homeostasis of the number of origins of replication despite the lack of coupling between replication and division cycle. This condition is satisfied if the empirical distributions of Δ_*d*_ and Δ_*i*_ do not overlap, which can be checked in the experimental data. Figure 3 shows the empirical distributions of Δ_*d*_ and Δ_*i*_ obtained in slow growth conditions in [18, 19]. As highlighted by the black circles, the supports of the distributions of Δ_*i*_ and Δ_*d*_ do overlap, suggesting that if the cell cycles are controlled following the DIAM, in these particular conditions, there is a positive probability for a cell to divide when it has only one origin of replication, leading to an anucleated cell. From these experimental distributions, it is possible to simulate the distributions of 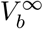 and 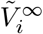 and estimate this probability as it is equal to 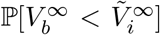 (see Figure 3 and Material and Methods). Through a Monte-Carlo method, we computed the probability of having an anucleated cell and showed that it is in the order of 10^*−*3^, and 10^*−*4^ according to the data of [18] in MG1655 acetate and NCM3722 arginine, respectively, and 10^*−*3^ in glycerol according to the data of [19]. This should lead to relatively frequent anucleated cells, which has not been reported, suggesting that an additional control process is necessary. To better understand the predictions of the DIAM in these conditions, we compared simulated and experimental data. As illustrated in Figure S1, in the experimental data, bacterial cells that have a large initiation volume also have a large birth size, making the experimental distribution strikingly different from the simulated data from the DIAM. Moreover, the DIAM also predicts outliers in the “forbidden” region, as seen by cells admitting a smaller birth size 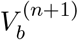 than the initiation size per origin 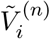. This further suggests that, at least in specific experimental conditions, single process models such as the DIAM cannot fully recapitulate the data.

**Figure 3.**
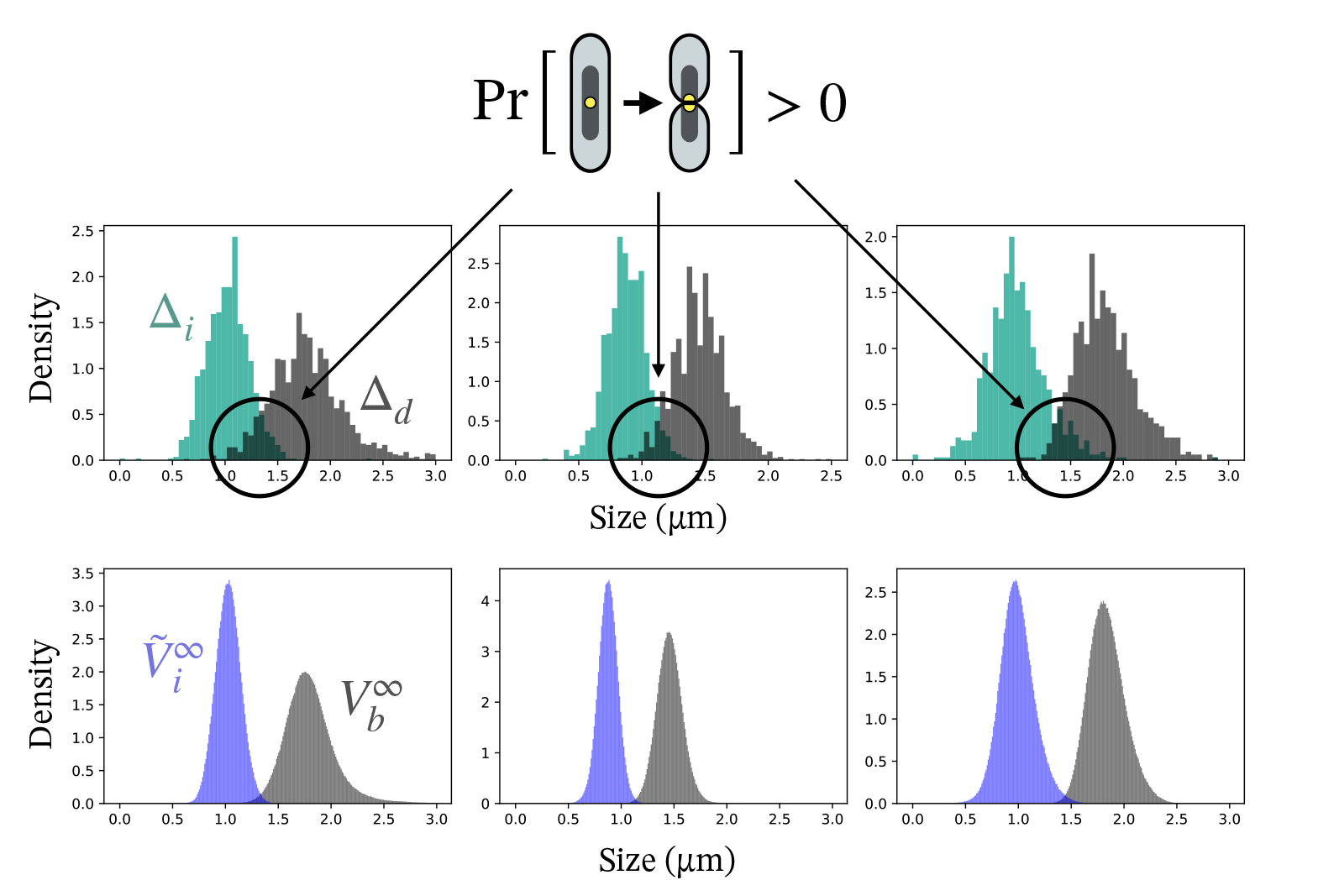
Probability of one-origin-cell division according to the DIAM. (Top) Empirical distributions of Δ_*i*_, and Δ_*d*_ in MG1655 acetate, NCM3722 arginine and BW27378 glycerol from left to right, taken from [18, 19]. The overlap of the distributions of Δ_*i*_, and Δ_*d*_ indicates a positive probability for cells to divide with one origin of replication. (Bottom) Simulated distributions of 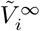 and 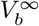 from Monte Carlo method, which allowed to compute the probability of one-origin-cell division (see Material and Methods).

When considering the RDAM, due to the stochastic nature of the added volume per origin over a C+D period Δ_*id*_, it is theoretically possible that a C+D period may start and finish within the duration of the previous one. This scenario would imply that the (*n* + 1)-th birth could occur before the *n*-th birth, meaning that a daughter cell could divide before its mother. We show in Proposition 4 that this scenario is unlikely (of probability 0) if and only if

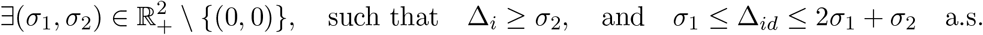

These conditions involve restrictions on the supports of the densities of Δ_*i*_ and Δ_*id*_. One can estimate the probability of a daughter cell dividing before its mother from the observations of Δ_*I*_ and Δ_*id*_ (see Material and Methods). Based on datasets published in [18] and [19], we show that this probability is positive in every condition: ranging from approximately 10^*−*4^ for the strains MG1655 [18] and BW27378 [19] to 10^*−*8^ in the strain NCM3722 [18] (see Table S1).

Taken together, the results of this section indicate that accounting for a unique process - either coupling the initiation of replication or the birth with division may lead to physiological inconsistencies with positive probability, and thus might not be enough to fully capture the underlying biological process. However, this might only be relevant in specific growth conditions. In the following section, we expand our analysis to statistically compare single-process models with double-process models in a systematic way in multiple growth conditions.

### Distribution inference and systematic comparison of replication-division coordination models

So far, the comparison of replication-division coordination models has mainly relied on assessing their ability to reproduce the correlations seen in the data. These analyses have suggested that certain cell variables are independent, based on the observation of the absence of correlation [18, 19, 26, 32, 31]. This conclusion holds only if the variables of interest are normally distributed. However, Shapiro–Wilk normality tests indicate that division, birth, and initiation sizes deviate significantly from normal distributions (see Table S2). Moreover, correlation analysis gives a limited indication of the performance of the double-process models, as they are based on non-linear relationships between cell variables, which has led to the use of approximations in previous studies [27, 28, 29, 31]. Here, to go beyond correlations, we assess models on their abilities to reproduce the joint distribution between cell variables. Specifically, as our main question is to understand what controls the division volume, we chose to evaluate the four models previously introduced according to their ability to reproduce the division volume distribution conditioned on three covariates: the volume at birth, the volume per origin at replication initiation of the corresponding C+D period, and the cell elongation rate over this C+D period.

To this end, we analysed 17 datasets from [18, 19, 30], which report measurements obtained using microfluidic Mother Machine devices. These datasets include information on cell length at birth and division, cell length per origin at initiation and C+D period duration. As cell width is constant over time in a given growth condition, we used cell length as a proxy of cell volume [35, 40].

### Conditional distributions of division volume

Here, we define the conditional distribution of division volume given birth and initiation volume and cell elongation rate over the C+D period. In our framework, this corresponds to the analysis of the distribution of 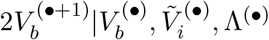. As shown in the Mathematical Supplementary, Proposition 5, the conditional distribution of the volume at division 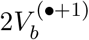, given that 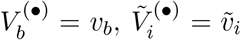, and Λ^(*•*)^ = *λ* satisfies for the different models the following expressions for any *v >* 0:

- The DIAM:

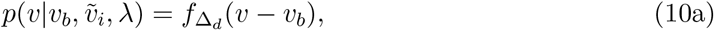
- The RDAM:

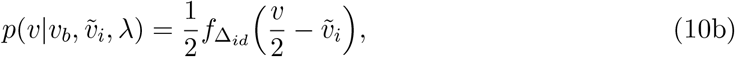
- The CPM:

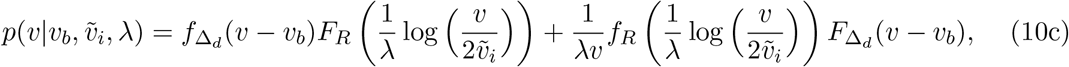
- The CAM:

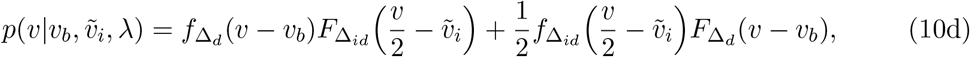

With 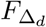, 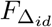 and *F*_*R*_ the cumulative distribution functions of Δ_*d*_, Δ_*id*_, and *R* respectively.

### Parameter inference for each model

As already mentioned, Δ_*d*_ and Δ_*id*_ are directly observable in the DIAM and RDAM, so that a nonparametric estimation can be carried out [35, 41, 42]. However, this is not the case for the CAM and the CPM, for which the use of latent variables makes nonparametric estimation impractical. To provide a unified framework for model comparison, we therefore used a parametric inference method. In the following, we assume that the i.i.d. random variables involved in all models follow parametric densities, specifically, Generalized Gamma distributions. This three-parameter distribution gathers a broad family of unimodal probability distributions supported on R_+_, such as the Gamma, Weibull, Log-Normal, inverse-Gamma, inverse-Weibull distribution and others [43]. As can be observed in Figure S3, this family of parametric distributions allows us to fit in a satisfactory manner all the empirical distributions we extract from the data, such as the inter-division added volume [44], the C+D period, and the added volume per origin over the C+D period. This is why we used this parametrisation for the density of the random variables Δ_*d*_, Δ_*id*_ and *R*, even if Δ_*d*_, *R*, and Δ_*id*_ cannot be directly measured in the CPM and the CAM. In the absence of well-established molecular mechanisms underlying the control variables of the models, this assumption provides flexibility, allowing the comparison below to avoid potential bias introduced by overly specific constraints on the distributions.

For a given dataset 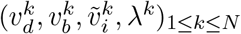 containing the measurements for *N* cell cycles of the division volume 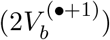, birth volume 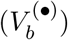, initiation volume per origin 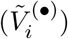, and cell elongation rate over the C+D period (Λ^(*•*)^), we define the likelihood for each model:

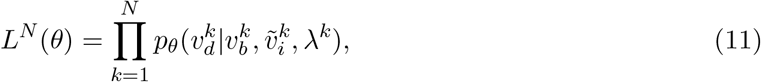

with *p*_*θ*_ the conditional distribution of division volume defined by either (10a)–(10d), with *θ* containing the parameters of the generalised Gamma distributions of the respective i.i.d. random variables.

The distribution parameters are estimated by maximising this likelihood (see Materials and Methods). For all four models and all 17 datasets tested, we observe that the predicted distributions for the volume at division correctly recapitulate the empirical distributions (Figure S4).

In the DIAM and in the RDAM, the variables Δ_*d*_ and Δ_*id*_ are directly observed making the parameters of their distribution identifiable. In these cases, Formulae (10a) and (10b) show that maximising the likelihood (11) amounts to estimating the parameters of the generalised Gamma distribution describing respectively Δ_*d*_ and Δ_*id*_ (Figure S3, first two columns). In contrast, in the CPM and CAM, the variables Δ_*d*_, *R*, and Δ_*id*_ are latent, and thus their parameters’ identifiability is not guaranteed. In Proposition 6 (see Mathematical Supplementary), we provide a sufficient condition for it. We then verified that this condition is met in all datasets. In Figure S5, we compare the distributions estimated for *R* and Δ_*d*_ (respectively for Δ_*id*_ and Δ_*d*_) in the CPM (respectively the CAM) to the empirical data. This revealed significant discrepancies, as expected, because latent variables do not directly correspond to empirical data, which are the observation of the maximum of the two latent variables. In Figure S1 we illustrate these differences and their interpretation. For example, if the replication-independent process is the limiting one (i.e. the slower one) in a given condition, then its related latent parameter is close to the empirical distribution, while the parameter describing the replication-dependent process is far from its empirical distribution. This allows us to estimate the probability that one of the two processes is dominant in the two-process models (Figure S6).

### Model comparison using BIC: the double-process models best predict division size control

After finding *θ*^*\**^ maximizing the likelihood (Equation 11) for each model, one can compare them by computing the Bayesian Information Criterion (BIC), defined as *BIC* = *−* log(*L*^*N*^ (*θ*^*\**^)) + *ρ* log(*N*)*/*2, with *ρ* being the number of parameters (*ρ* = 3 for the DIAM and the RDAM; *ρ* = 6 for the CPM and the CAM). The BIC allows model comparison based on an approximation of their posterior probability given the data. A lower BIC value indicates a greater probability that a model actually explains the data and penalizes models with a higher number of parameters thus ensuring a fair comparison [45]. For a complete comparison, we also computed the Akaike Information Criterion (AIC) of the models, see Table S3 and Figure S7. We chose to base our conclusions on the BIC criterion because its penalty term penalizes the double process models more strongly, thus limitting overfitting.

We computed the BIC score over the 17 datasets for each model (Table S3, Figure 4). Firstly, across most datasets, the double-process models, namely the CPM and the CAM provided the best predictions of division size (conditionally to the birth and initiation size and cell elongation rate), in a large majority of datasets. This indicates that accounting for both replication-dependent and replication-independent processes is essential to describe division size control accurately. Secondly, under slow growth conditions (elongation rate less than 0.5 *h*^*−*1^), the CPM, the CAM and the RDAM consistently outperform the DIAM, which relies solely on birth volume. This suggests that in slow growth conditions, replication is a key constraint in the control of division, consistent with recent results [29, 30, 31]. Finally, the CAM and the RDAM show lower BIC scores than the CPM under slow growth conditions (elongation rate less than 0.3 *h*^*−*1^). Since both the CAM and the RDAM implement an initiation–division adder, in contrast to the initiation–division timer in the CPM, this result suggests that replication-based control of division under these conditions is more accurately described by an adder-like mechanism. Notably, in these conditions, the CAM predicts a probability of replication-limited division approaching 1, whereas the CPM never exceeds 0.5 (see Figure S6). This indicates that, in slow growth regimes, the CPM fails to capture the strong coupling between replication and division, which may explain its lower predictive performance compared to the CAM and the RDAM.

**Figure 4.**
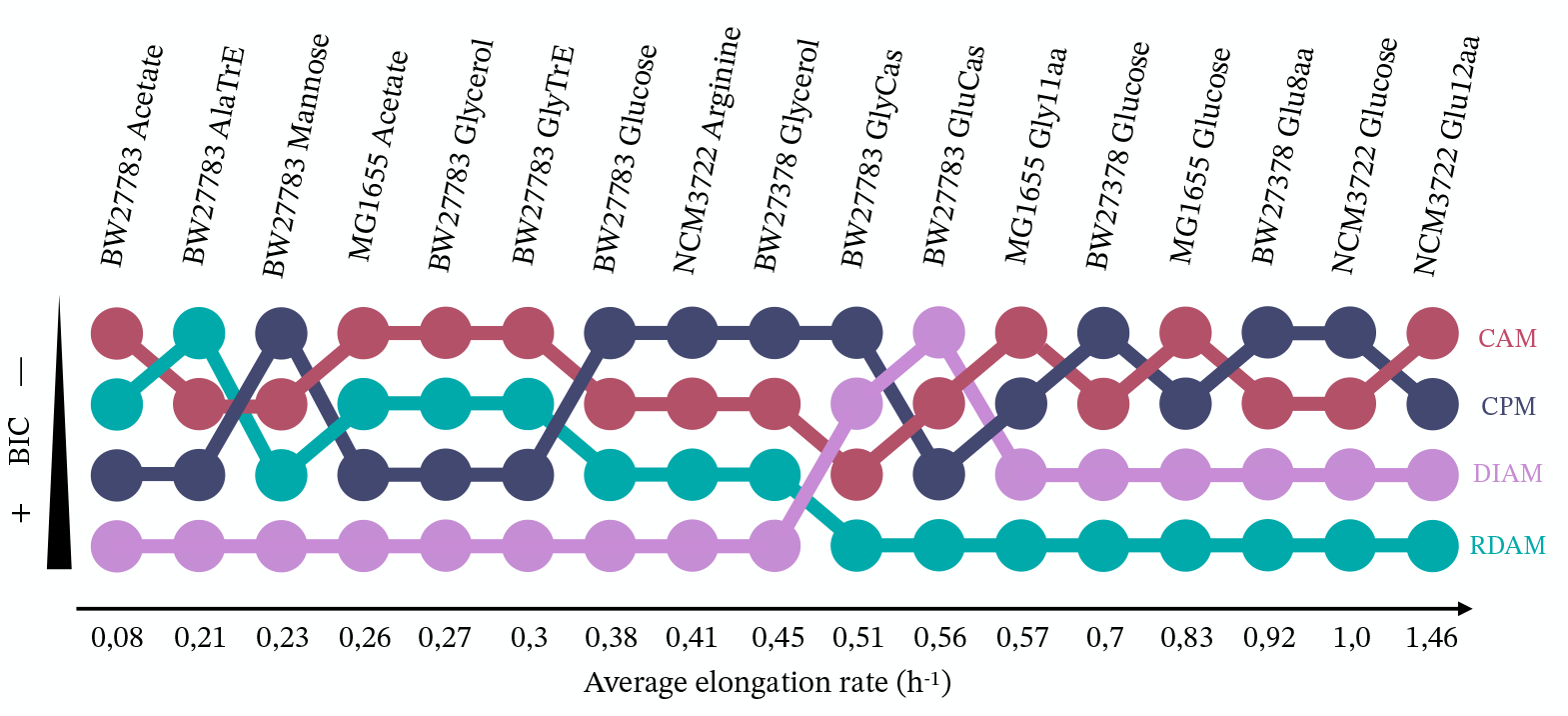
Comparison of the performances of the models based on the BIC. The BIC scores have been computed for each model (DIAM in pink, RDAM in cyan, CPM in dark blue and CAM in dark red) for multiple datasets of four strains. Here are shown the ranks of the models based on their BIC score, with a lower BIC score indicating a greater probability that a model actually explains the data.

### Comparison of initiation-division sizes joint distributions using Wasserstein distance: CAM best reconstructs initiation-division sizes dependence

The results above, based on maximum likelihood comparison using BIC, suggest that the CPM and the CAM best capture the replication-division coordination over a wide range of growth conditions, without a clear preference for one or the other in intermediate and fast growth condition. However, while BIC highlights the overall likelihoods of the models, it does not provide clear information on how each model reconstructs the dependence between cell variables. Comparing models based on their ability to reproduce the pairwise dependencies between cell variables provides additional insight and discriminative features, as proposed in [35, 46]. Specifically, the CAM and the CPM differ in how they integrate initiation size in the division control. Therefore, our objective in this section is to compare these models in their ability to reproduce the dependence between initiation size and division size and to compare to experimental data. By simulating division sizes conditionally on measured cell covariates, using best-fit parameters from the maximum likelihood, one can assess which model between the CPM and the CAM best captures the joint distribution of initiation and division sizes. To illustrate this, we display in Figure 5A the heat maps of the experimental joint distribution of division and initiation sizes from selected datasets (top, black) and their counterparts when division size is simulated under either the CAM (middle, red) or the CPM (bottom, blue). Visual inspection suggests that the CAM produces distributions that are closer to the data than the CPM in the MG1655 Acetate (a) and BW27378 Glycerol (c) datasets, while the difference is less pronounced for the MG1655 Glucose (b) and BW2738 Glucose8a (d) datasets. To quantify the dissimilarity between joint distributions, previous studies have used a distance between the densities of the distributions [35, 47]. However, this method requires regularising the experimental data. To avoid this regularisation step, we directly compared the experimental data to the one simulated under the CAM and the CPM using the Wasserstein distance (also known as the Earth Mover’s Distance). This distance is the natural metric to compare empirical distributions and is computed as the minimum average distance between paired points of two empirical distributions (see Figure 5B) [48, 49]. We used *W*, the Wasserstein distance between the experimental joint distribution of initiation and division volumes 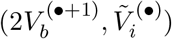, and the joint distribution obtained when the division volume is simulated according to either the CPM or the CAM (see Figure 5B).

**Figure 5.**
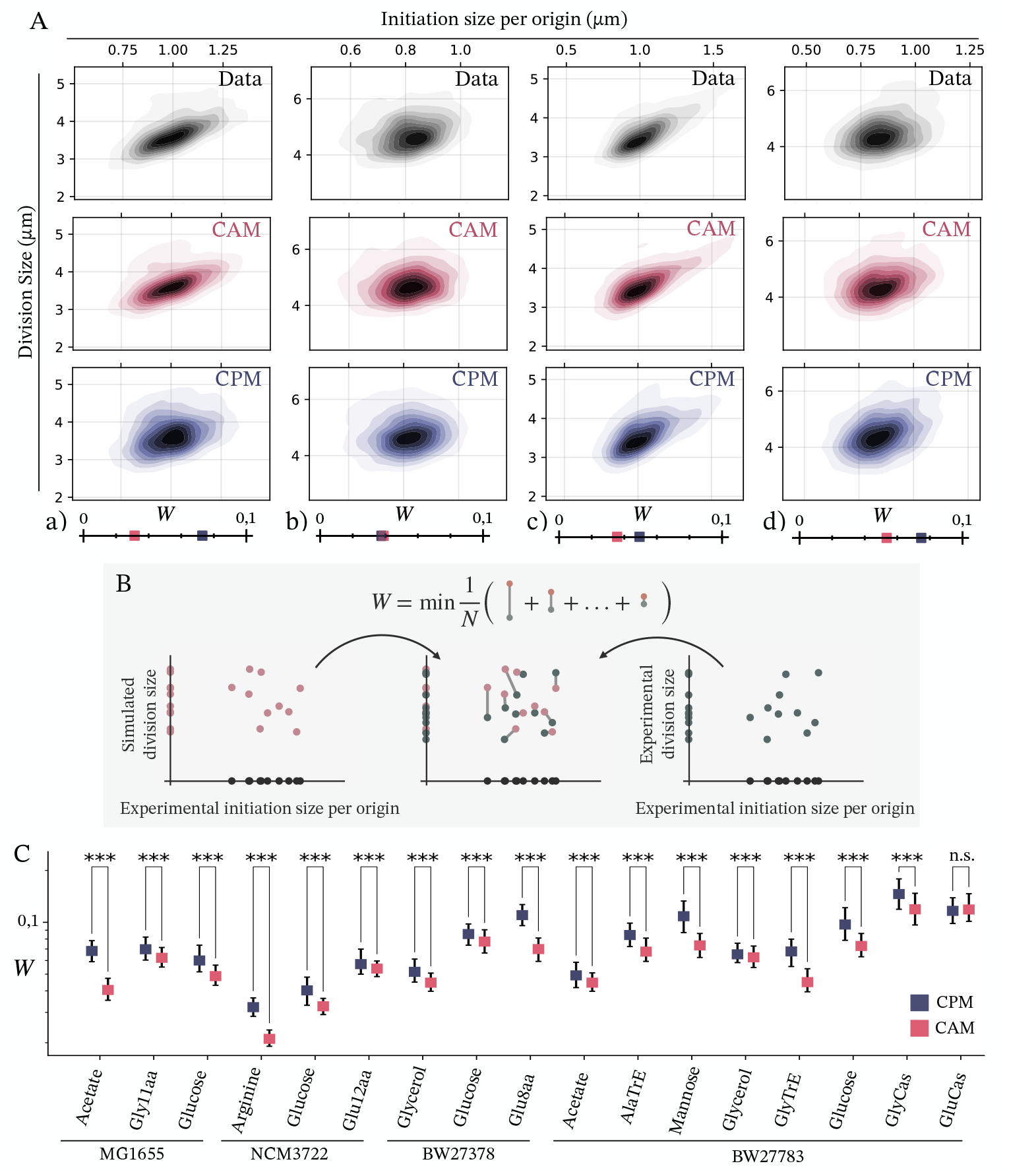
The CAM best reconstructs initiation-division sizes dependence. (A) Examples of joint distributions between division size and initiation size from datasets ((a) MG1655 Acetate, (b) MG1655 Glucose, (c) BW27378 Glycerol, (d) BW27378 Glucose8a), and corresponding simulations of the CAM and the CPM. At the bottom are mapped the according Wasserstein distances between the data and the simulations (CPM in blue and CAM in red). (B) The Wasserstein distance between two empirical joint distributions represents the minimum average distance between paired points. This measures the dissimilarity between two empirical distributions. Intuitively, a greater Wasserstein distance indicates a greater dissimilarity. In this work, we compute the Wasserstein distance *W* between the experimental joint distribution of division size and initiation size, and the corresponding distribution when the division volume is simulated according to the CPM or the CAM. (C) Wasserstein distances between experimental and simulated joint distributions of division and initiation sizes over 100 simulations according to the CPM and CAM. Significant differences according to Wilcoxon-Mann Whitney tests are indicated: ***p-value *<* 0.001, n.s.= not significant (p-value *>* 0.05).

We computed this distance using 100 simulations under each model. As shown in Figure 5C, *W* between the experimental data and the data simulated under the CAM is smaller in average than *W* between the experimental data and the data simulated under the CPM in 16 of the 17 datasets. This difference is statistically significant, as confirmed by the Wilcoxon-Mann-Whitney test (see Materials and Methods). These results indicate that the CAM more accurately reconstructs the joint distribution of division and initiation sizes, reflecting a better reproduction of the dependence between these two variables. By computing similarly the Wasserstein distance of the joint distribution of division and birth sizes, we also found that the CAM more accurately reconstructs the dependence between these two variables (see Figure S8). Taken together, these results suggest that the initiation-division adder better reproduces the replication-based control of division under every growth condition, compared to the timer mechanism used in the CPM.

## Discussion

In this work, we proposed a framework to analyse and compare models that describe the coordination between replication and division cycles in *E. coli*. We formulated four models with different degrees of complexity using a unifying generation-wise setting and systematically compared them.

We relaxed the Gaussian assumptions required in previous correlation-based studies, included nonlinear models (without relying on linear approximations), and were able to estimate distributions of latent control variables. Two of these models are based on the assumption that division is controlled by a single process: the DIAM, which assumes that division is solely triggered according to a division adder; and the RDAM, which places replication at the center of division control, assuming that cells divide according to an initiation-division adder. The two other models assume that cell division is controlled by the concurrence of two processes: the CPM assumes that division occurs once two independent processes are completed, one corresponding to a division adder and the other to a timer over the C+D period; and the CAM, a novel model proposed here, which is similar to the CPM except that it considers the replication-based control of division as an initiation-division adder, as in the RDAM.

We first used our framework to derive the conditions of validity of both single-process models and to test whether these conditions are met in the data. This theoretical study led to the conclusion that if cells follow the DIAM, cell division with one origin of replication resulting in anucleated cells should happen with a probability up to 10^*−*3^ in certain datasets. As anucleated cells are very rarely reported in WT strains of *E. coli*, this suggests that, at least in certain growth conditions, the DIAM may not be robust enough to describe finely the coordination between replication and division cycles. As for the RDAM, it opens the possibility that daughter cells would divide before their mother with a probability in the order of 10^*−*4^ an event which cannot be interpreted biologically - again suggesting limitations in the robustness of this model. However, we were able to provide necessary and sufficient conditions for these models to avoid unphysiological events. Specifically in the DIAM, we provide conditions regarding the supports of the densities 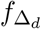 and 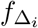 to ensure that the number of replication origins at birth remains greater than or equal to one with probability one. This finding suggests that there are conditions where independent replication and division processes can maintain the number of replication origins within physiological ranges. This opens the possibility of engineering synthetic cells with balanced coupling of replication and division cycles driven solely by independent molecular networks, thus greatly simplifying the control of synthetic minimal cells [50].

We then used our framework to develop a statistical method for systematic comparison of models, based on their ability to reconstruct the conditional distribution of the division size. We computed the likelihoods constructed from their respective conditional distribution, which allows us to estimate the distributions of the control variables and to compare model performance using BIC. The results suggest that double process models provide better predictions of division size than single-process models in every growth condition and highlight that both initiation and birth size are key variables in the division control, even under slow growth conditions, contrary to previous conclusions [31]. Moreover, this analysis revealed that the CAM performs better than the CPM, specifically in slow growth conditions. To better understand the predictions of the double-process models, we simulated the data under the CAM and CPM using the best-fit parameters and then compared them to experimental data employing the Wasserstein distance. The additional insights gained from this analysis allowed us to demonstrate that the CAM produces simulated data that closely resemble the experimental data and more accurately reconstructs the relationship between division and initiation size than all three other models (Figure 5, S9-S10) under most growth conditions. Taken together, our statistical analyses thus support the hypothesis that division is controlled by two concurrent processes under every growth conditions, which includes an initiation-division adder.

In [18] the DIAM was experimentally tested by showing that perturbing the initiation volume through synthetic oscillations of the replication initiator protein DnaA, barely affected the volume at birth, suggesting that division is independent of replication. Indeed, when we simulated oscillations in the initiation volume under the DIAM, we observed that the birth volume remained unaffected (Figure S11) while simulations under RDAM show minor oscillations. These results may appear at first glance to contradict the conclusion that double-process models are necessary to explain the replication-division coordination [18]. To settle this contradiction, we also simulated similar perturbations under the CAM and the CPM and observed how they may affect the volume at birth. As shown in Figure 6, under the CAM, our simulations display similar behaviour to that observed in the experiment: while the initiation volume oscillates, the birth volume remains largely unaffected. This indicates that small perturbations in the initiation volume may not cause a change in the birth volume, suggesting that this experiment cannot exclude all double process models. In fact, this finding implies that the molecular mechanisms underlying the CAM might play a key role in ensuring robustness in the regulation of cell division size, thereby buffering cells against minor perturbations in initiation volume. However, when applied to the CPM, simulations reveal that under oscillations of the initiation size, the birth size also strongly oscillates, implying that that this model is less consistent with the experimental observations of [18] due to its high sensitivity to variations in the initiation volume (Figure 6).

**Figure 6.**
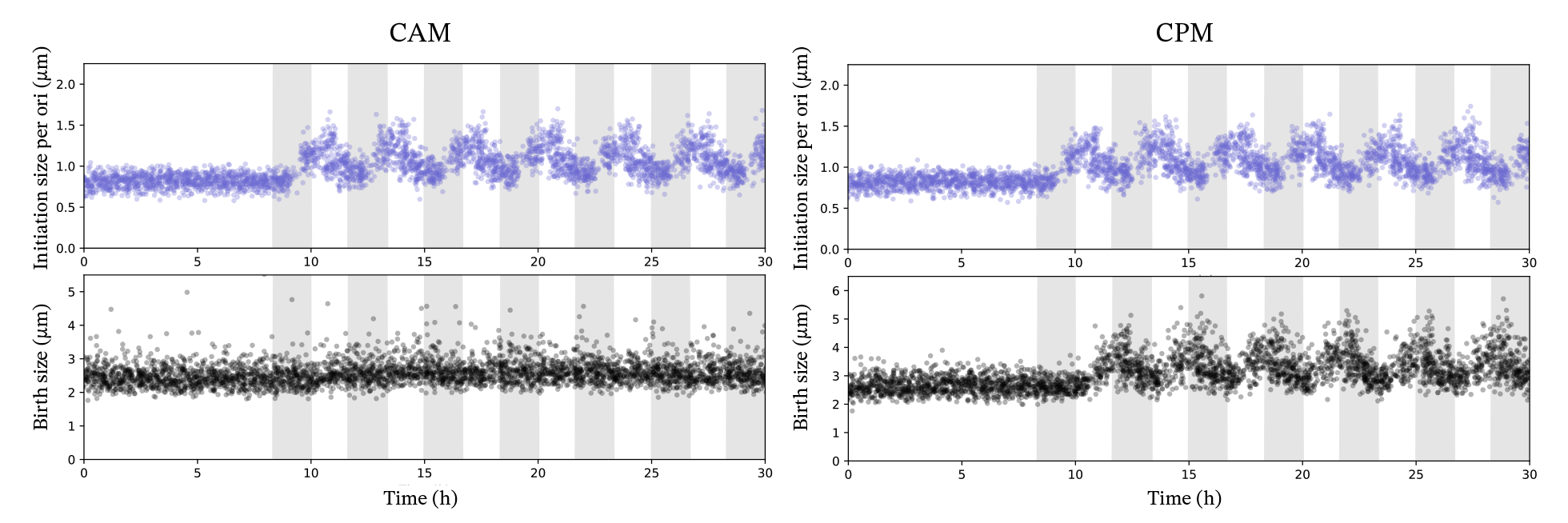
Simulation of the CAM(Left) and the CPM (Right) under oscillating distribution of Δ_*i*_. In this simulation, we follow 100 lineages in which Δ_*i*_ is multiplied by a given factor and returns to normal periodically with a period equal to 4 times the generation time (see Methods). The grey shaded areas correspond to the periods during which the Δ_*i*_ is increased. At the top, is displayed the initiation size per origin and at the bottom is displayed the birth size.

Additional elements of our analysis support the CAM. We provide in Figure S6 an estimation of the probability that replication is the triggering mechanism for cell division as a function of growth rate in the CAM and the CPM. We observe that contrary to the CPM, the CAM strongly supports a dominant role of replication in controlling division in slow growth conditions. This is consistent with previous results in which it has been observed that the onset of cell constriction is blocked by nucleoid occlusion and that this process is mainly limiting for cell division in slow growth conditions [30][31]. Finally, a very recent study has proposed a new model of replication-division coordination, supported by new experimental results, which is a modified RDAM where 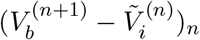 is not i.i.d. but a Markov chain [51]. Interestingly, under the CAM, 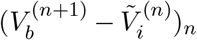 is indeed a Markov chain, making the CAM compatible with the results presented in [51].

Whilst our framework has allowed us to rigorously compare model predictions with data, it has limitations linked to its underlying assumptions. For example, we assume symmetric division, an assumption that has been shown to have strong consequences on the distribution and the correlations of cell variables [19, 26, 32, 29, 52]. We analysed the consequences of relaxing this assumption, using the same definition as in [19, 26, 32, 29] to formalise asymmetric division in our framework (Mathematical Supplementary). Using datasets for which we had precise division information for enough lineages, we showed that this has minimal impact on the estimation of the distributions, with no changes in the ranking of the models based on the BIC and AIC (Table 1 and Figure 3 of Mathematical Supplementary). Another underlying assumption of the framework concerns the noise structure. So far, the noise structure of the control variables has been implicitly assumed to be Gaussian, which we relaxed by assuming a Generalized Gamma distribution. Although this distribution encompasses the broadest family of unimodal distributions of positive random variables, we acknowledge that the underlying molecular mechanisms are still not fully understood. Future work might provide more precise mechanistic information on the distribution of control variables, thus constraining the family of distributions these variables can admit. The statistical method we introduced could be extended in the future by incorporating other covariates into the conditional distribution of division size, such as the size at the end of replication, or by constructing a conditional distribution of the constriction size, since this variable has been shown to be a key factor in cell division [31, 53]. Furthermore, this framework could be expanded to include additional variables of interest, such as the concentration of a control protein and other continuous time-dependent processes, as proposed in [53]. Such an improvement of the framework would enable comparison of mechanistic models and refine our understanding of the coordination of replication-division cycles in bacteria.

## Materials and methods

### The data

In this study the statistical analysis was conducted over the datasets collected by [18] found at https://www.cell.com/current-biology/fulltext/S0960-98221930491-9#mmc3, and by [19] found at https://github.com/guiwitz/DoubleAdderArticle/blob/master/Data_export/Fig1_2_3.csv and by [30] found at https://data.mendeley.com/datasets/hwzcywscc4/draft?a=d486eddf-c4c6-40ee-b262-913f79e5d47b.

From each dataset, four variables have been extracted: the size at division 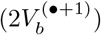, the size at birth 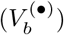, the size per origin right after the initiation of replication at the beginning of the C+D period 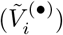, and the C+D period duration 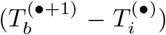. From these variables, the elongation rate over the C+D period (Λ^(*•*)^) was computed by log 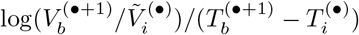.

### Maximization of the likelihoods

The likelihoods defined by Equation (11) of the DIAM and RDAM have been maximised using the Nelder Mead method implemented in the package *optimize* of SciPy in Python [54]. To maximise the likelihoods of the CPM and the CAM, an EM algorithm was implemented (see Mathematical Supplementary). The maximisation steps were performed, using Nelder Mead method implemented in the package *optimize* of SciPy and CMA-ES algorithm implemented in the package *cma* in Python [55, 56]. The likelihood maximization was performed 15 times with different initial points to identify the global maximum.

### Wasserstein distances computation

To compute the empirical Wasserstein distances used in this study, we proceeded as follows: for a given model *M* (chosen among DIAM, RDAM, CPM or CAM), we simulated the division sizes 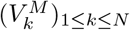, conditionally to the triplet 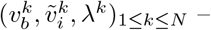 corresponding to an observation of 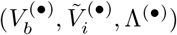. Then we computed the Wasserstein distance between the experimental joint distribution (*v*_*k*_, *x*_*k*_)_*k*_ and its simulated counterpart 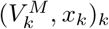 where (*x*_*k*_)_*k*_, denotes either 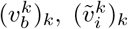 or (*λ*^*k*^)_*k*_. By using the Python package POT [57] we computed the following Wasserstein distance:

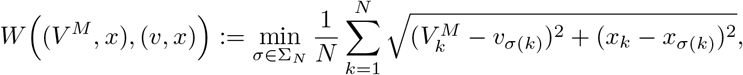

where Σ_*N*_ represents all possible permutations over a *N* -uplet. The parameters used in the simulations were previously estimated from the data by maximizing the likelihood defined by (11). For each dataset and each model *M*, we computed the Wasserstein distances for 100 simulations. From these, we compared the Wasserstein distances of the models using the Wilcoxon-Mann-Whitney test. By denoting 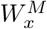 the Wasserstein distance between 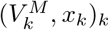 and (*v*_*k*_, *x*_*k*_)_*k*_ for one simulation, according to the model *M*, we

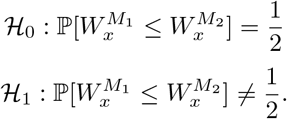

### Estimation of the one-origin-cell division probability in the DIAM

For each dataset, the empirical distributions of Δ_*d*_ and Δ_*i*_ have been regularized through Gaussian kernel density estimation with the package *stats* from SciPy of Python. Then, *L* = 10^6^ cell lineages have been simulated over 100 generations. The final collection of volumes at birth 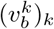 and volumes per origin at initiation 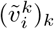 have been used to compute the probability by counting the number of couples that satisfy 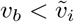 :

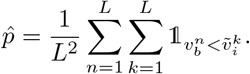

Since 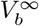 and 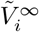 are independent, we can use every couple (*n, k*) *∈ {*1, *· · ·, L}*^2^ to estimate the probability 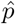.

### Estimation of the probability of a daughter cell dividing before its mother in the RDAM

The probability that a daughter cell divides before its mother in the RDAM has been computed from the experimental measurements of *N* realisations of Δ_*i*_ and Δ_*id*_, noted 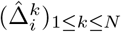 and 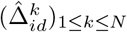 respectively, by counting the number of times the inequality 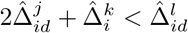 is true for any 1 *≤ j, k, l ≤ N* . This gives the following empirical probability:

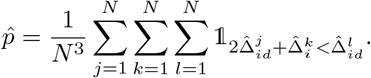

### Simulation of the models under oscillating Δ_*i*_

For this simulation, we generated birth and initiation volumes of 100 lineages, initializing each with 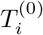 distributed uniformly over [*− τ*, 0], where *τ* denotes the average generation time. To closely replicate the conditions of the DnaA underexpression experiment conducted by [18], we simulated the models using parameters inferred from the MG1655 Glucose dataset. The timeline was divided into intervals of normal expression and intervals of DnaA underexpression. Assuming Δ_*i*_ follows a Generalized Gamma distribution with parameters *a priori* estimated from the same dataset, we modeled the effect of DnaA underexpression by multiplying 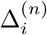 by a factor *f >* 1, when 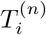 falls in an interval with DnaA underexpression. As in the original experiment, the perturbation and non-perturbation intervals follow one another and each last 2*τ*, resulting in an oscillatory pattern with a period of 4*τ* . In Figure 6, the simulation was performed using a scaling factor *f* = 1, 75.

## Supporting information

Mathematical Supplementary

## Code availability

The codes used in this work can be found at https://github.com/Alex-PERRIN/Coordination-division-replication-E.coli.

## Acknowledgments

The authors acknowledge Luce Breuil, Oskar Girardin, and Valentin Pesce for their fruitful discussions. The authors also acknowledge the reviewers for their very constructive remarks that helped us improve the manuscript. This work has been supported by the Chair “Modélisation Mathématique et Biodiversité” of Veolia Environnement - Ecole Polytechnique - Museum National d’Histoire Naturelle - Fondation X. Funded by the European Union (ERC, SINGER, 101054787). Views and opinions expressed are those of the author(s) only and do not necessarily reflect those of the European Union or the European Research Council. Neither the European Union nor the granting authority can be held responsible for them.

## Supporting information (2 files)

1. **Mathematical Supplementary**
2. **Supplementary Figures**
  - Table S1: **Estimation of the probability of** 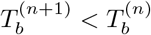 **in the RDAM**.
  - Table S2: **P-values of Shapiro–Wilk normality tests of cell quantities**.
  - Table S3: **BIC and AIC scores**
  - Figure S1: **Simulated cells with less than one origin of replication according to the DIAM**
  - Figure S2: **Illustration of the discrepancy between observed and latent variables in the CPM and the CAM**
  - Figure S3: **Fitting Generalized Gamma distributions to inter-division added size, C+D period, and added size per origin over the C+D period**
  - Figure S4: **Theoretical and empirical distributions of the size at division**
  - Figure S5: **Inferred distributions of the latent variables of the CPM and the CAM**
  - Figure S6: **Probability of having replication limiting in division in the CPM and CAM**
  - Figure S7: **Comparison of the performances of the models based on the AIC**
  - Figure S8: **Wasserstein distances between experimental and simulated joint distributions of division and birth sizes**
  - Figure S9: **Comparison of Wasserstein distances between DIAM/RDAM and CPM**
  - Figure S10: **Comparison of Wasserstein distances between DIAM/RDAM and CAM**
  - Figure S11: **Simulation of the DIAM and RDAM under oscillating distribution of** Δ_*i*_

