## Supplementary material for "Deciphering the Replication-Division Coordination in *E. coli* : A Unified Mathematical Framework for Systematic Model Comparison": Mathematical Supplementary

This supplementary document provides every mathematical proof on which the article relies. In this work, robust mathematical results are shown in the context of a defined framework. This framework reconstructs a survey of a cell lineage in a Mother Machine device. At each generation, a mother cell is randomly replaced by one of its two daughters. This framework relies on four assumptions, stated in the Main Text, and that we recall here.

**Assumptions:**

1. The division is perfectly symmetric, i.e. the volume of the daughter cell at birth is equal to half of the volume of the mother at division.
2. At each replication initiation event, every origin of replication in the cell fires a new round of replication at the same time, thus doubling their number.
3. The origins of replication of a mother cell are equally partitioned into the daughters at division.
4. Over the lineage, the cell volume grows exponentially according to a growth rate function of time $\lambda(t)$, that is positive and locally integrable.

In the following, we study four observable quantities (see Figure 1. A):

- $(V_b^{(n)})_{n\geq 0}$ the sequence of birth volumes where for all $n$, $V_b^{(n)}$ represents the birth volume of the mother cell giving rise to a daughter cell with a birth volume $V_b^{(n+1)}$.
- $(T_b^{(n)})_{n\geq 0}$ the dates of division giving birth to the cell of volume $V_b^{(n)}$
- $(\tilde{V}_i^{(n)})_{n\geq 0}$ the sequence of volumes per origin of replication measured immediately after each initiation event. For every $n$, the initiation corresponding to $\tilde{V}_i^{(n+1)}$ is the one that occurs directly after the initiation associated with $\tilde{V}_i^{(n)}$
- $(T_i^{(n)})_{n\geq 0}$, the date at initiation of replication of the cell with a volume per origin $\tilde{V}_i^{(n)}$

The indexation of these sequences is set as such the sister chromosomes obtained through the replication initiated at $T_i^{(n)}$, are separated at the division occurring at $T_b^{(n+1)}$. This defines the interval $T_b^{(n+1)} - T_i^{(n)}$ as a C+D period, generically referred in the literature. From an experimental point of view the initialization of these sequences can be made as follow (see Figure 1. B): we start with a cell initiating a round of replication setting $T_i^{(0)} = 0$, and $\tilde{V}_i^{(0)}$. Then, the division occurring at the end of the C+D period initiating at $T_i^{(0)}$ sets $T_b^{(1)}$ and $V_b^{(1)}$. Finally, the birth of the cell dividing at $T_b^{(1)}$ sets $T_b^{(0)}$ and $V_b^{(0)}$. In our framework, we choose $\tilde{V}_i^{(0)} > 0$ and $V_b^{(0)} > 0$ to be free variables whose definitions can thus be set arbitrarily. However, we will impose on the initialisation of the sequences that

$$V_b^{(0)} = 2\tilde{V}_i^{(0)} \exp\left(\int_{T_i^{(0)}}^{T_b^{(0)}} \lambda(s) ds\right). \tag{1}$$

This formula enables us to mathematically define the C+D period, which definition is not straightforward. Indeed, the C+D period is often described in the literature as the concatenation of two periods: the C period, which corresponds to the duration of chromosome replication, and the D period, which corresponds to the duration between the termination of replication and the next division. However, the D period is more accurately described as the time between replication termination and division, at which the resulting sister chromosomes are separated. For example, one can imagine a cell with two chromosomes that are replicating into four copies of chromsomes which terminate through the cell cycle. According to the common description in the literature, the D period should correspond to the time between that termination of replication and the division of the cell. However, in fact, the D period spans until the division of the daughter cell, at which the sister chromosomes that terminated replicating are split. Formula (1) captures this phenomenon, which propagates by induction throughout the lineage (see Lemma 1). As a final remark, another variable could have been defined to complete the framework: the volume per termination region at the time of replication termination. This could be defined in a similar way to the sequence of initiation volume per origin and would benefit from a similar initialisation to that in Equation (1). As no data on this quantity has been collected, we did not introduce such a variable into the framework. However, in the context of future work investigating the role of replication termination in controlling division, such a variable could exhibit similar properties to those of the initiation volume per origin and provide a basis for assessing models involving replication termination in division.

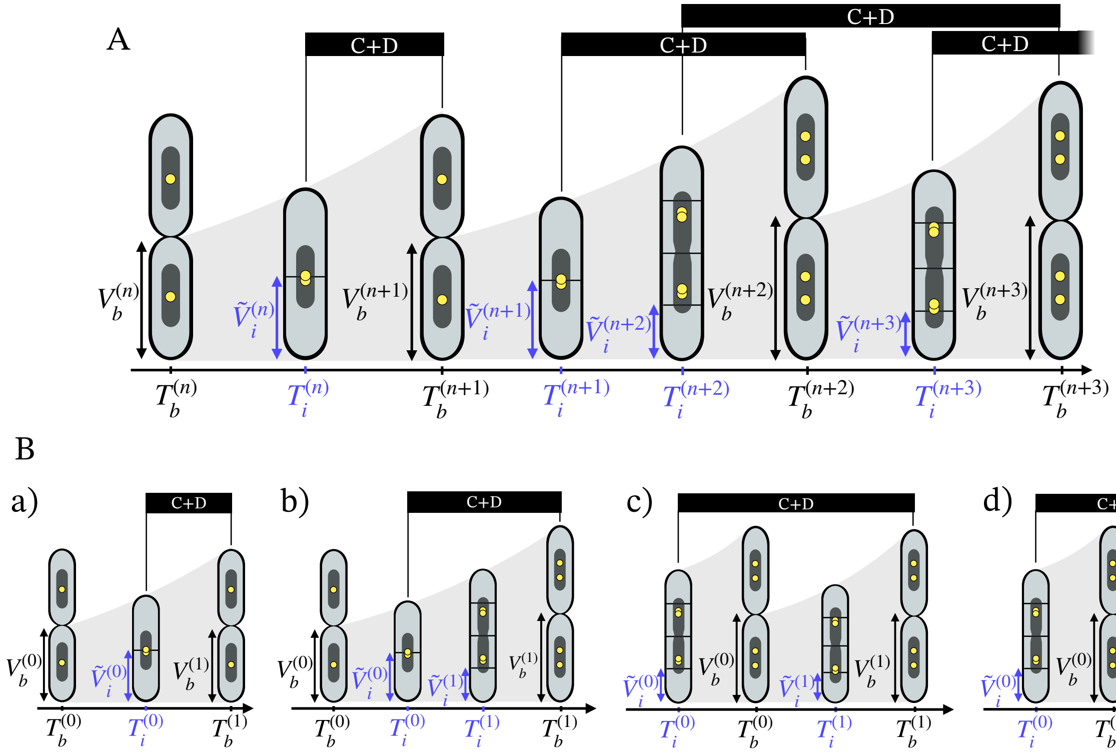


Figure 1: **The framework** We consider a cell lineage where one daughter is kept at each division. Two types of events are monitored: division events and the initiation events. The division events are modeled by the sequences $(T_b^{(n)})_{n\geq 0}$, the dates at division events, and $(V_b^{(n)})_{n\geq 0}$ the birth volume at these events. The initiation of replication events are modeled by the sequences $(T_i^{(n)})_{n\geq 0}$, the dates at initiation of replication events, and $(\tilde{V}_i^{(n)})_{n\geq 0}$ the volume per origin of replication right after these events. The order of occurrence of these events is not defined by the framework. In consequence infinitely many scenarios may set the order of occurrence of these events. (A) Illustration of a cell lineage undergoing an over-initiation, switching from a slow growth regime to a fast growth regime. (B) Illustration of possible scenarios for the initialization of the sequences. Scenario a) depicts a classical cell cycle in a slow growth regime. Scenario b) depicts an over initiation within a slow growing cell. Scenario c) depicts a classical cell cycle in a fast growth regime. Scenario d) depicts an over division of a fast growing cell. In addition to these four scenarios, the framework allows for infinite other possibilities.

### 1 First Mathematical expressions

#### 1.1 Relationship between $(V_b^{(n)})_{n\geq 0}$ and $(T_b^{(n)})_{n\geq 0}$

According to Assumptions 1 and 4 one may find the following recurrent expression for the volume at birth $(V_b^{(n)})_{n\geq 0}$:

$$\forall n \geq 0, \qquad V_b^{(n+1)} = \frac{V_b^{(n)}}{2} \exp\left( \int_{T_b^{(n)}}^{T_b^{(n+1)}} \lambda(s) ds \right). \tag{2}$$

From this recurrent expression, one can derive by induction the following expression that will be useful in the hereafter work:

$$\forall n \geq 1, \qquad V_b^{(n)} = \frac{V_b^{(1)}}{2^{n-1}} \exp\left( \int_{T_b^{(1)}}^{T_b^{(n)}} \lambda(s) ds \right). \tag{3}$$

#### 1.2 Relationship between $(\tilde{V}_i^{(n)})_{n\geq 0}$ and $(T_i^{(n)})_{n\geq 0}$

Similarly to the volume at birth, one may study the trajectory of the volume per origin of replication, observed at replication initiation. According to Assumptions 1 and 3, the cell volume per origin trajectory does not show discontinuity at division events. However, it shows discontinuity at replication initiation events as it is halved by two according to Assumption 2. According to Assumptions 1, 2, 3 and 4 one may find that the sequence of volume per origin at replication initiation follows the recurrence equation

$$\forall n \geq 0, \qquad \tilde{V}_i^{(n+1)} = \frac{\tilde{V}_i^{(n)}}{2} \exp\left( \int_{T_i^{(n)}}^{T_i^{(n+1)}} \lambda(s) ds \right). \tag{4}$$

This gives us by induction:

$$\forall n \geq 0, \qquad \tilde{V}_i^{(n)} = \frac{\tilde{V}_i^{(0)}}{2^{n}} \exp\left( \int_{T_i^{(0)}}^{T_i^{(n)}} \lambda(s) ds \right). \tag{5}$$

#### 1.3 General formula for replication initiation-division coupling

**Lemma 1.** *Assuming that the sequences of birth and initiation volumes and dates are initialized in such a way that* $V_b^{(0)} = 2\tilde{V}_i^{(0)} \exp\big( \int_{T_i^{(0)}}^{T_b^{(0)}} \lambda(s) ds \big)$*, one has* $\forall n \geq 0$*:*

$$V_b^{(n+1)} = \tilde{V}_i^{(n)} \exp\left( \int_{T_i^{(n)}}^{T_b^{(n+1)}} \lambda(s) ds \right). \tag{6}$$

*Proof.* From Equations (3) and (5), one has:

$$
\begin{aligned}
& \frac{V_b^{(n+1)}}{\tilde{V}_i^{(n)}} = \frac{V_b^{(1)}}{\tilde{V}_i^{(0)}} \exp\left( \int_{T_b^{(1)}}^{T_b^{(n+1)}} \lambda(s)ds - \int_{T_i^{(0)}}^{T_i^{(n)}} \lambda(s)ds \right) \\
\iff \quad & \frac{V_b^{(n+1)}}{V_b^{(1)}} = \frac{\tilde{V}_i^{(n)}}{\tilde{V}_i^{(0)}} \exp\left( \int_{T_i^{(n)}}^{T_b^{(n+1)}} \lambda(s)ds + \int_{T_b^{(1)}}^{T_i^{(n)}} \lambda(s)ds - \int_{T_b^{(1)}}^{T_i^{(n)}} \lambda(s)ds - \int_{T_i^{(0)}}^{T_b^{(1)}} \lambda(s)ds \right) \\
\iff \quad & \frac{V_b^{(n+1)}}{V_b^{(1)}} = \frac{\tilde{V}_i^{(n)} \exp\left( \int_{T_i^{(n)}}^{T_b^{(n+1)}} \lambda(s)ds \right)}{\tilde{V}_i^{(0)} \exp\left( \int_{T_i^{(0)}}^{T_b^{(1)}} \lambda(s)ds \right)}.
\end{aligned}
$$

It follows that there exists a $K > 0$ such that for all $n \geq 0$:

$$
V_b^{(n+1)} = K \tilde{V}_i^{(n)} \exp\left( \int_{T_i^{(n)}}^{T_b^{(n+1)}} \lambda(s)ds \right).
$$

By taking the equality form Equation (1) $2V_b^{(1)} = V_b^{(0)} \exp\big( \int_{T_b^{(0)}}^{T_b^{(1)}} \lambda(s)ds \big)$ and the initialization assumption $V_b^{(0)} = 2\tilde{V}_i^{(0)} \exp\big( \int_{T_i^{(0)}}^{T_b^{(0)}} \lambda(s)ds \big)$, it follows that $K = 1$. □

#### 1.4 The number of origins of replication at birth

**Lemma 2.** *Let $\omega^{(n)}$ be the number of origins of replication at birth at $T_b^{(n)}$. Given that $(T_i^{(n)})_{n \geq 0}$ is non-decreasing, for every $n \geq 0$ and $k \geq -n+1$ :*

$$
\begin{aligned}
\omega^{(n)} = 2^k \quad & \iff \quad T_i^{(n+k-1)} \leq T_b^{(n)} < T_i^{(n+k)}, \\
\omega^{(n)} \leq 2^k \quad & \iff \quad T_b^{(n)} < T_i^{(n+k)}.
\end{aligned}
$$

*Proof.* We first introduce two new quantities: the number of initiation events occurred between $T_i^{(0)} = 0$ and time $t > 0$, $N_i^t := \#\{m \geq 0 : \ 0 < T_i^{(m)} \leq t\}$, and the volume per origin over time defined for $t > 0$ by $\tilde{V}_t := 2^{-N_i^t} \tilde{V}_i^{(0)} \exp\left( \int_0^t \lambda(s)ds \right)$. Note that this equation when evaluated at each date of replication initiation given by the sequence $(T_i^{(n)})_{n \geq 0}$, provides the same results as Equation (5). One may notice that the number of origins of replication at the $n$-th birth is precisely the ratio between the volume at the $n$-th birth and the volume per origin at that event. For every $n \geq 1$ and $k \geq -n+1$, one has:

$$
\begin{aligned}
& \omega^{(n)} = 2^k \\
\iff \quad & V_b^{(n)} = 2^k \tilde{V}_{T_b^{(n)}} \\
\iff \quad & \tilde{V}_i^{(n-1)} \exp\left( \int_{T_i^{(n-1)}}^{T_b^{(n)}} \lambda(s)ds \right) = 2^{k - N_i^{T_b^{(n)}}} \tilde{V}_i^{(0)} \exp\left( \int_0^{T_b^{(n)}} \lambda(s)ds \right) \quad \text{by (6)}, \\
\iff \quad & 2^{-n+1} \tilde{V}_i^{(0)} \exp\left( \int_0^{T_b^{(n)}} \lambda(s)ds \right) = 2^{k - N_i^{T_b^{(n)}}} \tilde{V}_i^{(0)} \exp\left( \int_0^{T_b^{(n)}} \lambda(s)ds \right) \quad \text{by (5)}, \\
\iff \quad & N_i^{T_b^{(n)}} = n + k - 1 \\
\iff \quad & T_i^{(n+k-1)} \leq T_b^{(n)} < T_i^{(n+k)} \quad \text{since } (T_i^{(n)})_n \text{ is increasing.}
\end{aligned}
$$

Similarly, one may find through the same operations that for every $n \geq 1$ and $k \geq -n+1$,

$$\omega^{(n)} \leq 2^k \iff N_i^{T_b^{(n)}} \leq n+k-1 \iff T_b^{(n)} < T_i^{(n+k)}.$$

□

### 2 Long-time behavior of the DIAM

According to the Double Independent Adders Model (DIAM) proposed by [S21], the sequences of volume at birth and volume per origin at replication initiation follow for all $n \geq 0$:

$$2V_b^{(n+1)} = V_b^{(n)} + \Delta_d^{(n)} \tag{7}$$

$$2\tilde{V}_i^{(n+1)} = \tilde{V}_i^{(n)} + \Delta_i^{(n)} \tag{8}$$

with $(\Delta_d^{(n)})_{n\geq 0}$ and $(\Delta_i^{(n)})_{n\geq 0}$ being i.i.d. random variables of density $f_{\Delta_d}$ and $f_{\Delta_i}$ respectively. The definition of $\tilde{V}_i^{(0)}$ and $V_b^{(0)}$ can be set arbitrarily, so we will assume that these random variables are independent.

**Remark:** Since $\Delta_d$ and $\Delta_i$ admit densities it turns out that $V_b^{(1)}$ and $\tilde{V}_i^{(1)}$ admit densities for any $V_b^{(0)} > 0$ and $\tilde{V}_i^{(0)} > 0$. Then, by induction for all $n \geq 2$, $V_b^{(n)}$ and $\tilde{V}_i^{(n)}$ are expressed as linear combinations of two independent random variables which both admit densities, thereby ensuring that $V_b^{(n)}$ and $\tilde{V}_i^{(n)}$ themselves admit densities. Furthermore, it is immediate that $V_b^{(n)}$ and $\tilde{V}_i^{(n)}$ are positive for every $n$.

#### 2.1 Convergence of $(V_b^{(n)})_{n\geq 0}$ and $(\tilde{V}_i^{(n)})_{n\geq 0}$

One of the most notable findings in the field of single-cell measurements is the narrow distribution of cell sizes and its stability across experiments. This phenomenon testifies a cell size control which has motivated numerous researches, notably involving partial differential equations [S8, S20, S7]. So far, this size homeostasis is attributed to the division adder model, which has first explained the convergence of the mean of cell size to a steady state [S22, S2, S4]. However fewer works have been led to show the convergence of the distribution of cell size at birth. Gabriel and Martin (2018) [S10] demonstrated that, under the division adder model, the distribution of cell size at birth is the solution to a fixed-point problem. More recently, Madrid Canales (2023) showed that the division adder allowing asymmetric division leads to a stationary distribution for birth size that admits a density [S14].

Here, we extend these findings by showing that even with symmetric division, the volume at birth converges in distribution to a random variable which admits a density, in the condition that the added volume has a density and a finite first moment. While [S10] employed operator theory and [S14] used measure-valued processes, our approach addresses this problem within the constraints of our framework by modeling the sequence of birth volumes as a homogeneous Markov chain.

**Proposition 1.** *According to the DIAM, assuming that $\Delta_d$ is drawn from a density $f_{\Delta_d}$, and such that $\mathbb{E}[\Delta_d] < \infty$, the sequence $(V_b^{(n)})_{n\geq 0}$ converges in law towards a random variable $V_b^{\infty}$*

*which admits a density $f_{V_b^\infty}$.*

*Proof.* Let us first show that $(V_b^{(n)})_n$ converges in law.
Let $\phi_{V_b^{(n)}}$ and $\phi_{\Delta_d}$ be the characteristic functions of $V_b^{(n)}$ and $\Delta_d$ defined for every $t \in \mathbb{R}$, as $\phi_{V_b^{(n)}}(t) := \mathbb{E}[e^{itV_b^{(n)}}]$, and $\phi_{\Delta_d}(t) := \mathbb{E}[e^{it\Delta_d}]$, we have by Equation (7) :

$$\begin{aligned}\forall n \geq 1, \qquad \phi_{V_b^{(n)}}(\xi) &= \phi_{V_b^{(n-1)}}(\xi/2)\phi_{\Delta_d}(\xi/2) \\ &= \phi_{V_b^{(0)}}(\xi/2^n)\prod_{k=1}^{n}\phi_{\Delta_d}(\xi/2^k).\end{aligned}$$

As the characteristic function $\phi_{V_b^{(0)}}$ is continuous and equal to 1 at 0, $\phi_{V_b^{(0)}}(\xi/2^n)$ converges to 1 as $n$ goes to infinity. Now we prove that the right hand term converges and is continuous at 0. For this it suffices to show that the series $\sum_{k\geq 1}|\phi_{\Delta_d}(\xi/2^k) - 1|$ is finite for all $\xi$ (see [S1], Chapter 5: 2.2, Theorem 6), then that $\sum_{k\geq 1}\log(\phi_{\Delta_d}(\xi/2^k))$ is continuous in a neighborhood of zero.

Since we have $\mathbb{E}[\Delta_d] < \infty$, it follows that $\phi_{\Delta_d}$ is differentiable. By applying the Taylor-Young development at 0 to the first order, one ends up for any $t \in \mathbb{R}$:

$$\begin{aligned} & \phi_{\Delta_d}(t) = \phi_{\Delta_d}(t)\Big|_{t=0} + i\mathbb{E}[\Delta_d e^{it\Delta_d}]\Big|_{t=0} t + o(t) \\ \iff \quad & \phi_{\Delta_d}(t) - 1 = i\mathbb{E}[\Delta_d]t + o(t).\end{aligned}$$

The definition of Landau's notation provides us:

$$\forall \epsilon > 0,\ \exists \eta > 0,\ \text{st}\ \forall x \in ]-\eta, \eta[,\ |o(x)| \leq \epsilon|x|.$$

By considering $\epsilon = 1$, there exists $\eta_1 > 0$ such that $\forall x \in ]-\eta_1, \eta_1[$, $|o(x)| \leq |x|$. Since for any $\xi \in \mathbb{R}$ there exists $k_\xi \geq 1$ such that $]-2^{-k_\xi}|\xi|, 2^{-k_\xi}|\xi|[\ \subseteq\ ]-\eta_1, \eta_1[$, it follows that

$$\forall x \in ]-|\xi|2^{-k_\xi},\ |\xi|2^{-k_\xi}\ [,\ |o(x)| \leq |x|.$$

Since $\forall k > k_\xi$, $\xi 2^{-k} \in\ ]-|\xi|2^{-k_\xi},\ |\xi|2^{-k_\xi}\ [$, one has that $\sum_{k>k_\xi}|o(\xi/2^k)| \leq \sum_{k>k_\xi}|\xi/2^k| = 2^{-k_\xi}|\xi|$. Thus, it follows that the series $\sum_{k\geq 1}|\phi_{\Delta_d}(\xi/2^k) - 1|$ is finite for any $\xi \in \mathbb{R}$:

$$\begin{aligned}\sum_{k\geq 1}|\phi_{\Delta_d}(\xi/2^k) - 1| &\leq i\mathbb{E}[\Delta_d]|\xi| + \sum_{k\geq 1}|o(\xi/2^k)| \\ &\leq i\mathbb{E}[\Delta_d]|\xi| + \sum_{k=1}^{k_\xi}|o(\xi/2^k)| + 2^{-k_\xi}|\xi| < \infty.\end{aligned}$$

Then one can show the uniform convergence of $(\sum_{k=1}^{n}\log(\phi_{\Delta_d}(\xi/2^k)))_n$ in a neighborhood of 0. First, one may proceed to a Taylor-Young development of $\log(\phi_{\Delta_d})$ in a neighborhood of zero, in which case the principal value of the complex logarithm is well defined since $\phi_{\Delta_d}$ evaluated on such neighborhood is defined on a neighborhood of one:

$$\begin{aligned}&\log(\phi_{\Delta_d}(t)) = \log(\phi_{\Delta_d}(t))\Big|_{t=0} + \frac{i\mathbb{E}[\Delta_d e^{it\Delta_d}]}{\phi_{\Delta_d}(t)}\Bigg|_{t=0} t + o(t) \\ &\log(\phi_{\Delta_d}(t)) = i\mathbb{E}[\Delta_d]t + o(t).\end{aligned}$$

Then, one can notice that for $\delta > 0$ fixed, $\exists\, k_\delta \geq 1$ such that $\forall \xi \in ]-\delta,\delta[,\ 2^{-k_\delta}|\xi| \leq 2^{-k_\delta}\delta \leq \eta_1$. As previously, it follows that,

$$\exists\, k_\delta \geq 1,\ \forall \xi \in ]-\delta,\delta[,\ \forall n \geq k_\delta, \quad \sum_{k>n} |o(\xi/2^k)| \leq \sum_{k>n} |\xi/2^k| = 2^{-n}|\xi| \leq 2^{-n}\delta.$$

Thus for $\delta > 0$ fixed, $\exists\, k_\delta \geq 1$, such that $\forall \xi \in ]-\delta,\delta[,\ \forall n \geq k_\delta$,

$$\begin{aligned}\left|\sum_{k>n} \log(\phi_{\Delta_d}(\xi/2^k))\right| &\leq \sum_{k>n} \left|\log(\phi_{\Delta_d}(\xi/2^k))\right| \\ &\leq 2^{-n}\mathbb{E}[\Delta_d]|\xi| + \sum_{k>n} |o(\xi/2^k)| \\ &\leq 2^{-n}\delta\left(\mathbb{E}[\Delta_d]+1\right).\end{aligned}$$

Thus, for $\delta > 0$ fixed, $\exists\, k_\delta \geq 1$, such that $\forall \epsilon > 0$, $\exists\, N \geq k_\delta$ such that $\forall n \geq N$, $2^{-n}\delta\left(\mathbb{E}[\Delta_d]+1\right) \leq \epsilon$. This concludes that the series uniformly converges in a neighbourhood of 0. Since there exists a neighbourhood of zero where $\phi_{\Delta_d}$ is continuous, it follows that the series $\sum_{k\geq 1} \log(\phi_{\Delta_d}(\xi/2^k))$ is continuous on that neighbourhood, and therefore at zero.

Thus, by Levy's theorem (see [S9] Chapter XV: 3, Theorem 2) since $\phi_{V_b^{(n)}}$ converges pointwise to a complex function continuous at zero, $V_b^{(n)}$ converges in law to a random variable $V_b^\infty$.

Now we prove that the random variable $V_b^\infty$ has a density. To this end, the proof is inspired by the approach presented in Proposition 1 of [S17].
On one hand, one has for all $n \geq 1$ and for every continuous and bounded function $g$:

$$\mathbb{E}[g(V_b^{(n+1)})] = \mathbb{E}[\mathbb{E}[g(V_b^{(n+1)})|V_b^{(n)}]] \qquad (*)$$

with

$$\mathbb{E}[g(V_b^{(n+1)})|V_b^{(n)} = y] = \int_0^{+\infty} g\left(\frac{x+y}{2}\right) f_{\Delta_d}(x)dx.$$

Since $y \mapsto \mathbb{E}[g(V_b^{(n+1)})|V_b^{(n)} = y]$ is continuous and bounded for any continuous and bounded function $g$, it follows by the definition of the weak convergence (see [S3] Chapter 1: 4.) that

$$\lim_{n\to\infty} \mathbb{E}[\mathbb{E}[g(V_b^{(n+1)})|V_b^{(n)}]] = \int_0^{+\infty}\int_0^{+\infty} g\left(\frac{x+y}{2}\right) f_{\Delta_d}(x)dx\ \pi(dy),$$

with $\pi$ the distribution of $V_b^\infty$. Then, one can get through a change of variable and the application of Fubini's theorem:

$$\begin{aligned}\int_0^{+\infty}\int_0^{+\infty} g\left(\frac{x+y}{2}\right) f_{\Delta_d}(x)dx\ \pi(dy) &= 2\int_0^{+\infty}\int_{\frac{y}{2}}^{+\infty} g(z) f_{\Delta_d}(2z-y)dz\ \pi(dy) \\ &= 2\int_0^{+\infty} g(z)\int_0^{2z} f_{\Delta_d}(2z-y)\pi(dy)\ dz.\end{aligned}$$

On another hand, one has from the definition of the convergence in law of a random variable that for every continuous and bounded function $g$:

$$\begin{aligned}\lim_{n\to\infty} \mathbb{E}[g(V_b^{(n+1)})] &= \mathbb{E}[g(V_b^\infty)] \\ &= \int_0^{+\infty} g(z)\pi(dz).\end{aligned}$$

Thus from the equality $(*)$, by passing through the limit, one can notice that the distribution of $V_b^\infty$ is absolutely continuous with respect to the Lebesgue measure and thus admits a density.

□

Supposing that $\mathbb{E}[\Delta_i] < \infty$, with the same proof, one can show that the sequence of volumes per origin at replication initiation $(\tilde{V}_i^{(n)})_{n\in\mathbb{N}}$ converges in law to a a random variable $\tilde{V}_i^\infty$.

**Proposition 2.** *According to the initiation adder, assuming that $\Delta_i$ is drawn from a density $f_{\Delta_i}$, and such that $\mathbb{E}[\Delta_i] < \infty$, the sequence $(\tilde{V}_i^{(n)})_{n\in\mathbb{N}}$ converges in law to a random variable $\tilde{V}_i^\infty$ which admits a density $f_{\tilde{V}_i^\infty}$.*

#### 2.2 Convergence of the number of origins of replication

**Proposition 3.** *According to the DIAM, assuming that $V_b^{(0)}$ and $\tilde{V}_i^{(0)}$ are independent, the sequence $(\omega^{(n)})_{n\in\mathbb{N}*}$ converges in law towards a random variable $\omega_\infty$ such that $\forall\, k \in \mathbb{Z}$,*

$$\mathbb{P}\left[\log_2(\omega_\infty) \leq k\right] = \mathbb{P}\left[V_b^\infty < 2^{k+1}\tilde{V}_i^\infty\right]. \tag{9}$$

*Proof.* From the result provided by Proposition 1, one has $\forall\, n \geq 1$ and $\forall\, k \geq -n$,

$$\begin{aligned}
\mathbb{P}\left[\log_2(\omega^{(n)}) \leq k\right] &= \mathbb{P}\left[T_b^{(n)} < T_i^{(n+k)}\right] \\
&= \mathbb{P}\left[\exp\left(\int_0^{T_b^{(n)}} \lambda(s)ds\right) < \exp\left(\int_0^{T_i^{(n+k)}} \lambda(s)ds\right)\right] \\
&= \mathbb{P}\left[V_b^{(1)} \exp\left(\int_0^{T_b^{(n)}} \lambda(s)ds\right) < \tilde{V}_i^{(0)} \exp\left(\int_0^{T_b^{(1)}} \lambda(s)ds\right) \exp\left(\int_0^{T_i^{(n+k)}} \lambda(s)ds\right)\right] && \text{by (6),} \\
&= \mathbb{P}\left[V_b^{(1)} \exp\left(\int_{T_b^{(1)}}^{T_b^{(n)}} \lambda(s)ds\right) < \tilde{V}_i^{(0)} \exp\left(\int_0^{T_i^{(n+k)}} \lambda(s)ds\right)\right] \\
&= \mathbb{P}\left[V_b^{(n)} < 2^{k+1}\tilde{V}_i^{(n+k)}\right] && \text{by (3), (5).}
\end{aligned}$$

Since $V_b^{(0)}$ and $\tilde{V}_i^{(0)}$ are independent, the previous probability relates to an inequality between two independent and convergent random variables. It follows for every $k$ fixed that

$$\left(V_b^{(n)}, \tilde{V}_i^{(n+k)}\right) \xrightarrow{D}_{n\to\infty} \left(V_b^\infty, \tilde{V}_i^\infty\right).$$

Then, thanks to the continuous mapping theorem (see [S3] Chapter 1: 5.), we have:

$$V_b^{(n)} - 2^{k+1}\tilde{V}_i^{(n+k)} \xrightarrow{D}_{n\to\infty} V_b^\infty - 2^{k+1}\tilde{V}_i^\infty.$$

Then, since $\tilde{V}_i^\infty$ and $V_b^\infty$ have a density and are independent, we have thanks to the Portemanteau Theorem (see [S3] Chapter 1: 4. Theorem 2.1) that

$$\lim_{n\to\infty} \mathbb{P}\left[V_b^{(n)} - 2^{k+1}\tilde{V}_i^{(n+k)} < 0\right] = \mathbb{P}[V_b^\infty - 2^{k+1}\tilde{V}_i^\infty < 0] =: p_k.$$

Since, $\forall n \in \mathbb{N}^*$ and $\forall k \geq -n$, $\mathbb{P}\left[\log_2(\omega^{(n)}) = k\right] = \mathbb{P}\left[\log_2(\omega^{(n)}) \leq k\right] - \mathbb{P}\left[\log_2(\omega^{(n)}) \leq k-1\right]$, it follows that for every $k \in \mathbb{Z}$

$$\lim_{n\to\infty} \mathbb{P}\left[\log_2(\omega^{(n)}) = k\right] = p_k - p_{k-1}.$$

One can integrate this quantity on $\mathbb{Z}$ as following. For any $n \geq 1$ one has,

$$\begin{aligned}\sum_{-n\leq k\leq n} (p_k - p_{k-1}) &= \sum_{k=0}^{n}(p_k - p_{k-1}) + \sum_{k=1}^{n}(p_{-k} - p_{1-k}) \\ &= p_n - p_{-n}\end{aligned}$$

Since $V_b^\infty$ and $\tilde{V}_i^\infty$ are positive almost surely, it follows that $\lim_{n\to\infty} p_n = \mathbb{P}[\tilde{V}_i^\infty/V_b^\infty > 0] = 1$ and $\lim_{n\to\infty} p_{-n} = \mathbb{P}[V_b^\infty/\tilde{V}_i^\infty \leq 0] = 0$. This concludes that $(\omega^{(n)})_{n\in\mathbb{N}^*}$ converges in law towards a random variable $\omega_\infty$ distributed on $\{2^k : k \in \mathbb{Z}\}$ which has a cumulative probability function given by (9). ☐

### 3 Necessary and sufficient conditions to avoid one-origin-cell division in the DIAM

**Theorem 1.** *The DIAM ensures $\mathbb{P}[\omega_\infty \geq 1] = 1$ if and only if there exists $\sigma > 0$ such that:*

$$Supp(f_{\Delta_i}) \subset [0, \sigma], \qquad and \qquad Supp(f_{\Delta_d}) \subset [\sigma, +\infty[.$$

*Proof.* In order to have a number of origins greater or equal to one almost surely, we aim to deduce necessary and sufficient constraints to impose to $\Delta_i$ and $\Delta_d$, satisfying $\mathbb{P}[\omega_\infty \geq 1] = 1$. From (9), this equality is equivalent to

$$\mathbb{P}\left[\tilde{V}_i^\infty \leq V_b^\infty\right] = 1. \qquad (*)$$

- First we show that $(*)$ is equivalent to

$$\exists \sigma > 0, \quad \mathbb{P}\left[V_b^\infty \geq \sigma\right] = 1 \quad \text{and} \quad \mathbb{P}\left[\tilde{V}_i^\infty \leq \sigma\right] = 1.$$

  - $\Longleftarrow$ It is immediate that if there exists $\sigma > 0$ such that $V_b^\infty \geq \sigma$ almost surely and $\tilde{V}_i^\infty \leq \sigma$ almost surely, then $\tilde{V}_i^\infty \leq \sigma \leq V_b^\infty$ almost surely.
  - $\Longrightarrow$ Now we show that if $\tilde{V}_i^\infty \leq V_b^\infty$ almost surely, then there exists a $\sigma > 0$ such that $\tilde{V}_i^\infty \leq \sigma \leq V_b^\infty$ almost surely. Since $V_b^\infty$ is a positive random variable, one may introduce $\sigma^* := \inf Supp(f_{V_b^\infty}) \geq 0$, with $f_{V_b^\infty}$ the density of $V_b^\infty$.
    Since $V_b^\infty$ and $\tilde{V}_i^\infty$ are independent, it follows for every $\epsilon > 0$,

$$\begin{aligned}\mathbb{P}[V_b^\infty \leq \tilde{V}_i^\infty] &\geq \mathbb{P}[V_b^\infty \leq \sigma^* + \epsilon \leq \tilde{V}_i^\infty] \\ &\geq \mathbb{P}[V_b^\infty \leq \sigma^* + \epsilon]\mathbb{P}[\sigma^* + \epsilon \leq \tilde{V}_i^\infty].\end{aligned}$$

    By definition, $\mathbb{P}[V_b^\infty \leq \sigma^* + \epsilon] > 0$ for every $\epsilon > 0$. Hence, if $\mathbb{P}[V_b^\infty \leq \tilde{V}_i^\infty] = 0$, then $\mathbb{P}[\sigma^* + \epsilon \leq \tilde{V}_i^\infty] = 0$ for every $\epsilon > 0$ and therefore $\tilde{V}_i^\infty \leq \sigma^*$ almost surely. Finally, since $\tilde{V}_i^\infty$ is positive almost surely, one has necessarily $\sigma^* > 0$.

- Now, we show that in order to have $\mathbb{P}\left[V_b^\infty \geq \sigma\right] = 1$, it is necessary and sufficient to have $\mathbb{P}\left[\Delta_d \geq \sigma\right] = 1$.

– $\Longleftarrow$: We suppose that $\mathbb{P}\left[\Delta_d \geq \sigma\right] = 1$. Since $\mathbb{P}[\bigcup_{k\geq 1}\{\Delta_d^{(k)} < \sigma\}] \leq \sum_{k\geq 1}\mathbb{P}[\Delta_d^{(k)} < \sigma] = 0$, it follows that $\mathbb{P}[\bigcap_{k\geq 1}\{\Delta_d^{(k)} \geq \sigma\}] = 1$. Since $\bigcap_{k\geq 1}\{\Delta_d^{(k)} \geq \sigma\} \subseteq \{\sum_{k\geq 1}\frac{\Delta_d^{(k)}}{2^k} \geq \sum_{k\geq 1}\frac{\sigma}{2^k}\}$,

$$\mathbb{P}\left[\sum_{k\geq 1}\frac{\Delta_d^{(k)}}{2^k} \geq \sigma\right] = 1.$$

As $V_b^\infty \overset{D}{=} \sum_{k\geq 1}\frac{\Delta_d^{(k)}}{2^k}$ from Proposition 2, one thus has $\mathbb{P}\left[\sum_{k\geq 1}\frac{\Delta_d^{(k)}}{2^k} \geq \sigma\right] = \mathbb{P}\left[V_b^\infty \geq \sigma\right] = 1$.

– $\Longrightarrow$: By contraposition, we aim to show that $\mathbb{P}\left[\Delta_d < \sigma\right] > 0$ implies that $\mathbb{P}\left[V_b^\infty < \sigma\right] > 0$.
Note that if $\mathbb{P}\left[\Delta_d < \sigma\right] > 0$, then there exists $\epsilon > 0$ such that $\mathbb{P}\left[\Delta_d \leq \sigma - \epsilon\right] > 0$.
One may start by noticing that for any $1 \leq m < M$, and for any $\epsilon > 0$

$$\left\{\sum_{k=1}^{m}\frac{\Delta_d^{(k)}}{2^k} \leq \sigma - \epsilon\right\} \cap \left\{\sum_{k=m+1}^{M}\frac{\Delta_d^{(k)}}{2^k} < \epsilon\right\} \subseteq \left\{\sum_{k=1}^{M}\frac{\Delta_d^{(k)}}{2^k} < \sigma\right\}.$$

Since $(\Delta_d^{(n)})_{n\in\mathbb{N}^*}$ is a sequence of independent random variables, it follows that for any $1 \leq m < M$ and any $\epsilon > 0$,

$$\mathbb{P}\left[\sum_{k=1}^{M}\frac{\Delta_d^{(k)}}{2^k} < \sigma\right] \geq \mathbb{P}\left[\sum_{k=1}^{m}\frac{\Delta_d^{(k)}}{2^k} \leq \sigma - \epsilon\right]\mathbb{P}\left[\sum_{k=m+1}^{M}\frac{\Delta_d^{(k)}}{2^k} < \epsilon\right]. \qquad (**)$$

The aim in the following is to show that the left hand side of $(**)$ is positive when $M \to \infty$ if there exists $\epsilon > 0$ such that $\mathbb{P}\left[\Delta_d \leq \sigma - \epsilon\right] > 0$.

Firstly, we show that the first term of the right-hand side of $(**)$ is positive for any $m \geq 1$ when $\epsilon$ satisfies $\mathbb{P}\left[\Delta_d \leq \sigma - \epsilon\right] > 0$. Indeed, one may notice that

$$\bigcap_{k=1}^{m}\left\{\Delta_d^{(k)} \leq \sigma - \epsilon\right\} \subseteq \left\{\sum_{k=1}^{m}\frac{\Delta_d^{(k)}}{2^k} \leq \sum_{k=1}^{m}\frac{\sigma - \epsilon}{2^k}\right\} \subseteq \left\{\sum_{k=1}^{m}\frac{\Delta_d^{(k)}}{2^k} \leq \sigma - \epsilon\right\}.$$

Since $(\Delta_d^{(n)})_{n\in\mathbb{N}^*}$ is a sequence of independent random variables, if $\epsilon$ satisfies $\mathbb{P}\left[\Delta_d \leq \sigma - \epsilon\right] > 0$, then for any $m \geq 1$

$$\mathbb{P}\left[\sum_{k=1}^{m}\frac{\Delta_d^{(k)}}{2^k} \leq \sigma - \epsilon\right] \geq \mathbb{P}\left[\Delta_d \leq \sigma - \epsilon\right]^m > 0.$$

Secondly, we show that for any $\epsilon > 0$, there exists a $m \geq 1$ such that the second term of the right-hand side of $(**)$ is positive when $M \to \infty$.
First, we have by the Markov inequality for any $\epsilon > 0$ and any $1 \leq m < M$,

$$\mathbb{P}\left[\sum_{k=m+1}^{M}\frac{\Delta_d^{(k)}}{2^k} \geq \epsilon\right] \leq \frac{\mathbb{E}[\sum_{k=m+1}^{M}\frac{\Delta_d^{(k)}}{2^k}]}{\epsilon} = \mathbb{E}[\Delta_d]\frac{1 - \frac{1}{2^{M-m}}}{2^m\epsilon}.$$

It follows that for any $\epsilon > 0$, and any $m \geq 1$

$$\lim_{M\to\infty}\mathbb{P}\left[\sum_{k=m+1}^{M}\frac{\Delta_d^{(k)}}{2^k} \geq \epsilon\right] \leq \frac{\mathbb{E}[\Delta_d]}{2^m\epsilon}.$$

Since $\mathbb{E}[\Delta_d] < \infty$, for any $\epsilon > 0$, there exists $m \geq 1$ such that $\mathbb{E}[\Delta_d]/2^m\epsilon < 1$. Hence, it follows that for any $\epsilon > 0$, there exists $m \geq 1$ such that

$$\lim_{M\to\infty} \mathbb{P}\left[\sum_{k=m+1}^{M} \frac{\Delta_d^{(k)}}{2^k} < \epsilon\right] > 0.$$

In summary, it follows that if there exists any $\epsilon > 0$ satisfying $\mathbb{P}\left[\Delta_d \leq \sigma - \epsilon\right] > 0$, then

$$\exists\, m \geq 1, \quad \lim_{M\to\infty} \left(\mathbb{P}\left[\sum_{k=1}^{m} \frac{\Delta_d^{(k)}}{2^k} \leq \sigma - \epsilon\right] \mathbb{P}\left[\sum_{k=m+1}^{M} \frac{\Delta_d^{(k)}}{2^k} < \epsilon\right]\right) > 0.$$

Regarding the limit of the left hand side of (∗∗), one has that $\sum_{k=1}^{M} \frac{\Delta_d^{(k)}}{2^k}$ converges in law to the random variable $V_b^\infty$ defined in Proposition 2 which admits a density. This convergence in law is equivalent to the convergence of the cumulative distribution functions of $\sum_{k=1}^{M} \frac{\Delta_d^{(k)}}{2^k}$ to the continuous cumulative distribution function of $V_b^\infty$ (see [S9] Chapter VIII: 1. Theorem 1), giving us for any $\sigma$

$$\lim_{M\to\infty} \mathbb{P}\left[\sum_{k=1}^{M} \frac{\Delta_d^{(k)}}{2^k} < \sigma\right] = \mathbb{P}\left[V_b^\infty < \sigma\right].$$

In conclusion, by passing the inequality (∗∗) to the limit when $M \to \infty$, one finally gets

$$\exists\, \epsilon > 0,\ \mathbb{P}\left[\Delta_d \leq \sigma - \epsilon\right] > 0 \implies \mathbb{P}\left[V_b^\infty < \sigma\right] > 0.$$

We thus showed that $\mathbb{P}\left[V_b^\infty \geq \sigma\right] = 1$ implies that $\mathbb{P}\left[\Delta_d \geq \sigma\right] = 1$ and reciprocally, which concludes the equivalence of these two assertions.

- Finally, we show that in order to satisfy $\mathbb{P}\left[\tilde{V}_i^\infty \geq \sigma\right] = 0$, it is necessary and sufficient to have $\mathbb{P}\left[\Delta_i \geq \sigma\right] = 0$.

  - $\Longleftarrow$ As in the previous proof, we assume that $\mathbb{P}\left[\Delta_i < \sigma\right] = 1$. Since $\mathbb{P}[\bigcup_{k\geq 1}\{\Delta_i^{(k)} \geq \sigma\}] \leq \sum_{k\geq 1} \mathbb{P}[\Delta_i^{(k)} \geq \sigma] = 0$ it follows that $\mathbb{P}[\bigcap_{k\geq 1}\{\Delta_i^{(k)} < \sigma\}] = 1$. Since $\bigcap_{k\geq 1}\{\Delta_i^{(k)} < \sigma\} \subseteq \{\sum_{k\geq 1} \frac{\Delta_i^{(k)}}{2^k} < \sum_{k\geq 1} \frac{\sigma}{2^k}\}$, it implies that

    $$\mathbb{P}\left[\sum_{k\geq 1} \frac{\Delta_i^{(k)}}{2^k} < \sigma\right] = 1.$$

    As $\tilde{V}_i^\infty \overset{D}{=} \sum_{k\geq 1} \frac{\Delta_i^{(k)}}{2^k}$, $\mathbb{P}\left[\sum_{k\geq 1} \frac{\Delta_i^{(k)}}{2^k} < \sigma\right] = \mathbb{P}\left[\tilde{V}_i^\infty < \sigma\right] = 1$.

  - $\Longrightarrow$ We now show that $\mathbb{P}[\tilde{V}_i^\infty < \sigma] = 1$ implies $\mathbb{P}[\Delta_i < \sigma] = 1$.

    First, since $\Delta_i$ is positive almost surely, the sequence of events $\left(\left\{\sum_{k=1}^{m} \frac{\Delta_i^{(k)}}{2^k} \geq \sigma\right\}\right)_{m\geq 1}$ is increasing. Thus, one has for any $1 \leq m < M$ that

    $$\left\{\sum_{k=1}^{m} \frac{\Delta_i^{(k)}}{2^k} \geq \sigma\right\} \subseteq \left\{\sum_{k=1}^{M} \frac{\Delta_i^{(k)}}{2^k} \geq \sigma\right\}.$$

Concurrently, one has for every $m \geq 1$ that

$$\bigcap_{k=1}^{m}\left\{\Delta_i^{(k)} \geq \frac{\sigma}{1-\frac{1}{2^m}}\right\} \subseteq \left\{\sum_{k=1}^{m}\frac{\Delta_i^{(k)}}{2^k} \geq \frac{\sigma}{1-\frac{1}{2^m}}\sum_{k=1}^{m}\frac{1}{2^k}\right\} = \left\{\sum_{k=1}^{m}\frac{\Delta_i^{(k)}}{2^k} \geq \sigma\right\}.$$

Since the sequence $(\Delta_i^{(k)})_{k\geq 1}$ is independent, one thus has for every $1 \leq m < M$ that

$$\mathbb{P}\left[\Delta_i \geq \frac{\sigma}{1-\frac{1}{2^m}}\right]^m \leq \mathbb{P}\left[\sum_{k=1}^{M}\frac{\Delta_i^{(k)}}{2^k} \geq \sigma\right].$$

Then, one can study the limit of the right-hand side as $M \to \infty$. As previously, one has that the cumulative distribution function of $\sum_{k=1}^{M}\frac{\Delta_i^{(k)}}{2^k}$ converges to the continuous cumulative distribution function of $\tilde{V}_i^{\infty}$ leading for every $m \geq 1$ to

$$\mathbb{P}\left[\Delta_i \geq \frac{\sigma}{1-\frac{1}{2^m}}\right]^m \leq \mathbb{P}\left[\tilde{V}_i^{\infty} \geq \sigma\right].$$

It finally turns out that if $\mathbb{P}\left[\tilde{V}_i^{\infty} \geq \sigma\right] = 0$, then $\mathbb{P}\left[\Delta_i \geq \frac{\sigma}{1-\frac{1}{2^m}}\right] = 0$ for every $m \geq 1$. Since for every $\epsilon > 0$ there exists $m \geq 1$ such that $\sigma + \epsilon \geq \frac{\sigma}{1-\frac{1}{2^m}}$ implying that $\mathbb{P}\left[\Delta_i \geq \sigma+\epsilon\right] \leq \mathbb{P}\left[\Delta_i \geq \frac{\sigma}{1-\frac{1}{2^m}}\right]$, we conclude that

$$\mathbb{P}\left[\tilde{V}_i^{\infty} \geq \sigma\right] = 0 \implies \forall \epsilon > 0,\ \mathbb{P}\left[\Delta_i \geq \sigma+\epsilon\right] = 0.$$

□

### 4 Necessary and sufficient conditions to prevent a daughter cell from dividing before its mother in the RDAM

The Replication Double Adders Model (RDAM) proposed by [S25] assumes an initiation adder and thus that the volume per origin at replication initiation follows the recurrent equation (8). Then, the RDAM assumes that the sequence of volume at birth satisfies for all $n$

$$V_b^{(n+1)} = \tilde{V}_i^{(n)} + \Delta_{id}^{(n)}, \tag{10}$$

with $(\Delta_{id}^{(n)})_{n\geq 0}$ being a i.i.d. random variables sequence with density $f_{\Delta_{id}}$.

**Remark:** As $\tilde{V}_i^{(n)}$ and $\Delta_{id}^{(n)}$ are independent and admit both densities for all $n \geq 0$ it turns out that $V_b^{(n)}$ admits also a density for all $n \geq 1$. Also, from Proposition 3, it follows that $(V_b^{(n)})_{n\geq 1}$ converges in distribution to a random variable $V_b^{\infty} \overset{D}{=} \tilde{V}_i^{\infty} + \Delta_{id}$.

**Proposition 4.** *In the RDAM, $(T_b^{(n)})_{n\in\mathbb{N}*}$ is almost surely non-decreasing if and only if there exist $(\sigma_1, \sigma_2) \in \mathbb{R}_+^2 \setminus \{(0,0)\}$, such that*

$$Supp(f_{\Delta_{id}}) \subseteq [\sigma_1, 2\sigma_1+\sigma_2] \qquad \textit{and} \qquad Supp(f_{\Delta_i}) \subseteq [\sigma_2, \infty[.$$

*Proof.* We have for all $n \in \mathbb{N}$

$$
\begin{aligned}
& T_b^{(n+1)} \leq T_b^{(n+2)} \quad a.s. & \\
\iff \quad & V_b^{(n+1)} \leq 2V_b^{(n+2)} \quad a.s. & \text{by (3)} \\
\iff \quad & \tilde{V}_i^{(n)} + \Delta_{id}^{(n)} \leq 2(\tilde{V}_i^{(n+1)} + \Delta_{id}^{(n+1)}) \quad a.s. & \text{by (10)} \\
\iff \quad & \Delta_{id}^{(n)} \leq 2\Delta_{id}^{(n+1)} + \Delta_i^{(n)} \quad a.s. & \text{by (8)}
\end{aligned}
$$

We now show that in order to have $\Delta_{id}^{(n)} \leq 2\Delta_{id}^{(n+1)} + \Delta_i^{(n)}$ almost surely, it is necessary and sufficient to have the existence of $(\sigma_1, \sigma_2) \in \mathbb{R}_+^2 \setminus \{(0,0)\}$ such that $\sigma_1 \leq \Delta_{id} \leq 2\sigma_1 + \sigma_2$ almost surely and $\sigma_2 \leq \Delta_i$ almost surely.

- $\Longleftarrow$ Supposing that there exist $(\sigma_1, \sigma_2) \in \mathbb{R}_+^2 \setminus \{(0,0)\}$ such that $\sigma_1 \leq \Delta_{id} \leq 2\sigma_1 + \sigma_2$ and $\sigma_2 \leq \Delta_i$ almost surely, it follows immediately that

$$\Delta_{id}^{(n)} \leq 2\sigma_1 + \sigma_2 \leq 2\Delta_{id}^{(n+1)} + \Delta_i^{(n)} \quad a.s.$$

- $\Longrightarrow$ Since $\Delta_{id}$ and $\Delta_i$ are positive almost surely, there exist $\sigma_1^* := \inf Supp(f_{\Delta_{id}}) \geq 0$ and $\sigma_2^* := \inf Supp(f_{\Delta_i}) \geq 0$. We now want to show that if $\mathbb{P}[\Delta_{id}^{(n)} \leq 2\Delta_{id}^{(n+1)} + \Delta_i^{(n)}] = 1$ therefore we have $\Delta_{id} \leq 2\sigma_1^* + \sigma_2^*$ almost surely.

  We have for every $\epsilon > 0$,

$$
\begin{aligned}
\mathbb{P}[\Delta_{id}^{(n)} > 2\Delta_{id}^{(n+1)} + \Delta_i^{(n)}] &\geq \mathbb{P}[\Delta_{id}^{(n)} > 2\sigma_1^* + \sigma_2^* + \epsilon \geq 2\Delta_{id}^{(n+1)} + \Delta_i^{(n)}] \\
&\geq \mathbb{P}[\Delta_{id}^{(n)} > 2\sigma_1^* + \sigma_2^* + \epsilon]\mathbb{P}[\Delta_{id}^{(n)} \leq \sigma_1^* + \frac{\epsilon}{4}]\mathbb{P}[\Delta_i^{(n)} \leq \sigma_2^* + \frac{\epsilon}{4}].
\end{aligned}
$$

  By definition, we have for every $\epsilon > 0$ that $\mathbb{P}[\Delta_{id}^{(n)} \leq \sigma_1^* + \frac{\epsilon}{4}] > 0$ and $\mathbb{P}[\Delta_i^{(n)} \leq \sigma_2^* + \frac{\epsilon}{4}] > 0$. In consequence, if $\mathbb{P}[\Delta_{id}^{(n)} > 2\Delta_{id}^{(n+1)} + \Delta_i^{(n)}] = 0$, then $\mathbb{P}[\Delta_{id}^{(n)} > 2\sigma_1^* + \sigma_2^* + \epsilon] = 0$ for every $\epsilon > 0$. Since $\Delta_{id}$ is almost surely positive, one necessarily has $(\sigma_1^*, \sigma_2^*) \neq (0,0)$.

In conclusion, in order to have $(T_b^{(n)})_{n \in \mathbb{N}^*}$ non-decreasing almost surely, it is necessary and sufficient to have $(\sigma_1, \sigma_2) \in \mathbb{R}_+^2 \setminus \{(0,0)\}$ such that $\sigma_1 \leq \Delta_{id} \leq 2\sigma_1 + \sigma_2$ almost surely and $\sigma_2 \leq \Delta_i$ almost surely. □

### 5 Conditional densities of the models: expressions, identifiablility and estimation

In this section, we present results for two other models: the Concurrent Processes Model (CPM) proposed by [S18, S19] and the Concurrent Adders Model (CAM) first proposed in this work.

For the CPM, the sequence of volume at birth $(V_b^{(n)})_{n \geq 0}$ follows $\forall\, n \geq 0$ the recurrent expression

$$V_b^{(n+1)} = \max\Big(\tilde{V}_i^{(n)} e^{\Lambda^{(n)} R^{(n)}},\ 2^{-1}(V_b^{(n)} + \Delta_d^{(n)})\Big), \tag{11}$$

with $(\Delta_d^{(n)})_{n \geq 0}$ and $(R^{(n)})_{n \geq 0}$ independent and i.i.d. random variables sequences, and $\Lambda^{(n)} := \frac{1}{R^{(n)}} \int_{T_i^{(n)}}^{T_i^{(n)} + R^{(n)}} \lambda(s) ds$ assumed to be independent from $(R^{(n)})_{n \geq 0}$.

For the CAM, the sequence of volume at birth $(V_b^{(n)})_{n \geq 0}$ follows $\forall\, n \geq 0$ the equation

$$V_b^{(n+1)} = \max\Big(\tilde{V}_i^{(n)} + \Delta_{id}^{(n)},\ 2^{-1}(V_b^{(n)} + \Delta_d^{(n)})\Big), \tag{12}$$

with $(\Delta_d^{(n)})_{n \geq 0}$ and $(\Delta_{id}^{(n)})_{n \geq 0}$ i.i.d. random variables sequences.

##### 5.1 Expressions of the conditional densities

In what follows we will denote by $F_X$ the cumulative distribution function of a random variable $X$.

**Proposition 5.**

*Let $p(\bullet|v_b, \tilde{v}_i, \lambda)$ be the conditional density of the division volume $2V_b^{(\bullet+1)}$ given that the triplet $(V_b^{(\bullet)}, \tilde{V}_i^{(\bullet)}, \Lambda^{(\bullet)})$ is equal to $(v_b, \tilde{v}_i, \lambda) \in \mathbb{R}_+^{*\,3}$. Let $f_R$, $f_{\Delta_d}$, $f_{\Delta_{id}}$ be the densities and $F_R$, $F_{\Delta_d}$, $F_{\Delta_{id}}$ the cumulative probability functions of the laws of $R$, $\Delta_d$, and $\Delta_i$ respectively. For the different models, the conditional distribution of the division volume satisfies for the different models the following expressions for any $v > 0$:*

- *The DIAM:*
$$p(v|v_b, \tilde{v}_i, \lambda) = f_{\Delta_d}(v - v_b), \tag{13a}$$
- *The RDAM:*
$$p(v|v_b, \tilde{v}_i, \lambda) = \frac{1}{2} f_{\Delta_{id}}\Big(\frac{v}{2} - \tilde{v}_i\Big), \tag{13b}$$
- *The CPM:*
$$p(v|v_b, \tilde{v}_i, \lambda) = f_{\Delta_d}(v - v_b) F_R\left(\frac{1}{\lambda}\log\left(\frac{v}{2\tilde{v}_i}\right)\right) + \frac{1}{\lambda v} f_R\left(\frac{1}{\lambda}\log\left(\frac{v}{2\tilde{v}_i}\right)\right) F_{\Delta_d}(v - v_b), \tag{13c}$$
- *The CAM:*
$$p(v|v_b, \tilde{v}_i, \lambda) = f_{\Delta_d}(v - v_b) F_{\Delta_{id}}\Big(\frac{v}{2} - \tilde{v}_i\Big) + \frac{1}{2} f_{\Delta_{id}}\Big(\frac{v}{2} - \tilde{v}_i\Big) F_{\Delta_d}(v - v_b). \tag{13d}$$

*Proof.* First, one may start from the definition of $p(\bullet|v_b, \tilde{v}, \lambda)$ for all $(v_b, \tilde{v}_i, \lambda) \in \mathbb{R}_+^{*\,3}$:

$$\forall v > 0, \quad \int_0^v p(x|v_b, \tilde{v}_i, \lambda)dx = \mathbb{P}[2V_b^{(\bullet+1)} \leq v | V_b^{(\bullet)} = v_b, \tilde{V}_i^{(\bullet)} = \tilde{v}_i, \Lambda^{(\bullet)} = \lambda].$$

One can express this quantity according to the different studied models.
For the DIAM by (7):

$$\begin{aligned}\mathbb{P}[2V_b^{(\bullet+1)} \leq v | V_b^{(\bullet)} = v_b, \tilde{V}_i^{(\bullet)} = \tilde{v}_i, \Lambda^{(\bullet)} = \lambda] &= \mathbb{P}[\Delta_d + v_b \leq v] \\ &= F_{\Delta_d}(v - v_b).\end{aligned}$$

For the RDAM by (10):

$$\begin{aligned}p[2V_b^{(\bullet+1)} \leq v | V_b^{(\bullet)} = v_b, \tilde{V}_i^{(\bullet)} = \tilde{v}_i, \lambda^{(\bullet)} = \lambda] &= \mathbb{P}[2(\Delta_{id} + \tilde{v}_i) \leq v] \\ &= F_{\Delta_{id}}\Big(\frac{v}{2} - \tilde{v}_i\Big).\end{aligned}$$

For the CPM by (11):

$$\begin{aligned}\mathbb{P}[2V_b^{(\bullet+1)} \leq v | V_b^{(\bullet)} = v_b, \tilde{V}_i^{(\bullet)} = \tilde{v}_i, \lambda^{(\bullet)} = \lambda] &= \mathbb{P}[2\tilde{v}_i e^{\lambda R} \leq v, \Delta_d + v_b \leq v] \\ &= \mathbb{P}[R \leq \frac{1}{\lambda}\log(\frac{v}{2\tilde{v}_i}), \Delta_d \leq v - v_b] \\ &= F_R(\frac{1}{\lambda}\log(\frac{v}{2\tilde{v}_i})) F_{\Delta_d}(v - v_b). \quad \text{(by independence)}\end{aligned}$$

For the CAM by (12):

$$\begin{aligned}\mathbb{P}[2V_b^{(\bullet+1)} \leq v | V_b^{(\bullet)} = v_b, \tilde{V}_i^{(\bullet)} = \tilde{v}_i, \lambda^{(\bullet)} = \lambda] &= \mathbb{P}[2(\tilde{v}_i + \Delta_{id}) \leq v, \Delta_d + v_b \leq v] \\ &= \mathbb{P}[\Delta_{id} \leq \frac{v}{2} - \tilde{v}_i, \Delta_d \leq v - v_b] \\ &= F_{\Delta_{id}}(\frac{v}{2} - \tilde{v}_i) F_{\Delta_d}(v - v_b) \quad \text{(by independence)}\end{aligned}$$

By taking the derivative of the obtained expressions according to $v$, one ends up with the expected results.

□

**Remark:** From these expressions of the conditional densities of the division size, one can obtain the division size distribution by integrating these expressions against the joint distribution of $(V_b^{(\bullet)}, \tilde{V}_i^{(\bullet)}, \Lambda^{(\bullet)})$. As an example, in Figure S3, we integrate the estimated conditional densities against the empirical distribution from the experimental data of birth size, initiation size and elongation rate $(v_b^k, \tilde{v}_i^k, \lambda^k)_{1 \leq k \leq N}$, providing the following division size distribution:

$$f_{V_d}(v) = \frac{1}{N} \sum_{k=1}^{N} p(v | v_b^k, \tilde{v}_i^k, \lambda^k).$$

#### 5.2 Identifiability of the conditional densities of the CPM and CAM

While it is immediate that the distribution of $\Delta_d$ in the DIAM, and the distribution of $\Delta_{id}$ in the RDAM are identifiable as they are observable, it is less clear that the distributions of $\Delta_d$, $R$ and $\Delta_{id}$ are identifiable in the CPM and the CAM, as these are latent variables. In the following, we investigate this question and show that under specific conditions over the observed covariates in the datasets, the latent variables in the CAM and CPM are identifiable.

Let us consider the general case where we have a collection of $n$ realizations of a random variable $Y$ with values in $\mathcal{Y}$ that is the maximum between two random variables such that

$$Y := \max\Big(h_X(Z_1),\ g_X(Z_2)\Big), \tag{14}$$

where $X$ is a collection of covariates that takes values in $\mathcal{X} \subseteq \mathbb{R}^d$, $Z_1$ and $Z_2$ are independent random variables distributed according to parametric cumulative distribution functions $F^1_{\theta_1}$ and $F^2_{\theta_2}$ with parameters $(\theta_1, \theta_2) \in \Theta_1 \times \Theta_2$. We assume that $F^1_{\theta_1}$ and $F^2_{\theta_2}$ are positive on $(0, +\infty)$ and equal to zero on $(-\infty, 0]$ for any $(\theta_1, \theta_2) \in \Theta_1 \times \Theta_2$, and that they belong to a class of distributions that are right-tail identifiable. We call $F_\theta$ a right-tail identifiable distribution if

$$\forall \theta, \alpha \in \Theta,\ \forall y \in \mathbb{R}, \quad \{\forall x > y,\ F_\theta(x) = F_\alpha(x) \implies \theta = \alpha\}.$$

Then, we assume that $\forall x \in \mathcal{X},\ h_x : \mathbb{R} \to \mathcal{H} \supseteq \mathcal{Y}$ and $g_x : \mathbb{R} \to \mathcal{G} \supseteq \mathcal{Y}$ are bijective, increasing, and satisfy $h_x(\mathbb{R}_+) \subseteq \mathcal{Y}$ and $g_x(\mathbb{R}_+) \subseteq \mathcal{Y}$.

For all $\theta \in \Theta_1 \times \Theta_2$, we denote by $p^\theta_{Y|X}(\bullet | x)$ the conditional density of $Y|X = x$ which allows us to define the conditional likelihood over a sample of $n$ independent realizations of $Y$ conditionally to $n$ realizations of $X$:

$$\begin{aligned} L^n : \mathcal{Y}^n \times \mathcal{X}^n \times \Theta_1 \times \Theta_2 &\longrightarrow \mathbb{R}_+ \\ (y_1, ..., y_n, x_1, ..., x_n, \theta) &\longmapsto \prod_{k=1}^{n} p^\theta_{Y|X}(y_k | x_k). \end{aligned}$$

We consider that the distributions of $Z_1$ and $Z_2$ are identifiable for a set of covariates defined on a space $E \subseteq \mathcal{X}^n$ if

$$\forall\, \mathbf{x} \in E,\ \forall\, \theta, \alpha \in \Theta_1 \times \Theta_2, \quad L^n(\bullet, \mathbf{x}, \theta) = L^n(\bullet, \mathbf{x}, \alpha) \implies \theta = \alpha.$$

In the following, we study a sufficient condition on $E$ in order to have $Z_1$ and $Z_2$ identifiable.

**Proposition 6.** *Let $E := \{(x_1, ..., x_n) \in \mathcal{X}^n :\ \exists\, i, j,\ \forall z > 0,\ g_{x_i}^{-1} \circ h_{x_i}(z) < g_{x_j}^{-1} \circ h_{x_j}(z)\}$ be the observed covariates space, with $n \geq 2$, $\forall\, \boldsymbol{x} \in E,\ \forall\, \theta, \alpha \in \Theta_1 \times \Theta_2$,*

$$L^n(\bullet, \boldsymbol{x}, \theta) = L^n(\bullet, \boldsymbol{x}, \alpha) \implies \theta = \alpha.$$

*Proof.* If $\forall\, \mathbf{x} := (x_1, ..., x_n) \in E,\ \forall\, \theta, \alpha \in \Theta_1 \times \Theta_2, \quad L^n(\bullet, \mathbf{x}, \theta) = L^n(\bullet, \mathbf{x}, \alpha)$ then $\forall\, (y_1, ..., y_n) \in \mathcal{Y}^n$,

$$\begin{aligned}
& \int_{-\infty}^{y_1} ... \int_{-\infty}^{y_n} L^n(\mathbf{x}, \mathbf{y}, \theta) d\mathbf{y} = \int_{-\infty}^{y_1} ... \int_{-\infty}^{y_n} L^n(\mathbf{x}, \mathbf{y}, \alpha) d\mathbf{y} \\
\iff\ & \prod_{k=1}^{n} \mathbb{P}_\theta[Y \leq y_k | X = x_k] = \prod_{k=1}^{n} \mathbb{P}_\alpha[Y \leq y_k | X = x_k] \\
\iff\ & \prod_{k=1}^{n} \mathbb{P}_\theta[Z_1 \leq h_{x_k}^{-1}(y_k)] \mathbb{P}_\theta[Z_2 \leq g_{x_k}^{-1}(y_k)] = \prod_{k=1}^{n} \mathbb{P}_\alpha[Z_1 \leq h_{x_k}^{-1}(y_k)] \mathbb{P}_\alpha[Z_2 \leq g_{x_k}^{-1}(y_k)],
\end{aligned}$$

by independence of $Z_1$ and $Z_2$.

We then denote by $F^1_{\theta_1}$, $F^1_{\alpha_1}$, $F^2_{\theta_2}$ and $F^2_{\alpha_2}$ the cumulative distribution functions of $Z_1$ and $Z_2$, with $(\theta_1, \theta_2) := \theta$ and $(\alpha_1, \alpha_2) := \alpha$, and then consider the equation when it is positive, namely $\forall\, (y_1, ..., y_n) \in \mathcal{Y}^n_+ := \{(y_1, ..., y_n) \in \mathcal{Y}^n : \forall k,\ y_k > h_{x_k}(0) \vee g_{x_k}(0)\}$, in which case we have,

$$\prod_{k=1}^{n} \frac{F^1_{\theta_1}(h_{x_k}^{-1}(y_k))}{F^1_{\alpha_1}(h_{x_k}^{-1}(y_k))} = \prod_{k=1}^{n} \frac{F^2_{\alpha_2}(g_{x_k}^{-1}(y_k))}{F^2_{\theta_2}(g_{x_k}^{-1}(y_k))}.$$

To simplify, we denote $H^1 := F^1_{\theta_1}/F^1_{\alpha_1}$ and $H^2 := F^2_{\alpha_2}/F^2_{\theta_2}$. The objective in the following is to show that $\exists\, y,\ \forall\, x > y,\ H^1(x) = H^2(x) = 1$ which will conclude that $\theta = \alpha$ by the right-tail identifiability property of $F^1$ and $F^2$. One can rewrite the previous equation as following:

$$\prod_{k=1}^{n} H^1 \circ h_{x_k}^{-1}(y_k) = \prod_{k=1}^{n} H^2 \circ g_{x_k}^{-1}(y_k).$$

This equality is equivalent for any $1 \leq k \leq n$ to

$$H^1 \circ h_{x_k}^{-1}(y_k) = H^2 \circ g_{x_k}^{-1}(y_k) \underbrace{\frac{\prod_{j \neq k} H^2 \circ g_{x_j}^{-1}(y_j)}{\prod_{j \neq k} H^1 \circ h_{x_j}^{-1}(y_j)}}_{=:\ C_k(y_1, ..., y_{k-1}, y_{k+1}, ..., y_n)}.$$

For each $k$, one can fix $y_1, ..., y_{k-1}, y_{k+1}, ... y_n$ and notice that $\forall y > h_{x_k}(0) \vee g_{x_k}(0)$,

$$H^1 \circ h_{x_k}^{-1}(y) = H^2 \circ g_{x_k}^{-1}(y) C_k(y_1, ..., y_{k-1}, y_{k+1}, ..., y_n),$$

and thus conclude that, $\forall\, k,\ \exists\, C_k > 0$, such that

$$\forall\, y > h_{x_k}(0) \vee g_{x_k}(0), \quad H^1 \circ h_{x_k}^{-1}(y) = C_k\ H^2 \circ g_{x_k}^{-1}(y).$$

As both $H^1$ and $H^2$ converge to one at $+\infty$, one can deduce that $\forall k$, $C_k = 1$, and thus

$$\forall\ k, \quad \forall\ y > h_{x_k}(0) \vee g_{x_k}(0), \quad H^1 \circ h_{x_k}^{-1}(y) = H^2 \circ g_{x_k}^{-1}(y). \qquad (*)$$

From $(*)$ one can deduce the following assertion:

$$\forall\ k, l, \quad \forall\ z > 0 \vee h_{x_k}^{-1} \circ g_{x_k}(0) \vee h_{x_l}^{-1} \circ g_{x_l}(0), \quad H^1(z) = H^2 \circ g_{x_k}^{-1} \circ h_{x_k}(z) = H^2 \circ g_{x_l}^{-1} \circ h_{x_l}(z).$$

Since $\forall\ (x_1, ..., x_n) \in E,\ \exists\ i, j,\ \forall z > 0,\ g_{x_i}^{-1} \circ h_{x_i}(z) < g_{x_j}^{-1} \circ h_{x_j}(z)$, it follows that for these particular $i, j$,

$$\forall\ z > 0 \vee g_{x_i}^{-1} \circ h_{x_i}(0), \qquad H^2(z) = H^2 \circ \varphi(z), \quad \text{with} \quad \varphi(z) := g_{x_j}^{-1} \circ h_{x_j} \circ h_{x_i}^{-1} \circ g_{x_i}(z) > z.$$

On one hand this provides us that $\forall\, n,\ H^2 = H^2 \circ \varphi^{\circ n}$ (with $\varphi^{\circ n} = \varphi \circ ... \circ \varphi$ $n$ times) and on another hand that $\lim_{n \to \infty} \varphi^{\circ n}(z) = +\infty$. As $H^2$ converges to one at $+\infty$, this concludes that $\forall z > 0 \vee g_{x_i}^{-1} \circ h_{x_i}(0),\ H^2(z) = 1$, and equivalently $\forall z > 0 \vee h_{x_i}^{-1} \circ g_{x_i}(0),\ H^2 \circ g_{x_j}^{-1} \circ h_{x_j}(z) = H^1(z) = 1$.

□

One can derive the corollaries of this result by defining the sufficient conditions to verify in the data in order to have the distributions of the latent variables in the CAM and the CPM identifiable.

In both models, the random variable $Y$ refers to $V_b^{(\bullet+1)}$ supported on $\mathbb{R}_+^*$, and the covariate $X$ refers to $(V_b^{(\bullet)}, \tilde{V}_i^{(\bullet)}, \lambda^{(\bullet)}) \in \mathbb{R}_+^{*\,3}$. For the CPM, $Z_1$ and $Z_2$ may refer to $\Delta_d$ and $R$ which thus identify $\forall (v', \tilde{v}', \lambda) \in \mathbb{R}_+^{*\,3}$,

$$h_{(v',\tilde{v}',\lambda)} : \mathbb{R} \to \mathbb{R}_+^* \qquad\qquad g_{(v',\tilde{v}',\lambda)} : \mathbb{R} \to \mathbb{R}$$
$$x \mapsto \tilde{v}' \exp(\lambda x) \qquad\qquad x \mapsto 2^{-1}(x + v').$$

Since $\Delta_d$ and $R$ are assumed to be distributed according to a Generalized Gamma distribution, they satisfy the conditions verified by $F_{\theta_1}^1$ and $F_{\theta_2}^2$. Thus, according to Proposition 6, the distributions of $\Delta_d$ and $R$ are identifiable if among the data there are $(v_1', \tilde{v}_1', \lambda_1)$, and $(v_2', \tilde{v}_2', \lambda_2)$ such that

$$\forall z > 0, \quad 2\tilde{v}_1' e^{\lambda_1 z} - v_1' < 2\tilde{v}_2' e^{\lambda_2 z} - v_2'.$$

For the CAM, $Z_1$ and $Z_2$ may refer to $\Delta_d$ and $\Delta_{id}$ which thus identify $\forall (v', \tilde{v}', \lambda) \in \mathbb{R}_+^{*\,3}$,

$$h_{(v',\tilde{v}',\lambda)} : \mathbb{R} \to \mathbb{R} \qquad\qquad g_{(v',\tilde{v}',\lambda)} : \mathbb{R} \to \mathbb{R}$$
$$x \mapsto x + \tilde{v}' \qquad\qquad x \mapsto 2^{-1}(x + v').$$

As for the CPM, since $\Delta_d$ and $\Delta_{id}$ are distributed according to a Generalized Gamma distribution one can apply Proposition 6, and state that $\Delta_d$ and $\Delta_{id}$ are identifiable if among the data there are $(v_1', \tilde{v}_1', \lambda_1)$, and $(v_2', \tilde{v}_2', \lambda_2)$ such that

$$2\tilde{v}_1' - v_1' < 2\tilde{v}_2' - v_2'.$$

In practice it is easy to verify in every dataset that there exist $(v_1', \tilde{v}_1', \lambda_1)$, and $(v_2', \tilde{v}_2', \lambda_2)$ such that $v_2' < v_1'$, $\tilde{v}_1' < \tilde{v}_2'$, and $\lambda_1 < \lambda_2$, which implies identifiability in both CPM and CAM.

##### 5.3 Expectation-Maximization (EM) algorithm for parameters estimation of the CPM and the CAM

In the previous section we provide sufficient conditions for the identifiability of the distributions of the latent variables $Z_1$ and $Z_2$ involved in Equation (14). Here we provide a method for estimating the parameters of the these distributions, involving an Expectation-Maximization (EM) algorithm. Adding to the hypotheses of Equation (14), we assume that $\forall x$ $g_x^{-1}$ and $h_x^{-1}$ are differentiable. First, one may notice that,

$$\begin{aligned}\mathbb{P}[Y \leq y, h_X(Z_1) \leq g_X(Z_2)|X = x] &= \mathbb{P}[h_x(Z_1) \leq g_x(Z_2) \leq y] \\ &= \int_0^y \mathbb{P}[h_x(Z_1) \leq s]\ dF^2_{\theta_2}(g_x^{-1}(s)) \\ &= \int_0^y \mathbb{P}[h_x(Z_1) \leq s]\ (g_x^{-1})'(s)\ f^2_{\theta_2}(g_x^{-1}(s))ds \\ &= \int_0^y F^1_{\theta_1}(h_x^{-1}(y))\ (g_x^{-1})'(s) f^2_{\theta_2}(g_x^{-1}(s))ds\end{aligned}$$

and similarly that,

$$\begin{aligned}\mathbb{P}[Y \leq y, g_X(Z_2) \leq h_X(Z_1)|X = x] &= \mathbb{P}[g_x(Z_2) \leq h_x(Z_1) \leq y] \\ &= \int_0^y \mathbb{P}[g_x(Z_2) \leq s]\ dF^1_{\theta_1}(h_x^{-1}(s)) \\ &= \int_0^y \mathbb{P}[g_x(Z_2) \leq s]\ (h_x^{-1})'(s)\ f^1_{\theta_1}(h_x^{-1}(s))ds \\ &= \int_0^y F^2_{\theta_2}(g_x^{-1}(s))\ (h_x^{-1})'(s)\ f^1_{\theta_1}(h_x^{-1}(s))ds.\end{aligned}$$

One can then introduce a random variable $W$ taking values in $\{1, 2\}$, such that $\{W = 1\} = \{h_X(Z_1) \geq g_X(Z_2)\}$ and $\{W = 2\} = \{h_X(Z_1) < g_X(Z_2)\}$. Let $p^\theta_{Y,W|X}$ be the joint distribution of $(Y, W)$ conditionally to $X$, with $\theta = (\theta_1, \theta_2)$. One can deduce from the previous equalities that it satisfies,

$$p^\theta_{Y,W|X}(y, w|x) = \begin{cases} (h_x^{-1})'(y)\ f^1_{\theta_1}(h_x^{-1}(y))\ F^2_{\theta_2}(g_x^{-1}(y)) & \text{if } w = 1 \\ (g_x^{-1})'(y)\ f^2_{\theta_2}(g_x^{-1}(y))\ F^1_{\theta_1}(h_x^{-1}(y)) & \text{if } w = 2. \end{cases}$$

Note that $p^\theta_{Y|X}(\bullet|x)$, the previously introduced conditional distribution of $Y$ given $X = x$, satisfies $p^\theta_{Y|X}(\bullet|x) = p^\theta_{Y,W|X}(\bullet, 1|x) + p^\theta_{Y,W|X}(\bullet, 2|x)$. By taking $L^N$ the likelihood previously defined, one can introduce the log-likelihood defined for $\mathbf{x} \in \mathcal{X}^n, \mathbf{y} \in \mathcal{Y}^n, \theta \in \Theta_1 \times \Theta_2$ by

$$\log L^N(\mathbf{y}, \mathbf{x}, \theta) = \sum_{k=1}^N \log p^\theta_{Y|X}(y_k|x_k).$$

Using Bayes formula, one can show [S16] for any $\theta' \in \Theta_1 \times \Theta_2$ that the log likelihood satisfies

$$\log L^N(\mathbf{y}, \mathbf{x}, \theta) = \underbrace{\sum_{k=1}^{N} \mathbb{E}_{\theta'} \left[ \log p^{\theta}_{Y,W|X}(Y, W|X) \middle| X = x_k, Y = y_k \right]}_{Q(\theta;\theta')}$$
$$- \underbrace{\sum_{k=1}^{N} \mathbb{E}_{\theta'} \left[ \log p^{\theta}_{W|X,Y}(W|X, Y) \middle| X = x_k, Y = y_k \right]}_{H(\theta;\theta')}$$
$$\geq Q(\theta; \theta') - H(\theta'; \theta') \quad \text{(by Gibb's inequality)},$$

with $p^{\theta}_{W|X,Y}(w|x, y) := \mathbb{P}_{\theta}[W = w|X = x, Y = y]$.

The fundamental principle of the EM algorithm is to iteratively maximize $\left(Q(\theta, \theta^{(h)})\right)_h$, such that $\theta^{(h+1)} = \arg\max_{\theta} Q(\theta; \theta^{(h)})$, which ensures the convergence of $(\theta^{(h)})_h$ to a local maximum of $\log L^N$ [S16]. In the following we detail $Q(\theta; \theta')$. To simplify, we denote for all $k$, $\pi^{\theta,k}_w := p^{\theta}_{W|X,Y}(w|x_k, y_k)$:

$$\begin{aligned}
Q(\theta; \theta') =& \sum_{k=1}^{N} \int \log p^{\theta}_{Y,W|X}(y_k, w|x_k) \; dp^{\theta'}_{W|X,Y}(w|x_k, y_k) \\
=& \sum_{k=1}^{N} \left( \log p^{\theta}_{Y,W|X}(y_k, 1|x_k) \; \pi^{\theta',k}_1 + \log p^{\theta}_{Y,W|X}(y_k, 2|x_k) \; \pi^{\theta',k}_2 \right) \\
=& \sum_{k=1}^{N} \left( \; \pi^{\theta',k}_1 \log \left( (h^{-1}_{x_k})'(y_k) \; f^1_{\theta_1}(h^{-1}_{x_k}(y_k)) \right) + \pi^{\theta',k}_2 \log F^1_{\theta_1}(h^{-1}_{x_k}(y_k)) \; \right) \\
&+ \sum_{k=1}^{N} \left( \; \pi^{\theta',k}_1 \log F^2_{\theta_2}(g^{-1}_{x_k}(y_k)) + \pi^{\theta',k}_2 \log \left( (g^{-1}_{x_k})'(y_k) \; f^2_{\theta_2}(g^{-1}_{x_k}(y_k)) \right) \; \right) \\
=& \underbrace{\sum_{k=1}^{N} \left( \; \pi^{\theta',k}_1 \log f^1_{\theta_1}(h^{-1}_{x_k}(y_k)) + \pi^{\theta',k}_2 \log F^1_{\theta_1}(h^{-1}_{x_k}(y_k)) \; \right)}_{Q^1(\theta_1;\theta')} + \sum_{k=1}^{N} \pi^{\theta',k}_1 \log(h^{-1}_{x_k})'(y_k) \\
&+ \underbrace{\sum_{k=1}^{N} \left( \; \pi^{\theta',k}_1 \log F^2_{\theta_2}(g^{-1}_{x_k}(y_k)) + \pi^{\theta',k}_2 \log f^2_{\theta_2}(g^{-1}_{x_k}(y_k)) \; \right)}_{Q^2(\theta_2;\theta')} + \sum_{k=1}^{N} \pi^{\theta',k}_2 \log(g^{-1}_{x_k})'(y_k).
\end{aligned}$$

One can observe that $Q(\theta; \theta')$ can be decomposed into the sum of three terms: a function of $\theta_1$ denoted by $Q^1(\theta_1; \theta')$; a function of $\theta_2$ denoted by $Q^2(\theta_2; \theta')$; and some quantities independent of $\theta$. This allows us to maximize $Q(\theta; \theta')$ by independently maximizing $Q^1(\theta_1; \theta')$ with respect to $\theta_1$ and $Q^2(\theta_2; \theta')$ with respect to $\theta_2$. This provides the following EM algorithm to apply for the problem of interest.

**Expectation step:** Computation for all $1 \leq k \leq N$ of

$$\pi_1^{\theta^{(h)},k} = \frac{p_{Y,W|X}^{\theta^{(h)}}(y_k, 1|x_k)}{p_{Y|X}^{\theta^{(h)}}(y_k|x_k)}, \qquad \pi_2^{\theta^{(h)},k} = 1 - \pi_1^{\theta^{(h)},k}.$$

**Maximization step:** Computation of $\theta^{(h+1)} := (\theta_1^{(h+1)}, \theta_2^{(h+1)})$ with

$$\theta_1^{(h+1)} = \arg\max_{\theta_1 \in \Theta_1} Q^1(\theta_1; \theta^{(h)}), \qquad \theta_2^{(h+1)} = \arg\max_{\theta_2 \in \Theta_2} Q^2(\theta_2; \theta^{(h)}).$$

In the context of the CPM and the CAM, the two latent variables $Z_1$ and $Z_2$ follow Generalized Gamma distributions. Considering this probability distribution in the expression of $Q^1$ and $Q^2$, prevents us from deriving an analytical solution for their maximization. However, these optimization problems remain much simpler than the original one and can be efficiently solved using numerical optimization algorithms such as Nelder-Mead (implemented in the package Scipy of python [S23]) and CMA-ES [S11] (implemented in the package cma in Python).

##### 5.4 The Generalized Gamma distribution

In this work, the independent and identically distributed sequences of random variables involved in the models have been assumed to be distributed according to a parameteric unimodal density called the Generalized Gamma distribution. This distribution well captures the empirical distribution of cell variables such as the inter-division added size, the C+D period and the added size per origin over the C+D period (see Figure S2). As described in [S6, S15], this density involves 3 parameters $\lambda \neq 0$, $\beta$ and $\sigma > 0$, satisfying the following equivalence:

$$A \sim \text{GenGamma}(\lambda, \beta, \sigma) \quad \iff \quad A \overset{D}{=} \left(\frac{W}{e^\beta}\right)^{\frac{\lambda}{\sigma}}, \text{ with } W \sim \text{Gamma}(1, \lambda^{-2}).$$

This provides the following expression for the density:

$$\forall x > 0, \quad f(x) = \frac{|\lambda|}{\sigma x \Gamma(\lambda^{-2})} \Big(\lambda^{-2} \Big(\frac{x}{e^\beta}\Big)^{\frac{\lambda}{\sigma}}\Big)^{\lambda^{-2}} \exp\Big( - \lambda^{-2} \big(\frac{x}{e^\beta}\big)^{\frac{\lambda}{\sigma}}\Big).$$

This definition encompasses multiple distributions namely the Gamma distribution when $\lambda = \sigma$, the Weibull distribution when $\lambda = 1$, as well as the inverse Weibull distribution when $\lambda = -1$. Additionnaly, one has that as $\lambda$ converges to 0, the density converges to the density of a Log-Normal distribution (specifically the density of a normally distributed random variable of mean $\beta$ and variance $\sigma^2$ after applying the exponential transformation).

#### 6 Discussion on Asymmetric Division

In the main article, we propose a framework in which cell division is assumed to be perfectly symmetric (the daughter cell size is equal to half of the mother's). This assumption has been shown to have strong consequences in the correlation analyses led in previous studies [S24, S13, S5], and to affect the cell size distribution [S12]. In this section, we extend the framework by relaxing the symmetric division assumption and discuss the results of this work under this relaxed

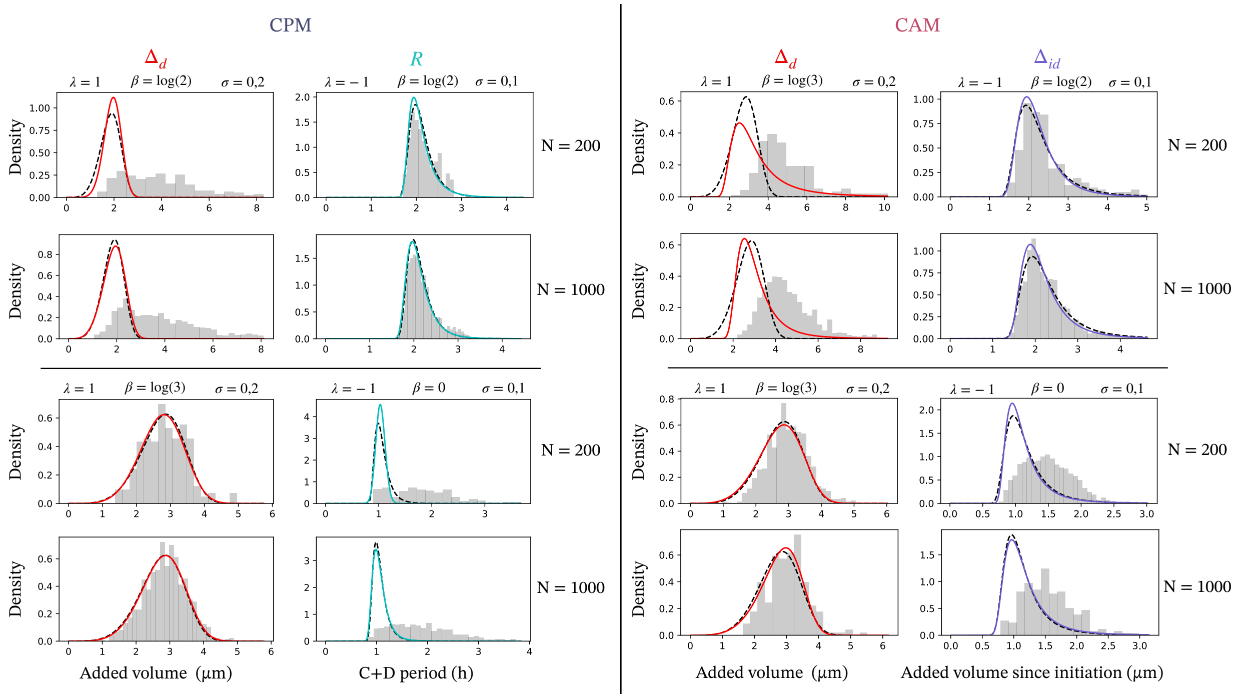


Figure 2: **Testing distribution estimation on synthetic data with known parameters** To evaluate the accuracy of our CPM and CAM distribution inference methods, we generated synthetic data using two sets of known parameters and two sample sizes. The latent random variables have been assumed to be distributed according to a Generalized Gamma distribution. The graphs display the estimated distribution (solid colored line), the true target distribution (dashed black line), and the synthetic data (gray histogram). The parameter sets were chosen to reflect scenarios where replication has minimal or strong influence on division control. Based on the experimental data sizes (ranging from 215 to 1806), we selected sample sizes $N = 200$ and $N = 1000$ for testing. The inferred distributions closely match the targets in these cases. However, the method may show severe limitations when replication rarely or exclusively limits division, and when the sample size is small.

assumption. Specially, we formalise the asymmetric division and show that the symmetric division assumption has a minimal impact on distribution inference of the control variables, resulting in no effect on the comparison method.

We introduce $(\alpha^{(n)})_n$ the sequence of random variables distributed on $(0,1)$, such that $\alpha^{(n)}$ is the ratio between the birth size of the daughter cell and the size of its mother at the division occurring at $T_b^{(n)}$. This provides a new expression for Equation (2),

$$\forall n \geq 0, \quad V_b^{(n+1)} = \alpha^{(n+1)} V_b^{(n)} \exp\left(\int_{T_b^{(n)}}^{T_b^{(n+1)}} \lambda(s) ds\right). \tag{15}$$

For Equation (4), the integration of the asymmetry in division is more subtle because the size per origin of replication is affected by the division events contrarily to the main framework of this work. If we consider $\tilde{V}_t$ the volume per origin of replication at time $t$ over the lineage, one has that for every division event $T_b^{(n)}$, the actual cell volume is multiplied by $\alpha^{(n)}$ and the number of origins of replication is divided by two, which provides that $\tilde{V}_{T_b^{(n)}} = 2\alpha^{(n)} \tilde{V}_{(T_b^{(n)})^-}$. One can thus derive the following equation linking initiation events,

$$\forall n \geq 0, \quad \tilde{V}_i^{(n+1)} = \frac{1}{2} \prod_{k \in \mathcal{D}_{ii}^{(n)}} (2\alpha^{(k)})\ \tilde{V}_i^{(n)} \exp\left(\int_{T_i^{(n)}}^{T_i^{(n+1)}} \lambda(s) ds\right), \tag{16}$$

with $\mathcal{D}_{ii}^{(n)} := \{k : T_i^{(n)} < T_b^{(k)} \leq T_i^{(n+1)}\}$ the set of indices of division dates which occur between the $n$-th and the $n+1$-th initiation of replication.

With the the same reasoning as in Lemma 1, assuming for the initial condition $V_b^{(1)} = \tilde{V}_i^{(0)} \exp\left(\int_{T_i^{(0)}}^{T_b^{(1)}} \lambda(t) dt\right)$, one can derive a new expression for Equation (6) coupling the replication and division cycles, that is,

$$\forall n \geq 0, \quad V_b^{(n+1)} = \prod_{k \in \mathcal{D}_{ib}^{(n)}} (2\alpha^{(k)})\ \tilde{V}_i^{(n)} \exp\left(\int_{T_i^{(n)}}^{T_b^{(n+1)}} \lambda(s) ds\right), \tag{17}$$

with $\mathcal{D}_{ib}^{(n)} := \{k : T_i^{(n)} < T_b^{(k)} \leq T_b^{(n+1)}\}$ being the set of index of division dates which occur over the C+D period $(T_i^{(n)}, T_b^{(n+1)}]$.

#### 6.1 Formulation of the models under asymmetric division

##### Division Adder

The division adder is straightforward under asymmetric division,

$$V_b^{(n+1)} = \alpha^{(n+1)} (V_b^{(n)} + \Delta_d^{(n)}). \tag{18}$$

##### Initiation-Division Timer

The initiation-division timer involved in the CPM is also straightforward thanks to the new Equation (17) that provides by simply substituting $T_b^{(n+1)}$ by $T_i^{(n)} + R^{(n)}$,

$$V_b^{(n+1)} = \prod_{k \in \mathcal{D}_{ib}^{(n)}} (2\alpha^{(k)}) \ \tilde{V}_i^{(n)} \exp\left( \int_{T_i^{(n)}}^{T_i^{(n)}+R^{(n)}} \lambda(s) ds \right). \tag{19}$$

**Initiation Adder**

The division of initiation adder is more subtle in the case of asymmetric division, as ambiguity may arise on what exactly the added size per origin of replication is, and how asymmetric division affects it. In the previous studies investigating the impact of asymmetric division on replication-division coordination [S24, S13, S5], simulations have relied on the following definition of $\Delta\tilde{V}_t^{(n)}$, the added size per origin of replication since $T_i^{(n)}$ over time $t$,

$$\begin{aligned} \Delta\tilde{V}_t^{(n)} &= A_{i,t}^{(n)} \int_{T_i^{(n)}}^{t} \lambda(s)\tilde{V}_s \ ds \\ &= \tilde{V}_t - A_{i,t}^{(n)}\tilde{V}_i^{(n)}, \end{aligned}$$

with $A_{i,t}^{(n)} := \prod_{\{k \ : \ T_i^{(n)} < T_b^{(k)} \leq t\}} (2\alpha^{(k)})$ and $\tilde{V}_t$ the volume per origin of replication at $t$ that satisfies for $t \in [T_i^{(n)}, T_i^{(n+1)})$, $\tilde{V}_t = A_{i,t}^{(n)}\tilde{V}_i^{(n)} \exp\left( \int_{T_i^{(n)}}^{t} \lambda(s) ds \right)$.

From this definition, one can define the next initiation event based on the initiation adder as follows, assuming that $(\Delta_i^{(n)})_n$ is an i.i.d. random variable sequence,

$$\forall n \geq 0, \qquad T_i^{(n+1)} = \inf\left\{ t > T_i^{(n)} \ : \ \Delta\tilde{V}_t^{(n)} \geq \Delta_i^{(n)} \right\}. \tag{20}$$

Interestingly, since $\Delta\tilde{V}_t^{(n)}$ is discontinuous at division events, one can imagine for a $k$ satisfying $T_b^{(k)} > T_i^{(n)}$, that $\Delta\tilde{V}_{(T_b^{(k)})^-}^{(n)} < \Delta_i^{(n)} < \Delta\tilde{V}_{T_b^{(k)}}^{(n)}$, in which case $T_i^{(n+1)} = T_b^{(k)}$, and therefore $\Delta\tilde{V}_{T_i^{(n+1)}}^{(n)} \neq \Delta_i^{(n)}$. This makes the initiation adder to be exclusively defined by the equation (20) which does not provide an explicit formulation for the sequence of initiation volumes as in Equation (8).

**Initiation-Division Adder**

The definition of the initiation-division adder in the RDAM under asymmetric division may also show ambiguity. We showed that the RDAM does not ensure that the sequence $(T_b^{(n)})_n$ is non-decreasing. Under asymmetric division, this asks in the case of a "daughter cell dividing before its mother" whether the asymmetric division of the daughter affects the size of the mother. In [S24, S13, S5] this ambiguity has been avoided, as the RDAM has been simulated under asymmetric division by constraining $(T_b^{(n)})_n$ to be non-decreasing, such that, assuming $(\Delta_{id}^{(n)})_n$ an i.i.d. sequence of random variables,

$$T_b^{(n+1)} = \inf\Big\{t \geq T_i^{(n)} \vee T_b^{(n)} \ : A_{id}^{(n)} \tilde{V}_i^{(n)} \Big( \exp\left( \int_{T_i^{(n)}}^t \lambda(s)ds \right) - 1 \Big) \geq \Delta_{id}^{(n)} \Big\}, \tag{21}$$

with $A_{id}^{(n)} := \prod_{\{k \ : \ T_i^{(n)} < T_b^{(k)} \leq T_b^{(n)}\}} (2\alpha^{(k)})$. From this definition, the event $T_b^{(n+1)} = T_b^{(n)}$ may arise with positive probability without assumptions on the distribution of $\alpha$ and $\Delta_{id}$, meaning that two successive division events can occur simultaneously. In such a case $V_b^{(n+1)} = \alpha^{(n+1)} V_b^{(n)}$ by Equation (15). Then, in the case where $T_b^{(n)} < T_b^{(n+1)}$, the volume at birth can be expressed as a function of $\Delta_{id}$: $V_b^{(n+1)} = 2\alpha^{(n+1)}(\Delta_{id}^{(n)} + A_{id}^{(n)} \tilde{V}_i^{(n)})$, by continuity and by Equation (17). Thus, the RDAM, as defined in Equation (21), satisfies

$$V_b^{(n+1)} = \max\Big(\alpha^{(n+1)} V_b^{(n)}, \quad 2\alpha^{(n+1)}(\Delta_{id}^{(n)} + A_{id}^{(n)} \tilde{V}_i^{(n)})\Big). \tag{22}$$

Studying the limitations of the RDAM according to this definition as we did in this work for the symmetric case, would be equivalent to show the necessary and sufficient conditions of having $T_b^{(n)} < T_b^{(n+1)}$ almost surely, which is equivalent to show the necessary and sufficient conditions of having

$$V_b^{(n)} < 2(\Delta_{id}^{(n)} + A_{id}^{(n)} \tilde{V}_i^{(n)}) \qquad a.s.$$

Finally, for the CAM, as the sequence of $(T_b^{(n)})_n$ is almost surely increasing thanks to the division adder, the sequence of birth volumes has the following explicit formulation,

$$V_b^{(n+1)} = \max\Big(\alpha^{(n+1)}\big(V_b^{(n)} + \Delta_d^{(n)}\big), \quad 2\alpha^{(n+1)}\big(\Delta_{id}^{(n)} + A_{id}^{(n)} \tilde{V}_i^{(n)}\big)\Big). \tag{23}$$

In conclusion, our framework can account for asymmetric divisions to express the sequences of birth volumes explicitly according to the models studied. However, the initiation adder relies on an implicit definition expressed in Equation (20), which does not provide an explicit formulation for the sequences of initiation volumes. Therefore, the mathematical analysis made in this work for the single-process models requires additional work to be adapted to the asymmetric case. Future investigations might be expected to clarify the definition of added size per origin under asymmetric division, in order to avoid simultaneous events of replication and division events. In the following section, we address the extent to which the symmetric division assumption we make in this work affects the distribution inference and model comparison method.

#### 6.2 Effect of relaxing symmetric division assumption on distributions inference and model comparison

In the statistical analysis, we only considered the conditional distribution of the division size, which relies on the explicit formulations defined in the previous section. Since we account for asymmetric division, the division size cannot be considered as $2V_b^{(n+1)}$ as in the main work. Therefore we introduce the sequence of division volumes $(V_d^{(n)})_n$, defined as $V_d^{(n)} := V_b^{(n+1)}/\alpha^{(n+1)}$.

When accounting for the division adder, under asymmetric division, the volume at division is expressed just like in the symmetric case:

$$V_d^{(n)} = V_b^{(n)} + \Delta_d^{(n)}.$$

Therefore, the conditional distribution of division size according to the division adder is unaffected by asymmetric division.

However, according to the initiation-division timer involved in the CPM, the volume at division is expressed as follows,

$$V_d^{(n)} = 2A_{id}^{(n)} \tilde{V}_i^{(n)} e^{\Lambda^{(n)} R^{(n)}},$$

with $\Lambda^{(n)} := \int_{T_i^{(n)}}^{T_i^{(n)}+R^{(n)}} \lambda(s)ds/R^{(n)}$, assumed to be independent of $R^{(n)}$.

Similarly, for the initiation-division adder - assuming that the division we observe always satisfies $T_b^{(n)} < T_b^{(n+1)}$ - we obtain the following expression of the division volume,

$$V_d^{(n)} = 2(\Delta_{id}^{(n)} + A_{id}^{(n)} \tilde{V}_i^{(n)}).$$

Using the previous expressions, one can define the distributions of the division volume $V_d^{(n)}$for the DIAM, RDAM, CPM and CAM conditionally to $(V_b^{(n)}, \tilde{V}_i^{(n)}, \Lambda^{(n)}, A_{id}^{(n)}) = (v_b, \tilde{v}_i, \lambda, a)$, $\forall v > 0$,

- The DIAM:
$$p(v|v_b, \tilde{v}_i, \lambda, a) = f_{\Delta_d}(v - v_b), \tag{24a}$$

- The RDAM:
$$p(v|v_b, \tilde{v}_i, \lambda, a) = \frac{1}{2} f_{\Delta_{id}}\Big(\frac{v}{2} - a\tilde{v}_i\Big), \tag{24b}$$

- The CPM:
$$p(v|v_b, \tilde{v}_i, \lambda, a) = f_{\Delta_d}(v - v_b) F_R\left(\frac{1}{\lambda}\log\left(\frac{v}{2a\tilde{v}_i}\right)\right) + \frac{1}{\lambda v} f_R\left(\frac{1}{\lambda}\log\left(\frac{v}{2a\tilde{v}_i}\right)\right) F_{\Delta_d}(v - v_b), \tag{24c}$$

- The CAM:
$$p(v|v_b, \tilde{v}_i, \lambda, a) = f_{\Delta_d}(v - v_b) F_{\Delta_{id}}\Big(\frac{v}{2} - a\tilde{v}_i\Big) + \frac{1}{2} f_{\Delta_{id}}\Big(\frac{v}{2} - a\tilde{v}_i\Big) F_{\Delta_d}(v - v_b). \tag{24d}$$

One can define a new likelihood based on these new expressions of the conditional distribution of division size,

$$L(\theta) = \prod_{k=1}^{N} p_\theta(v_d^k | v_b^k, \tilde{v}_i^k, \lambda^k, a_{id}^k), \tag{25}$$

with $(v_d^k, v_b^k, \tilde{v}_i^k, \lambda^k, a_{id}^k)_{1\le k\le N}$ the observations of $(V_b^{(n)}, \tilde{V}_i^{(n)}, \Lambda^{(n)}, A_{id}^{(n)})$ for $N$ cell cycles.

From this new likelihood, it is possible to compare models without using the symmetric division assumption, by using datasets containing cell cycles for which $A_{id}$ has been observed. The datasets from Si et al. (2019) allow to extract these observations. However, since $A_{id}$ has not been observed for every cell cycle, the comparison is restricted to smaller sample sizes than in the main work ($N$ = 1386, 1171, 1634 for MG1655 Acetate, Gly11aa, Glucose respectively, and $N$ = 1699, 1107, 758 for NCM3722 Arginine, Glucose, Glu12aa respectively). After computing the maximum of likelihood on these restricted datasets with and without the symmetric division assumption, it is possible to compare the BIC and AIC scores to evaluate the effect of this assumption. As shown in Table 1, the assumption of symmetric division does not affect the ranking of the models based on BIC and AIC scores.

Then, by using the estimated parameters from these maximum of likelihoods, we evaluate the effect of relaxing the symmetric division assumption on the estimation of the distribution of the control variables. As shown in Figure 3, the distributions of the control variables estimated with and without the symmetric division assumption closely match, suggesting a minimal impact of this assumption on the estimation of the distribution of the control variables.

Table 1: **BIC and AIC scores with and without symmetric division assumption** Using datasets from Si et al. (2019) restricted to cell cycles for which $A_{id}$ has been observed, we computed the BIC and AIC scores of the models with and without symmetric division assumption. Models minimizing BIC and AIC scores are highlighted in bold.

**BIC**

| | With symmetric division assumption | | | | Without symmetric division assumption | | | |
|---|---|---|---|---|---|---|---|---|
| Datasets | DIAM | RDAM | CPM | CAM | DIAM | RDAM | CPM | CAM |
| MG1655 acet | 501.16 | 322.90 | 381.30 | **133.12** | 501.16 | 312.18 | 380.82 | **135.75** |
| MG1655 gly11aa | 933.29 | 1135.70 | 922.41 | **914.65** | 933.29 | 1096.21 | 917.38 | **905.78** |
| MG1655 glu | 1054.11 | 1253.26 | 1037.13 | **981.16** | 1054.11 | 1194.26 | 1025.51 | **968.93** |
| NCM3722 arg | -260.43 | -358.83 | **-663.03** | -456.31 | -260.43 | -362.60 | **-663.17** | -457.19 |
| NCM3722 glu | 201.07 | 298.29 | **152.69** | 177.82 | 201.07 | 255.16 | **153.26** | 178.42 |
| NCM3722 glu12aa | 516.65 | 635.40 | **512.65** | 515.06 | 516.65 | 619.81 | **510.89** | 514.91 |

**AIC**

| | With symmetric division assumption | | | | Without symmetric division assumption | | | |
|---|---|---|---|---|---|---|---|---|
| Datasets | DIAM | RDAM | CPM | CAM | DIAM | RDAM | CPM | CAM |
| MG1655 acet | 493.31 | 315.04 | 365.60 | **117.42** | 493.31 | 304.33 | 365.12 | **120.04** |
| MG1655 gly11aa | 925.69 | 1128.10 | 907.21 | **899.89** | 925.69 | 1088.61 | 902.19 | **890.58** |
| MG1655 glu | 1046.01 | 1245.16 | 1020.93 | **964.96** | 1046.01 | 1186.16 | 1009.31 | **952.73** |
| NCM3722 arg | -268.59 | -366.99 | **-679.35** | -472.63 | -268.59 | -370.75 | **-679.48** | -473.50 |
| NCM3722 glu | 193.56 | 290.77 | **137.66** | 162.79 | 193.56 | 247.64 | **138.23** | 163.40 |
| NCM3722 glu12aa | 509.71 | 628.46 | **498.76** | 501.17 | 509.71 | 612.87 | **497.00** | 501.02 |

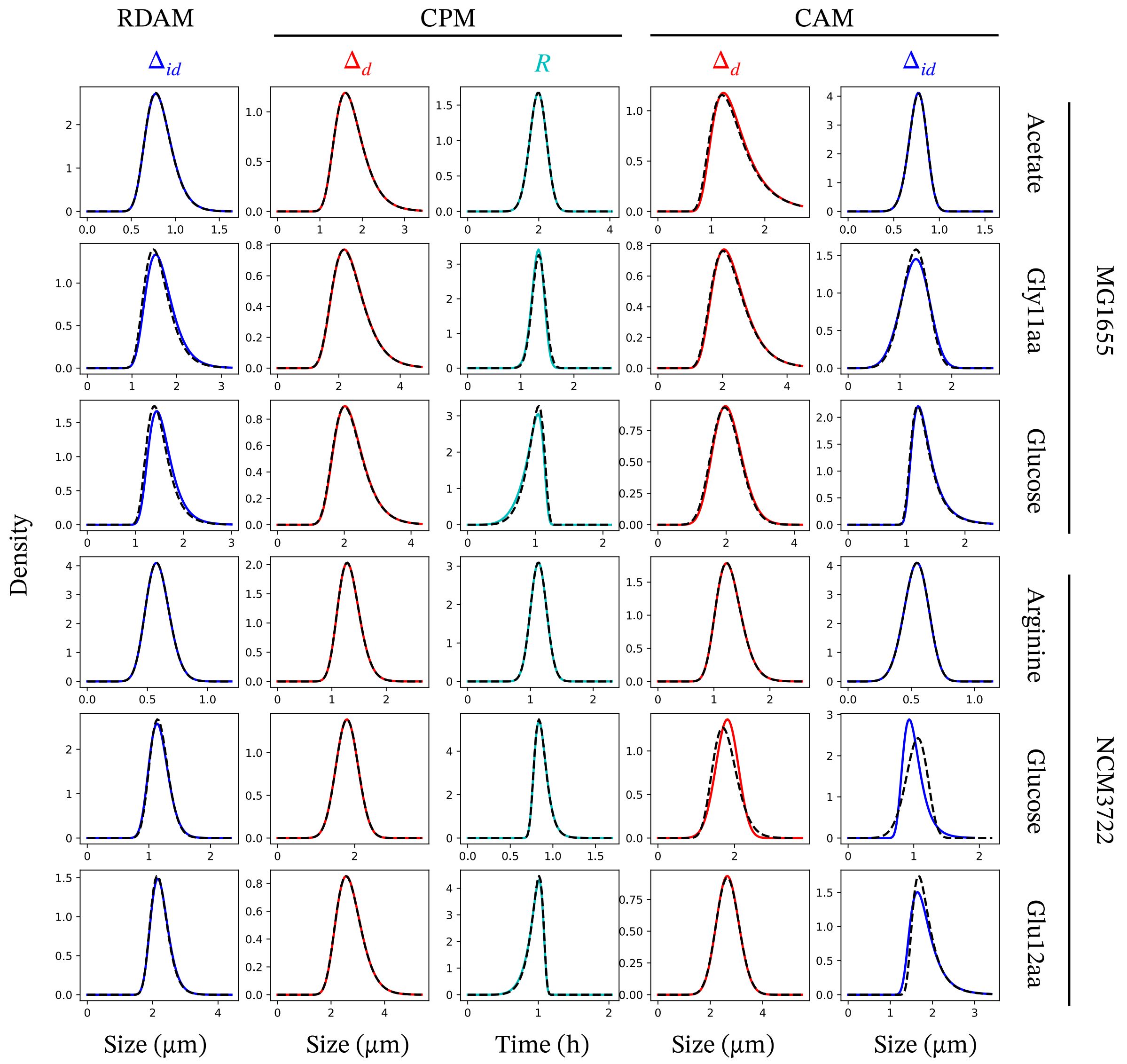


Figure 3: **Effect of the symmetric division assumption on distribution inference** Based on the likelihood maximization performed on the restricted datasets from Si et al. (2019), we have plotted the estimated distributions of the control variables with and without the symmetric division assumption. The distributions of $\Delta_d$ inferred in the DIAM are not shown, as they are exactly the same regardless of this assumption. The distributions of the control variables inferred without the symmetric division assumption are shown in black dashed lines. The distributions of the control variables inferred with the symmetric division assumption are shown in solid coloured lines.
