## Supplementary Figures for "Deciphering the Replication-Division Coordination in *E. coli* : A Unified Mathematical Framework for Systematic Model Comparison"

Table S1: **Estimation of the probability of $T_b^{(n+1)} < T_b^{(n)}$ in the RDAM.** From the datasets published in Si et al. (2019) and Witz et al. (2019), we have estimated the probability that a daughter cell divides before its mother, using the observations of $\Delta_i$ and $\Delta_{id}$ (Material and Methods)

| **Datasets** | **Probability of $T_b^{(n+1)} < T_b^{(n)}$** |
|---|---|
| MG1655 (Si et al.) Acetate | 1.23e-4 |
| MG1655 (Si et al.) Glycerol11aa | 5.78e-4 |
| MG1655 (Si et al.) Glucose | 6.49e-4 |
| NCM3722 (Si et al.) Arginine | 3.08e-7 |
| NCM3722 (Si et al.) Glucose | 2.64e-8 |
| NCM3722 (Si et al.) Glucose12aa | 7.88e-5 |
| BW27378 (Witz et al.) Glycerol | 1.05e-4 |
| BW27378 (Witz et al.) Glucose | 6.52e-4 |
| BW27378 (Witz et al.) Glucose8a | 3.97e-4 |

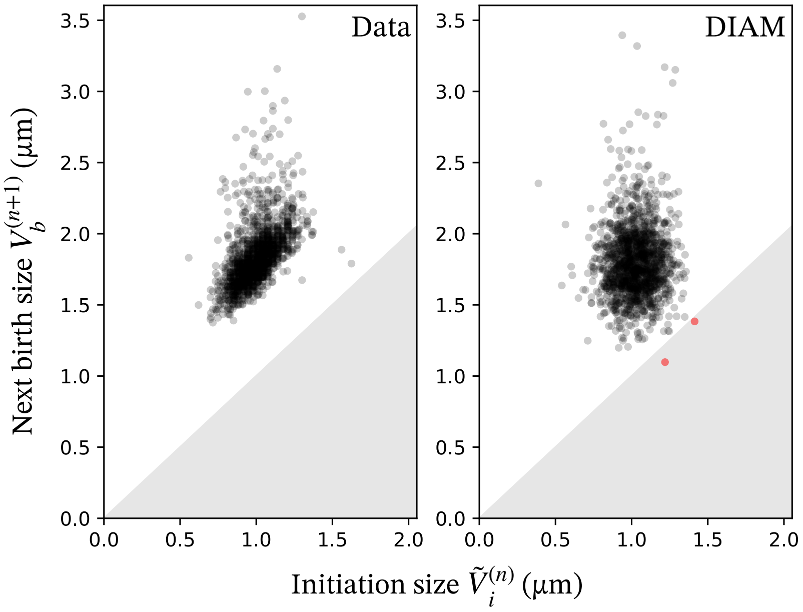


Figure S1: **Simulated cells with less than one origin of replication according to the DIAM.** Using the same simulations as the ones used to compute the probability of having a one-origin-cell to divide (see Material and methods) for the dataset MG1655 acetate, we display the simulated distribution of initiation size and birth size for N = 1554 cell cycles (right) in comparison with the experimental joint distribution of $(V_b^{(n+1)}, \tilde{V}_i^{(n)})$ from the dataset MG1655 acetate (left). The grey area $\{(x, y) : x > y\}$ is the area in which cells have less than one origin of replication at birth. As shown in red, in this simulation, the DIAM produces two cells with less than one origin of replication.

Table S2: **P-values of Shapiro–Wilk normality tests of cell quantities.** We performed normality tests on division, birth, and initiation sizes using the Shapiro–Wilk test. P-values greater than 0.01, indicating no significant deviation from a normal distribution, are shown in bold.

| **Datasets** | **Division size** | **Birth size** | **Initiation size** |
|---|---|---|---|
| MG1655 acet | 1.71e-30 | 4.17e-23 | 8.39e-08 |
| MG1655 gly11aa | 1.17e-25 | 1.64e-23 | 2.51e-11 |
| MG1655 glu | 3.60e-30 | 2.97e-29 | 3.08e-10 |
| NCM3722 arg | 9.37e-09 | 1.14e-12 | 9.82e-08 |
| NCM3722 glu | 1.18e-11 | 8.95e-12 | 3.21e-06 |
| NCM3722 glu12aa | 1.94e-23 | 2.32e-21 | 5.75e-20 |
| BW27378 gly | 1.03e-21 | 3.49e-27 | 1.30e-22 |
| BW27378 glu | 1.00e-25 | 5.80e-27 | 2.21e-19 |
| BW27378 glu8a | 8.81e-22 | 6.47e-20 | 1.23e-18 |
| BW27783 acet | 1.59e-07 | 3.09e-07 | 5.61e-07 |
| BW27783 alaTrE | 1.68e-06 | 2.06e-12 | 3.22e-12 |
| BW27783 man | **8.85e-01** | 1.28e-08 | 3.59e-09 |
| BW27783 gly | 1.45e-04 | 1.45e-12 | 1.76e-08 |
| BW27783 glyTre | 4.02e-05 | 8.14e-08 | **6.93e-01** |
| BW27783 glu | **1.82e-01** | **4.23e-01** | 2.95e-03 |
| BW27783 glyCas | 4.49e-05 | 8.21e-03 | 3.18e-04 |
| BW27783 gluCas | 1.32e-11 | 4.35e-06 | 6.58e-19 |

Table S3: **BIC and AIC scores.**

| | BIC scores | | | | AIC scores | | | |
|---|---|---|---|---|---|---|---|---|
| Datasets | DIAM | RDAM | CPM | CAM | DIAM | RDAM | CPM | CAM |
| MG1655 acet | 677.42 | 445.54 | 497.82 | 195.65 | 669.39 | 437.52 | 481.77 | 179.60 |
| MG1655 gly11aa | 1233.28 | 1494.29 | 1219.77 | 1198.14 | 1225.32 | 1486.33 | 1203.85 | 1182.22 |
| MG1655 glu | 1169.80 | 1364.29 | 1147.48 | 1076.43 | 1161.55 | 1356.04 | 1130.98 | 1059.93 |
| NCM3722 arg | -253.43 | -358.51 | -659.33 | -454.58 | -261.59 | -366.66 | -675.64 | -470.90 |
| NCM3722 glu | 269.03 | 378.46 | 188.57 | 221.41 | 261.13 | 370.56 | 172.77 | 205.61 |
| NCM3722 glu12aa | 989.38 | 1241.54 | 978.02 | 976.80 | 981.45 | 1233.61 | 962.15 | 960.94 |
| BW27378 gly | 140.21 | 101.21 | -24.41 | 18.88 | 133.18 | 94.17 | -38.47 | 4.81 |
| BW27378 glu | 412.90 | 630.59 | 343.75 | 409.96 | 405.58 | 623.26 | 329.10 | 395.31 |
| BW27378 glu8a | 766.69 | 886.78 | 701.16 | 741.83 | 759.20 | 879.29 | 686.19 | 726.85 |
| BW27783 acet | 54.27 | -37.54 | -10.12 | -42.00 | 48.28 | -43.53 | -22.10 | -53.99 |
| BW27783 alaTrE | 68.10 | 14.18 | 30.77 | 17.57 | 63.04 | 9.12 | 20.66 | 7.46 |
| BW27783 man | 114.00 | 108.90 | 79.12 | 93.30 | 108.45 | 103.35 | 68.02 | 82.20 |
| BW27783 gly | 179.26 | 131.06 | 155.34 | 117.16 | 173.20 | 125.00 | 143.22 | 105.04 |
| BW27783 glyTre | 59.71 | 31.28 | 37.40 | 5.30 | 53.95 | 25.52 | 25.88 | -6.22 |
| BW27783 glu | 125.57 | 123.97 | 116.18 | 120.77 | 120.23 | 118.64 | 105.51 | 110.10 |
| BW27783 glyCas | 193.60 | 252.14 | 131.86 | 199.31 | 188.32 | 246.86 | 121.30 | 188.74 |
| BW27783 gluCas | 359.90 | 471.72 | 365.20 | 363.81 | 354.03 | 465.85 | 353.47 | 352.07 |

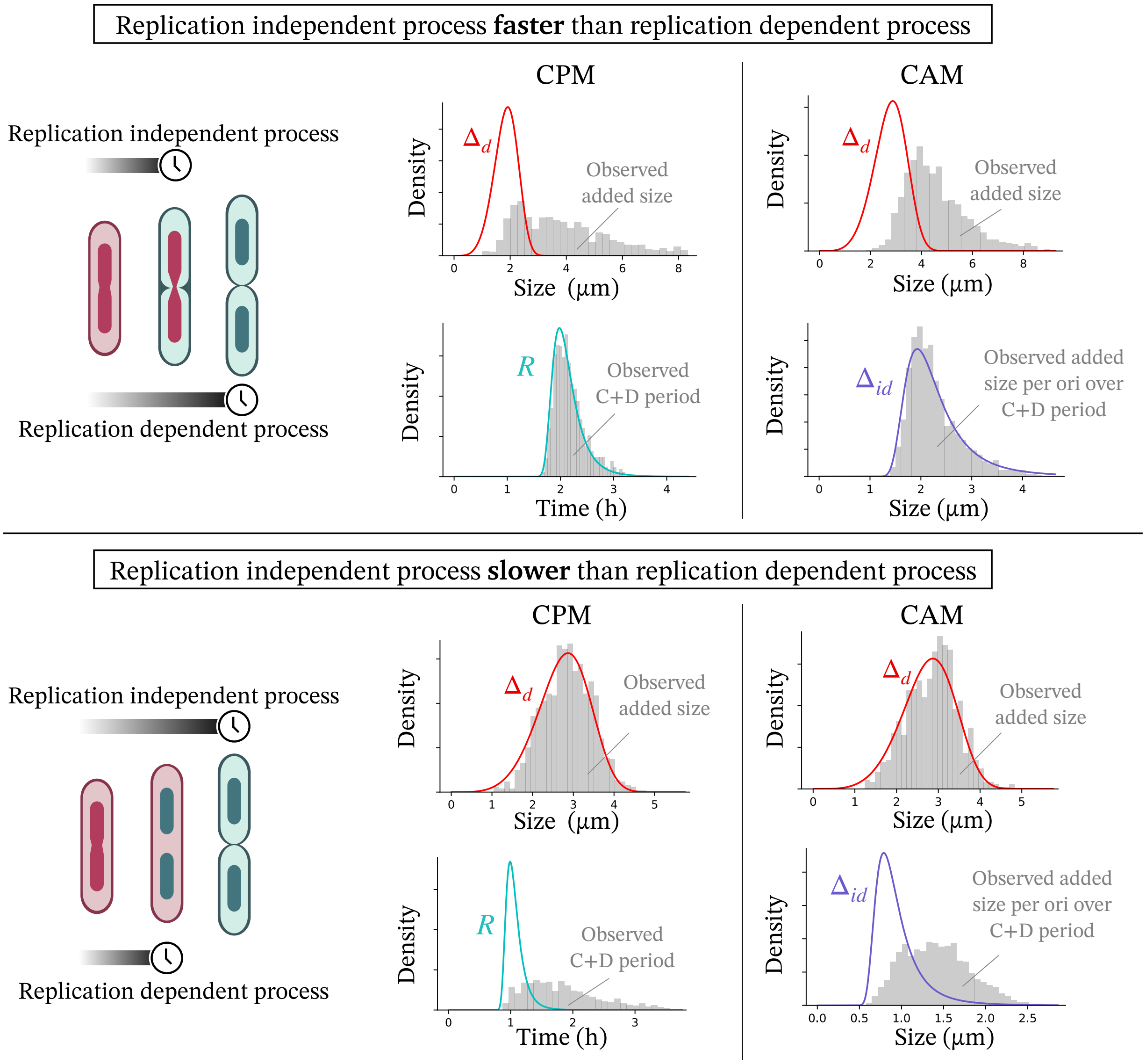


Figure S2: **Illustration of the discrepancy between observed and latent variables in the CPM and the CAM.** In both the CPM and the CAM, the division volume is determined as the maximum of two random variables: one associated with replication and the other with a division adder. As a consequence, observable quantities (e.g. added size between birth and division) do not always correspond directly to the latent variable (e.g. $\Delta_d$). This leads to a discrepancy between the true distributions of the latent variables $\Delta_d$, $R$, and $\Delta_{id}$ and the distributions of the observed added volume, the C+D period, and the added volume per origin over the C+D period, respectively. Such discrepancies arise particularly when one of the latent variables is sufficiently small that it is not taken into account by the maximum. To illustrate this, we consider two scenarios. In the first, replication is the main limiting factor for division (top), in which case the distributions of $R$ and $\Delta_{id}$ coincide with those of the C+D period and the size added per origin during the C+D period, respectively. In the second scenario (bottom), replication is not the main limiting factor for division, and the distribution of $\Delta_d$ coincides with that of the inter-division added size.

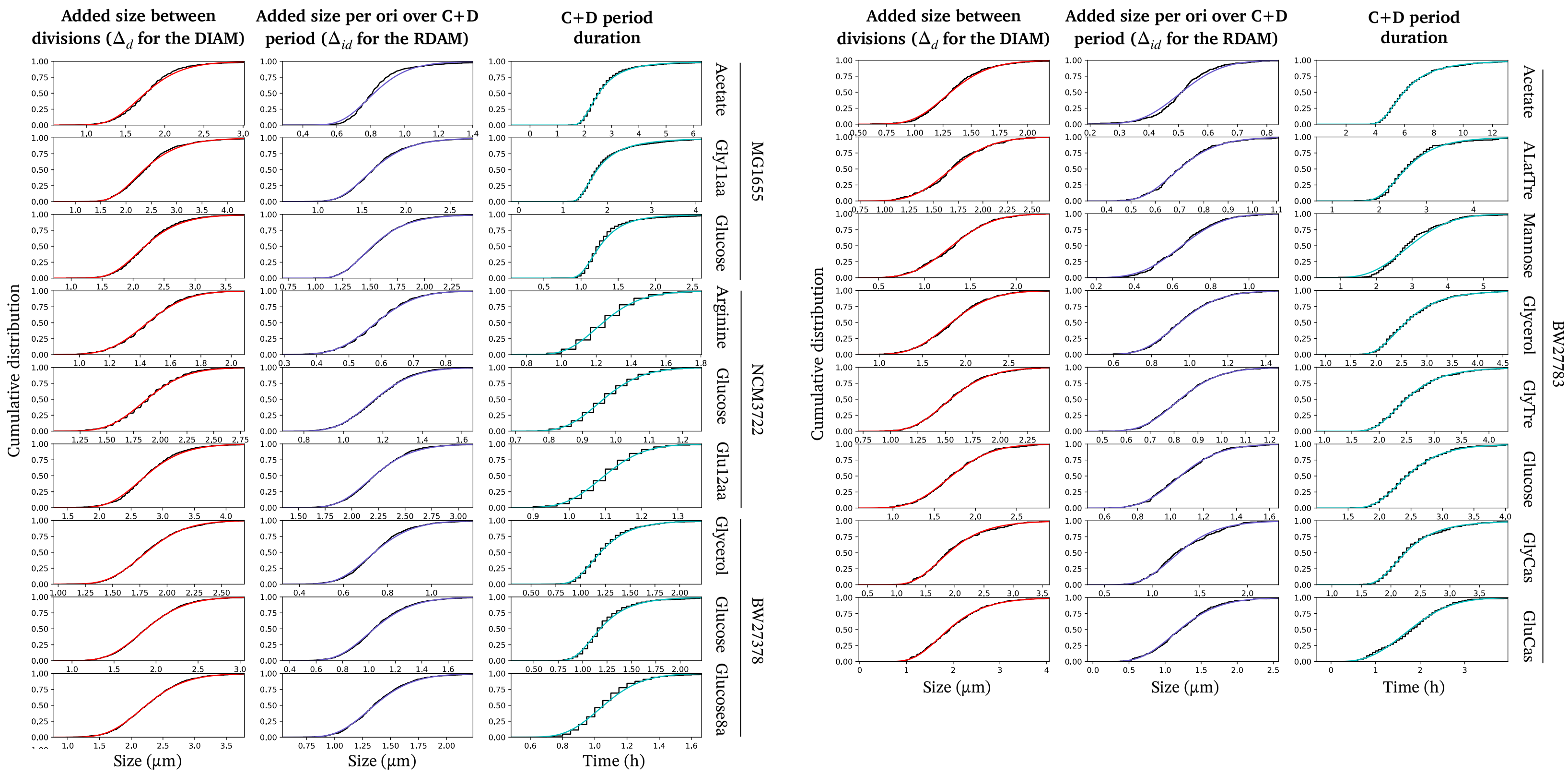


Figure S3: **Fitting Generalized Gamma distributions to inter-division added size, C+D period, and added size per origin over the C+D period.** In this work we assume that the i.i.d. random variables involved in the models follow Generalized Gamma distributions. The graphs show the empirical cumulative distribution functions (in black) and their corresponding fits by likelihood maximization to generalized Gamma distributions for three cell variables: size added between divisions (red), size added per origin during the C+D period, and the C+D period.

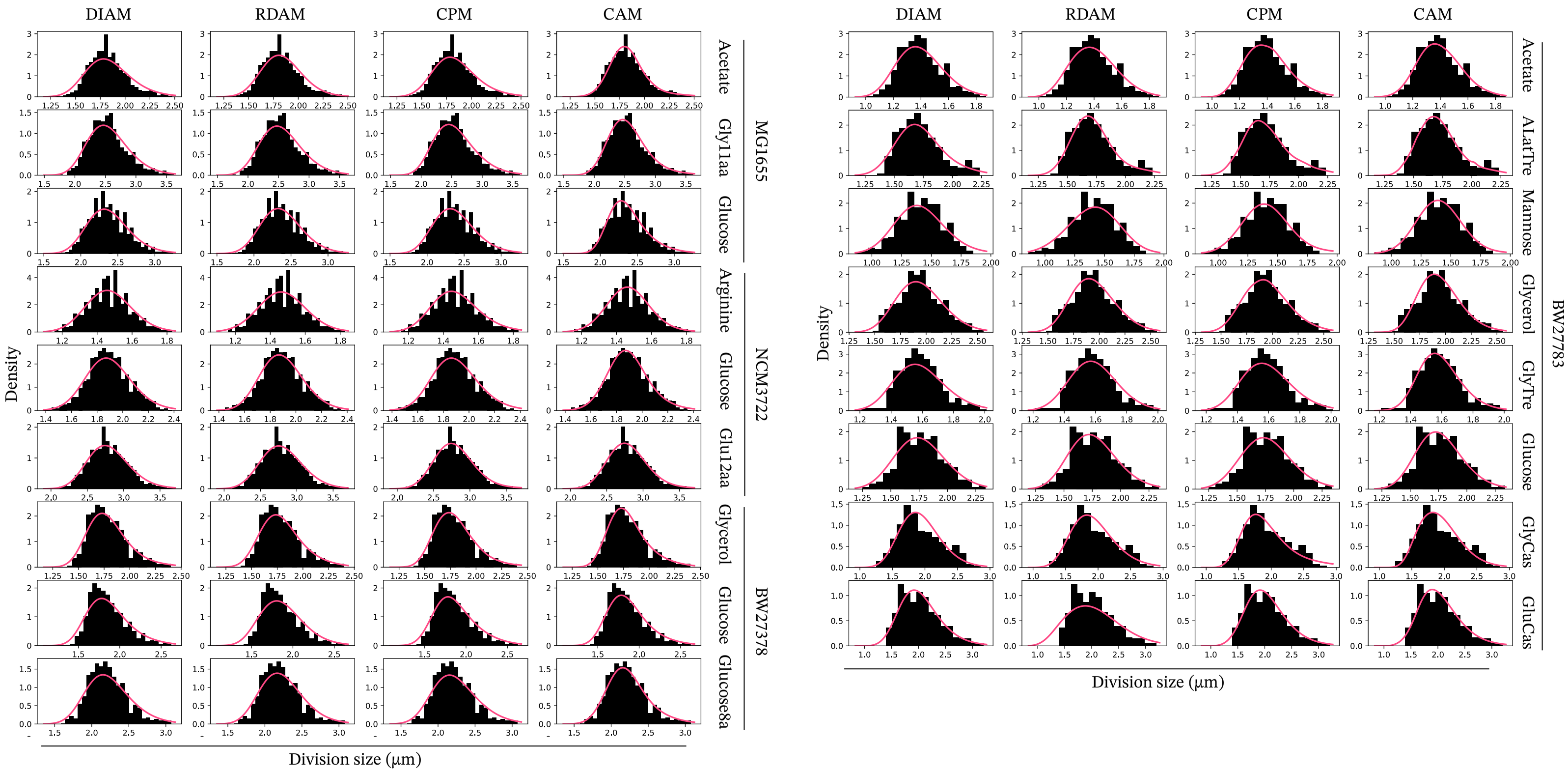


Figure S4: **Theoretical and empirical distributions of the size at division.** In black are shown the empirical distributions of the division size at division. The pink curve shows the theoretical distribution of division size according to the models computed from the experimental distribution of $(V_b^{(\bullet)}, \tilde{V}_i^{(\bullet)}, \Lambda^{(\bullet)})$.

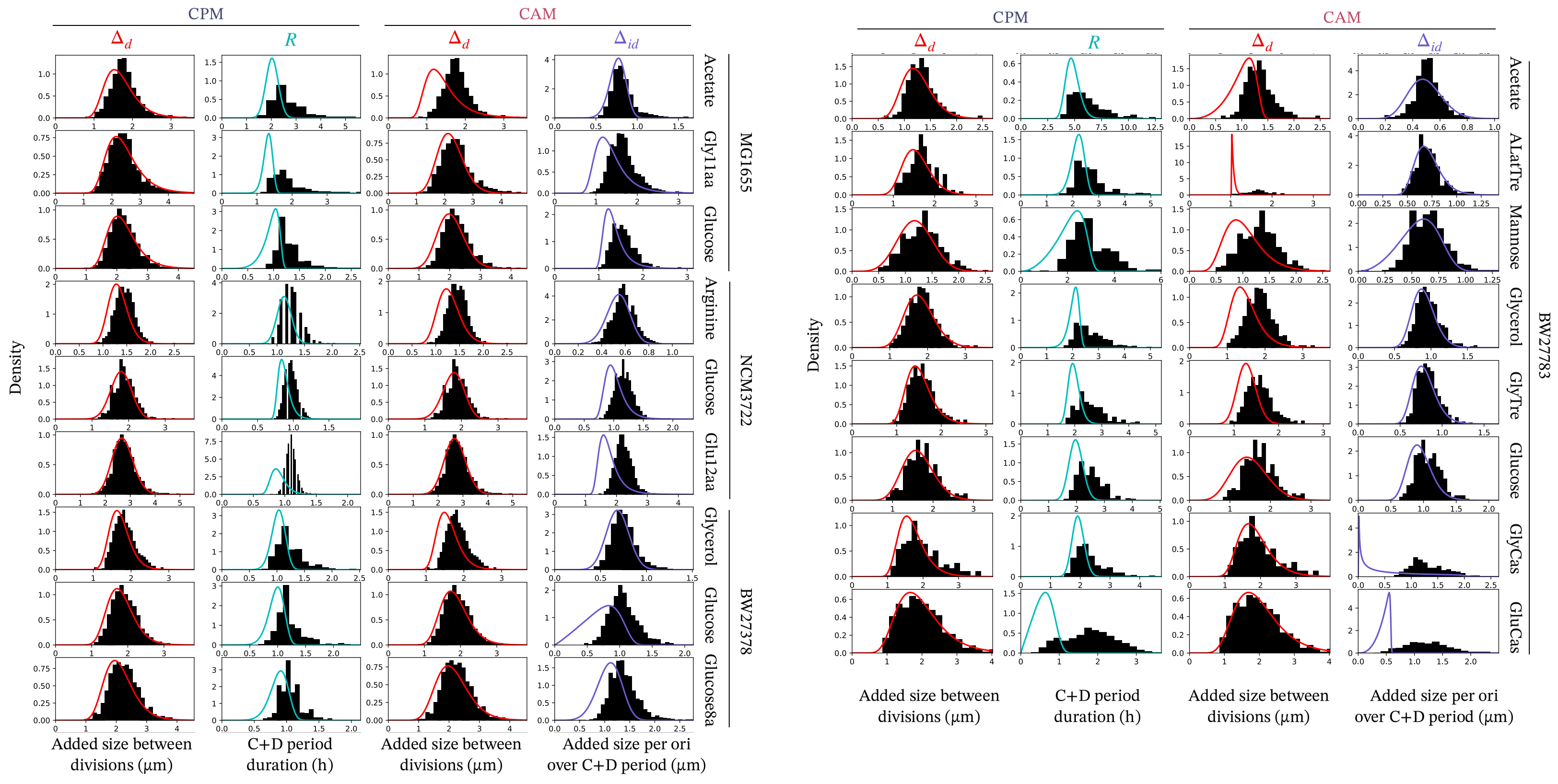


Figure S5: **Inferred distributions of the latent variables of the CPM and the CAM.** The Inferred distributions of the latent variables of the CPM and the CAM are shown by the colored curves. The black histograms shows the empirical distributions of the added size between divisions, the C+D period durations, and the added size per origin over the C+D period.

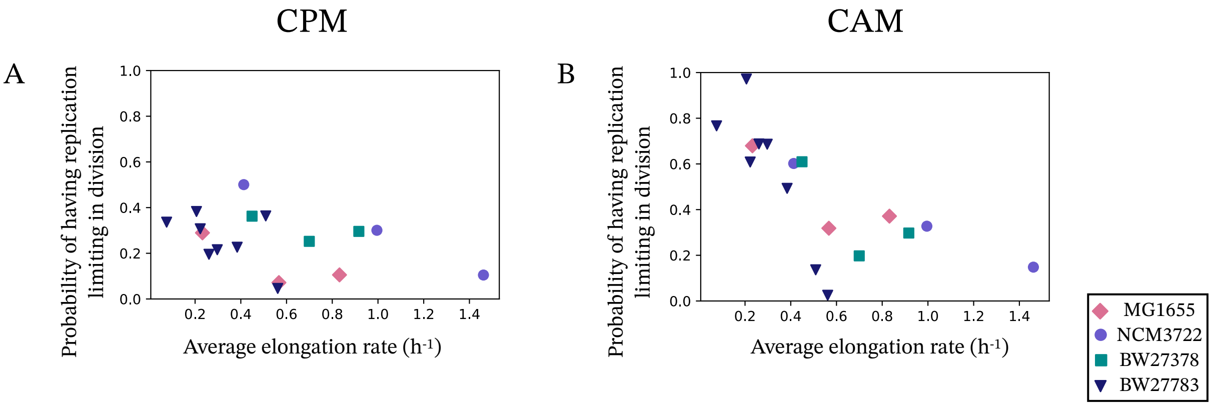


Figure S6: **Probability of having replication limiting in division in the CPM and CAM.** In A and B are displayed the probabilities of having the replication limiting in the division across growth conditions in the CPM and the CAM respectively. These probabilities have been obtained by computing in each dataset the average probability with the estimated distributions of $\Delta_d$, $\Delta_{id}$ and $R$, that the replication is limiting in the division process. As shown, the CAM predicts probabilities of having replication limiting in division close to one in slow growth conditions, which agrees with recent results (Tiruvadi et al. (2022), Kar et al. (2023))

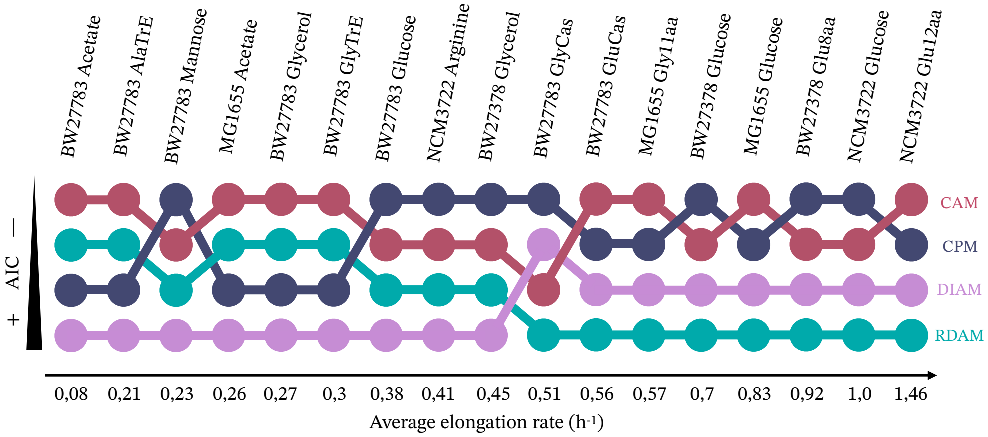


Figure S7: **Comparison of the performances of the models based on the AIC.** The AIC scores have been computed for each model (DIAM in pink, RDAM in cyan, CPM in dark blue and CAM in dark red) for multiple datasets of four strains. Here are shown the ranks of the models based on their AIC score.

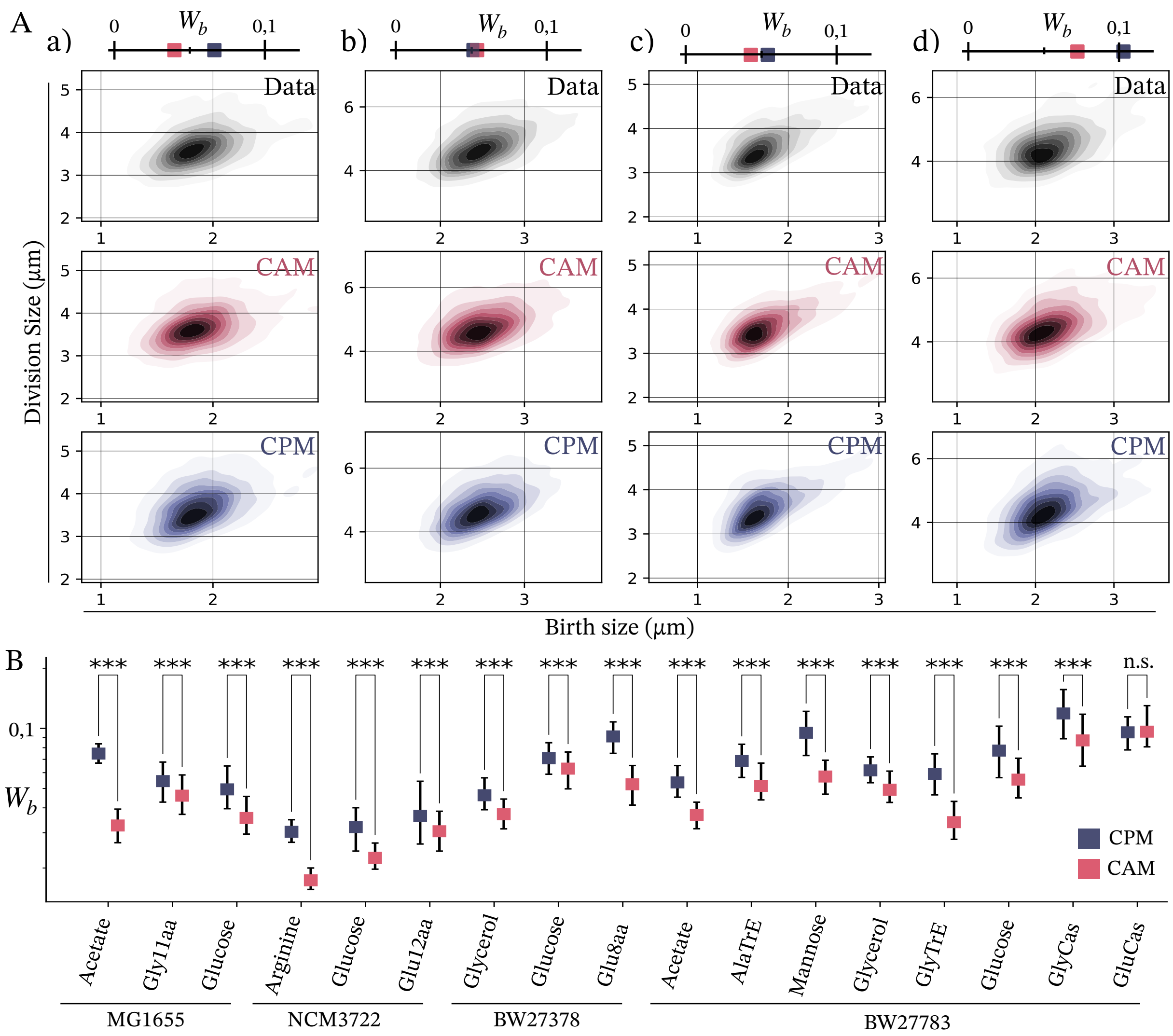


Figure S8: **Wasserstein distances between experimental and simulated joint distributions of division and birth sizes.** Wasserstein distances between experimental and simulated joint distributions of division and birth sizes for 100 simulations according to the CPM (blue) and CAM (red). Significant differences according to Wilcoxon-Mann Whitney tests are indicated: ***p-value $< 0.001$,n.s.= not significant (p-value $> 0.05$).

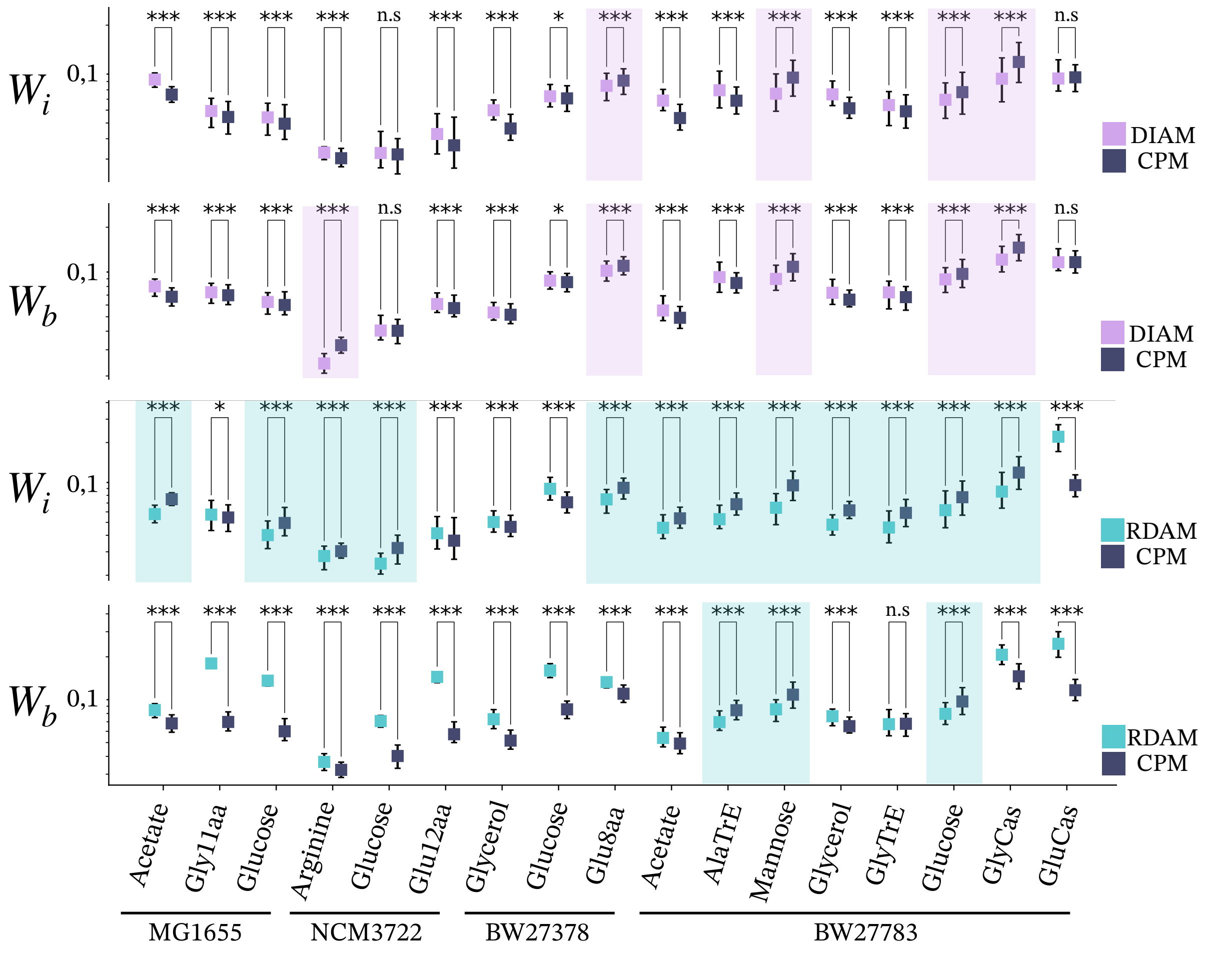


Figure S9: **Comparison of Wasserstein distances between DIAM/RDAM and CPM.** The Wasserstein distances between experimental and simulated joint distributions of initiation-division sizes and birth-division sizes have been computed for the RDAM and the DIAM. In comparison with the Wasserstein distances of the CPM, the DIAM and the RDAM shows significantly smaller Wasserstein distances in some datasets for both initiation-division sizes and birth-division sizes joint distributions. The shaded areas highlight the datasets where the CPM is outperformed by either DIAM or RDAM. $W_i$ denotes for the Wasserstein distance between experimental and simulated joint distribution of initiation-division sizes and $W_b$ for the Wasserstein distance between experimental and simulated joint distribution of birth-division sizes. Significant differences according to Wilcoxon-Mann Whitney tests are indicated: ***p-value $< 0.001$, **p-value $< 0.01$, *p-value $< 0.05$, n.s.= not significant

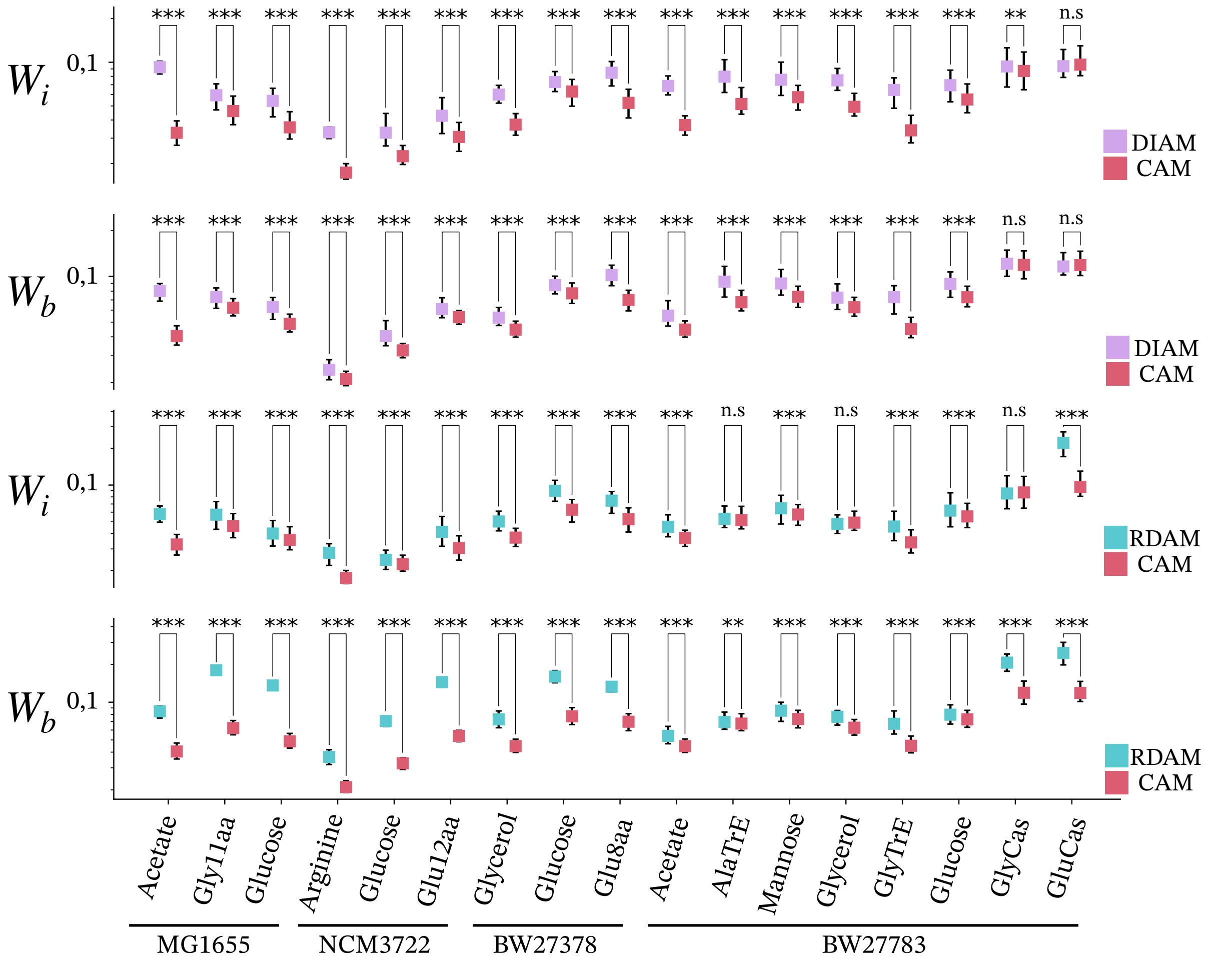


Figure S10: **Comparison of Wasserstein distances between DIAM/RDAM and CAM.** Similarly to figure S8, the Wasserstein distances between experimental and simulated joint distributions of initiation-division sizes and birth-division sizes have been computed for the RDAM and the DIAM. In comparison with the Wasserstein distances of the CAM, the DIAM and the RDAM do not show significantly smaller Wasserstein distances, and conversely, the CAM yields significantly smaller Wasserstein distances than the DIAM and RDAM in a strong majority of datasets. $W_i$ denotes for the Wasserstein distance between experimental and simulated joint distribution of initiation-division sizes and $W_b$ for the Wasserstein distance between experimental and simulated joint distribution of birth-division sizes. Significant differences according to Wilcoxon-Mann Whitney tests are indicated: ***p-value $< 0.001$, **p-value $< 0.01$, *p-value $< 0.05$, n.s.= not significant

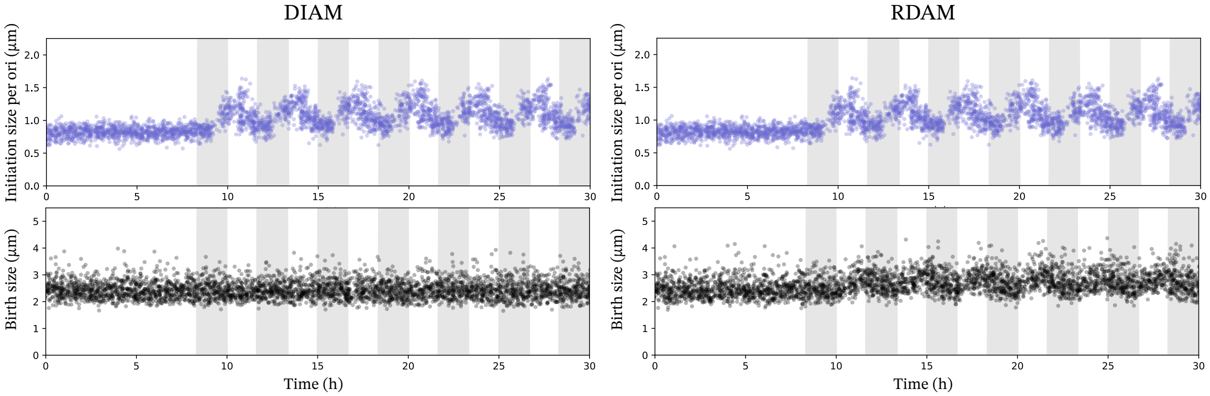


Figure S11: **Simulations of the DIAM and RDAM under oscillating distribution of $\Delta_i$.** As expected, oscillations of the initiation size does not affect the birth size in the DIAM, while it iduces oscillations of the birth size in the RDAM.
